# Temperature governs fitness effects of transgenes on plants non-monotonically: a global meta-analysis

**DOI:** 10.64898/2026.07.28.740297

**Authors:** Hairong Qian, Zhongyuan Wang, Jiangbo Xie

**Author notes:** Hairong Qian and Zhongyuan Wang contributed equally to this work and should be considered co-first authors. Correspondence author: Jiangbo Xie.

## Abstract

The global expansion of genetically modified plants supports modern food and ecological security^1^, yet whether the fitness cost—widely assumed to limit their spread after escape—actually operates in the wild has remained unclear, with empirical evidence remaining mixed^2–4^. Here, a global meta-analysis of 518 experiments across 48 species reveals that this ‘safety brake’ is real in the laboratory (fitness cost: ∼13%) but is largely attenuated in the world’s most intensively cultivated croplands. A phylogenetically robust, hump-shaped relationship between mean annual temperature and fitness effect is defined, with fitness benefits emerging specifically within 9.9–22.8°C. This ‘buffer window’ overlaps the productive croplands^5^, where the high-risk proportion (HRP; the area-weighted posterior probability of non-negative fitness effect) consistently exceeds that in natural vegetation and expands with global warming (cropland: +18.4% *versus* natural vegetation: +8.3% under SSP5-8.5). Notably, constitutive expression or dual-resistance (resist to both abiotic and biotic stressors) transgene can effectively compress the HRP below 30%, providing a quantitative baseline for management. These findings establish a framework for climate-smart deployment that balances agricultural productivity with ecological safety.

## INTRODUCTION

Genetically modified plants (GMPs) now cover >200 million hectares worldwide^1^, yet whether their elevating resistance carry the assumed ‘safety brake’—a fitness cost that should limit their spread upon escape^3,6^—remains unresolved after decades of debate^7,8^. This brake proves unreliable: it is frequently attenuated or even reversed into a fitness benefit in the field^8,9^, and is far from guaranteed even under controlled conditions^10,11^. The accumulated evidence thus supports cost, neutrality and benefit with near-equal weight^2^. Such tripartite ambiguity removes the directional consistency that deterministic risk models require, leaving regulatory tiering of ecological risk assessment (ERA) without a quantitative anchor^12^.

The conventional view holds a fitness cost arising from metabolic or pleiotropic effects^13,14^, which have been proven to exist at the molecular level^3^. Yet a comparable body of studies reports non-negative fitness effects^8^. This variability is not random; it stems from differences in genetic background^15^, resistance type^2,7^, generation^16^, gene number^17^, or expression system (e.g., inductive promoters can mask costs^18^). Recent syntheses have shown that costs are consistently expressed in wild genetic backgrounds but rarely in domesticated ones, pointing to artificial selection as a buffer^4^. Yet these insights into intrinsic mechanisms share a critical blind spot: the environment, an omission that has drawn increasing criticism^12^.

The environmental dependence of fitness effects was proposed over two decades ago^7^, with costs observed under controlled conditions but disappearing in the wild^19,20^. Despite frequent reports, the mechanistic basis of this buffering remains vague. Some attribute this to resource availability modulated by temperature, water, and nutrient supply in field conditions^7,8^; or to fluctuating conditions that benefit transgenic resistance in real fields^20–22^. Yet no predictive framework exists to map where and when the brake fails.

Consequently, most ERA models remain spatially implicit and deterministic^23^, unable to answer two simple but urgent questions: **(Q1)** is fitness cost an inherent constraint of GMPs, or a conditional outcome? and **(Q2)** when and where does it actually manifest?

Here we address this gap by assembling a global dataset of 1,256 observations covering 38 fitness indices across 48 species (Extended Data Fig. 1, Supplementary Table 1), drawn from 518 experiments in 224 studies (Fig. 1a, b). Fitness effects were derived by comparing high-resistance (HR) lines with low-resistance or wild type (LR) lines within the same genetic background under stress-free conditions^13^. A multi-stage analytical framework was applied to dissect the drivers of fitness effects across controlled and field conditions, including random forest, meta-regression, restricted cubic splines (RCS), and Bayesian modelling. The modulators considered encompassed plant genetic features, transgenic design and environmental context (Supplementary Table 2). Three climatic metrics were selected from 21 WorldClim variables^24^: annual mean temperature (MAT), annual mean precipitation (MAP), and precipitation of the driest month (PDM) (Extended Data Fig. 2, Supplementary Table 3–8). A parallel phylogenetic sensitivity analysis was performed to control for species relatedness (see Methods)^25^. Using the field-fitted models, the global HRP (high-risk proportion) in cropland/natural vegetation was projected, as a measure of ecological risk for escaped GMPs.

**Fig. 1.**
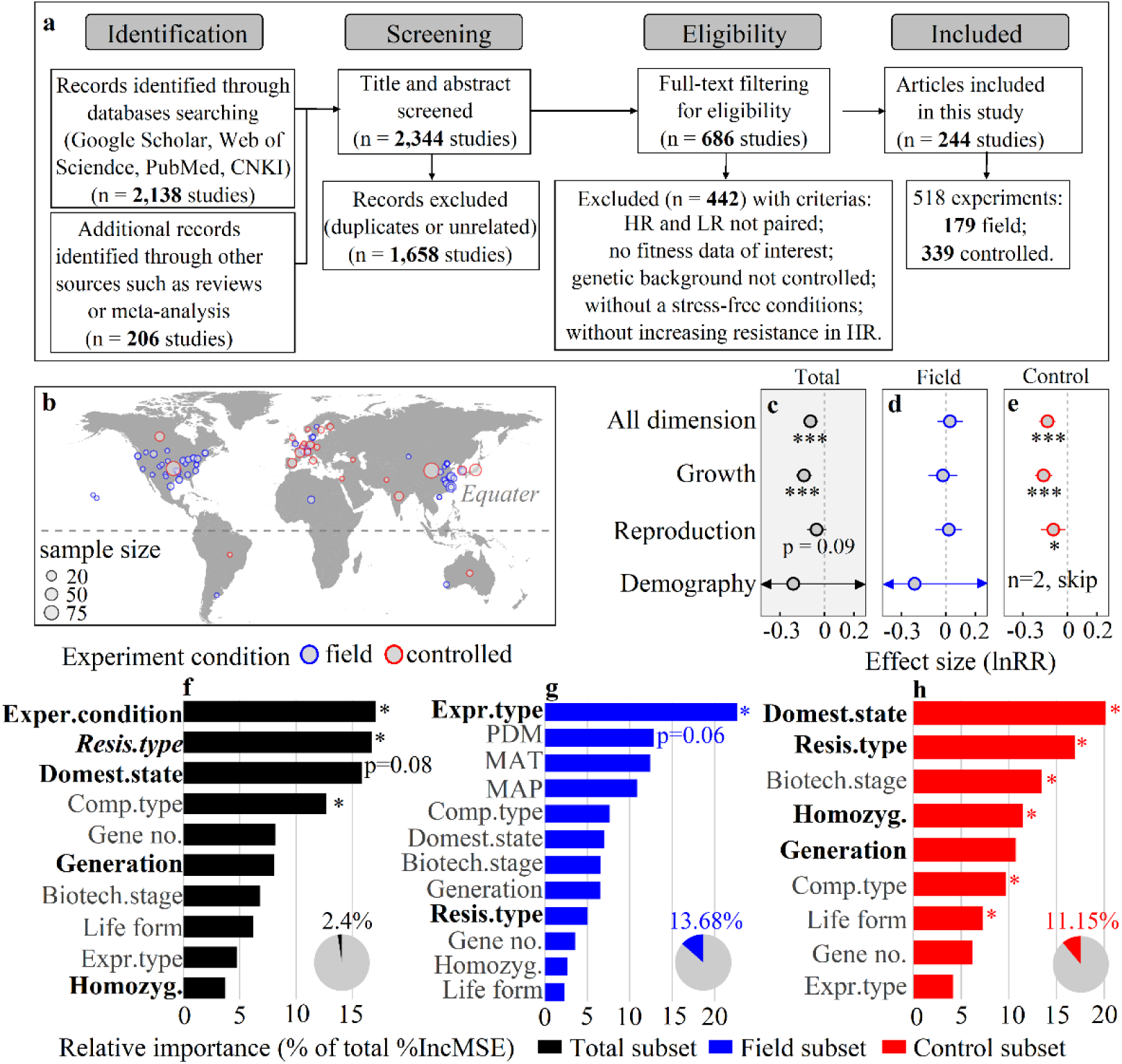
Overview of study selection, experimental distribution, effect-size estimation, and moderator importance. **(a)** PRISMA diagram of study selection. **(b)** Global distribution of 518 experiments in the total dataset. **(c-e)** effect sizes (non-phylogenetic model; phylogenetic correction yielded qualitatively identical results) for overall and for each dimension across total, field, and control datasets: *P < 0.05, **P < 0.01, ***P < 0.001. **(f-h)** Random-forest moderator importance rankings; asterisk denotes significant permutation test in random-forest. **Bold** = significant, ***italic*** = marginal in meta-regression. Full moderator names are provided in Supplementary Table 2.

*Three main findings emerge. First, a detectable fitness cost under controlled conditions is neutralized in the wild. Second, MAT regulates this effect in a unimodal manner, with fitness benefits concentrated in mid-latitude regions—the very band containing the world’s most intensively cultivated croplands. Third, future high-risk areas are consistently larger in cropland than in natural vegetation, and this disparity widens with global warming. Crucially, transgenic design features can compress the global HRP below 30%, providing a quantitative anchor for regulatory tiering*.

## RESULTS

### Estimation of fitness effect for transgenic resistance

Of the 1,256 observations, 55% showed non-negative effects (neutrality or benefits) and 45% showed costs (Supplementary Table 9). A multilevel meta-analytic model estimated the overall fitness effect to be -0.10 (95% CI: -0.14 to -0.05; Fig. 1c), largely driven by the controlled conditions (estimates = -0.14, 95% CI: -0.19 to -0.08; Fig. 1e, Supplementary Table 10). This negative effect was robust to imputation and quality filters (Extended Data Fig. 3, Supplementary Table 11) but was undetectable in the field subset (all CIs spanning zero; Fig. 1d). Phylogenetic correction produced qualitatively similar results (Extended Data Fig. 4a–c), with negligible phylogenetic signal (Supplementary Table 12, 13). This pattern was consistent across fitness dimensions, with a positive growth-reproduction correlation observed (Extended Data Fig. 4d, e).

### Drivers of fitness costs

In the random-forest model of total dataset (R² = 2.4%), experiment condition (field *vs.* control) was the dominant moderator (Fig. 1f), with a significant cost detected under controlled conditions but neutralized in the field (Fig. 2). Stratification by condition substantially improved model R², from 2.4% overall to 13.7% in the field and 11.2% under controlled conditions (Fig. 1g, h).

**Fig. 2.**
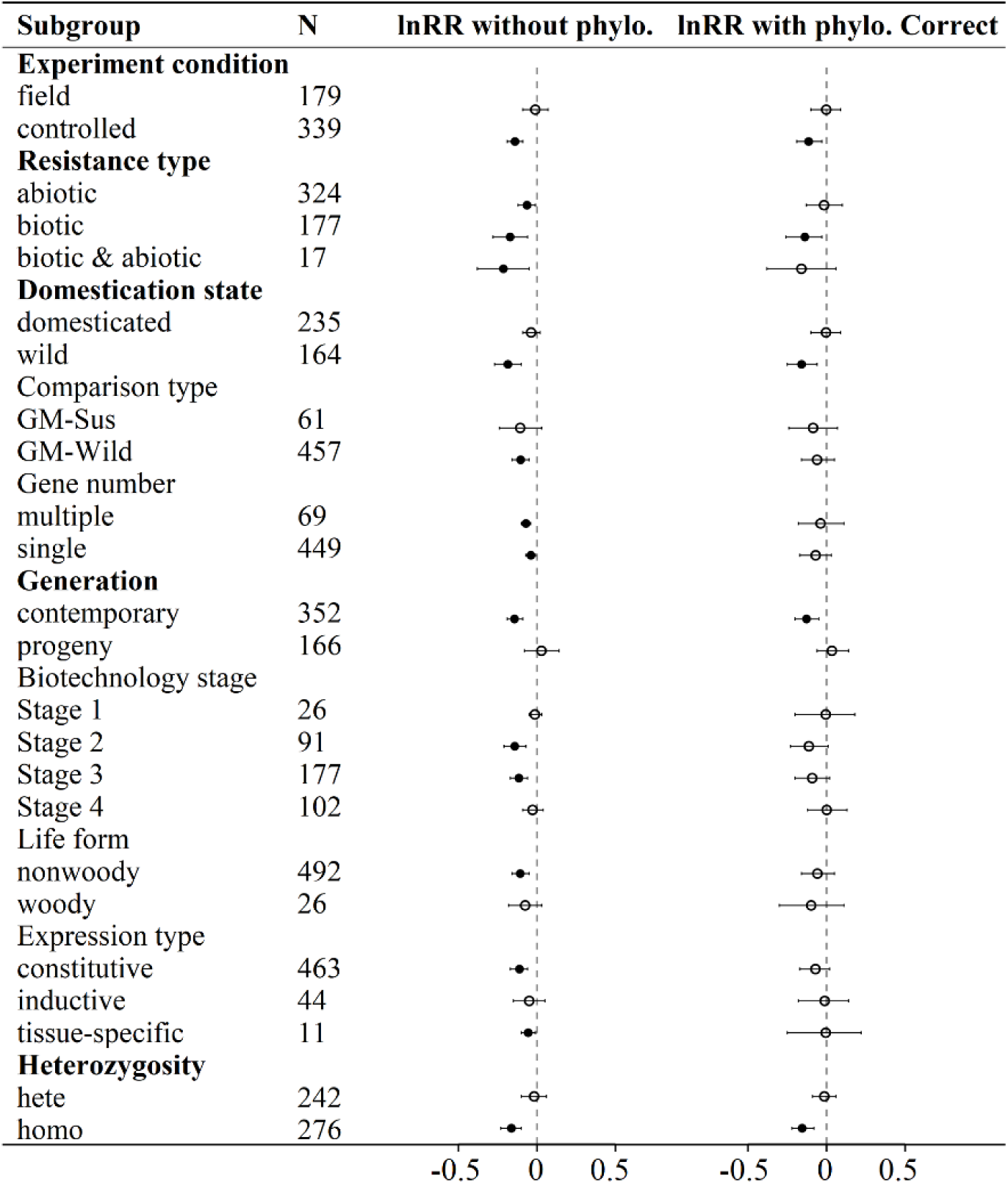
The effect of different levels of modulators on fitness effect in the total dataset. Significant subgroups in **bold**, with significant (p < 0.05, filled) levels and non-significant (open). Phylogenetic correction was applied only to the total dataset, as attempts on the field and control subsets failed due to weak phylogenetic signal (Supplementary Table 13).

Focusing on the field subset, multi-stage analysis yielded concordant findings: dual-resistance imposed a fitness cost (significant in random-forest and meta-analysis, Fig. 1g, Extended Data Figs. 5; Bayesian estimation = -0.21, Fig. 3e), whereas inductive expression systems conferred benefits (Bayesian estimate = 0.21; Fig. 3d). MAT, MAP and PDM ranked highly (Fig. 1g) but detected no linear trends (Extended Data Figs. 5). Instead, both RCS (β₂ = -0.07) and the Bayesian quadratic term (β₂ = -0.11) consistently identified a unimodal relationship between MAT and fitness effect (Fig. 3a, c; Supplementary Table 14). Positive predictions concentrated in the temperate zone (Fig. 3b), ranging from 9.9–22.8℃ (Fig. 3c). The Bayesian results after phylogenetic correction are highly consistent, with a negligible phylogenetic signal (σ_species = 0.01; Extended Data Fig. 6, Supplementary Table 15).

**Fig. 3.**
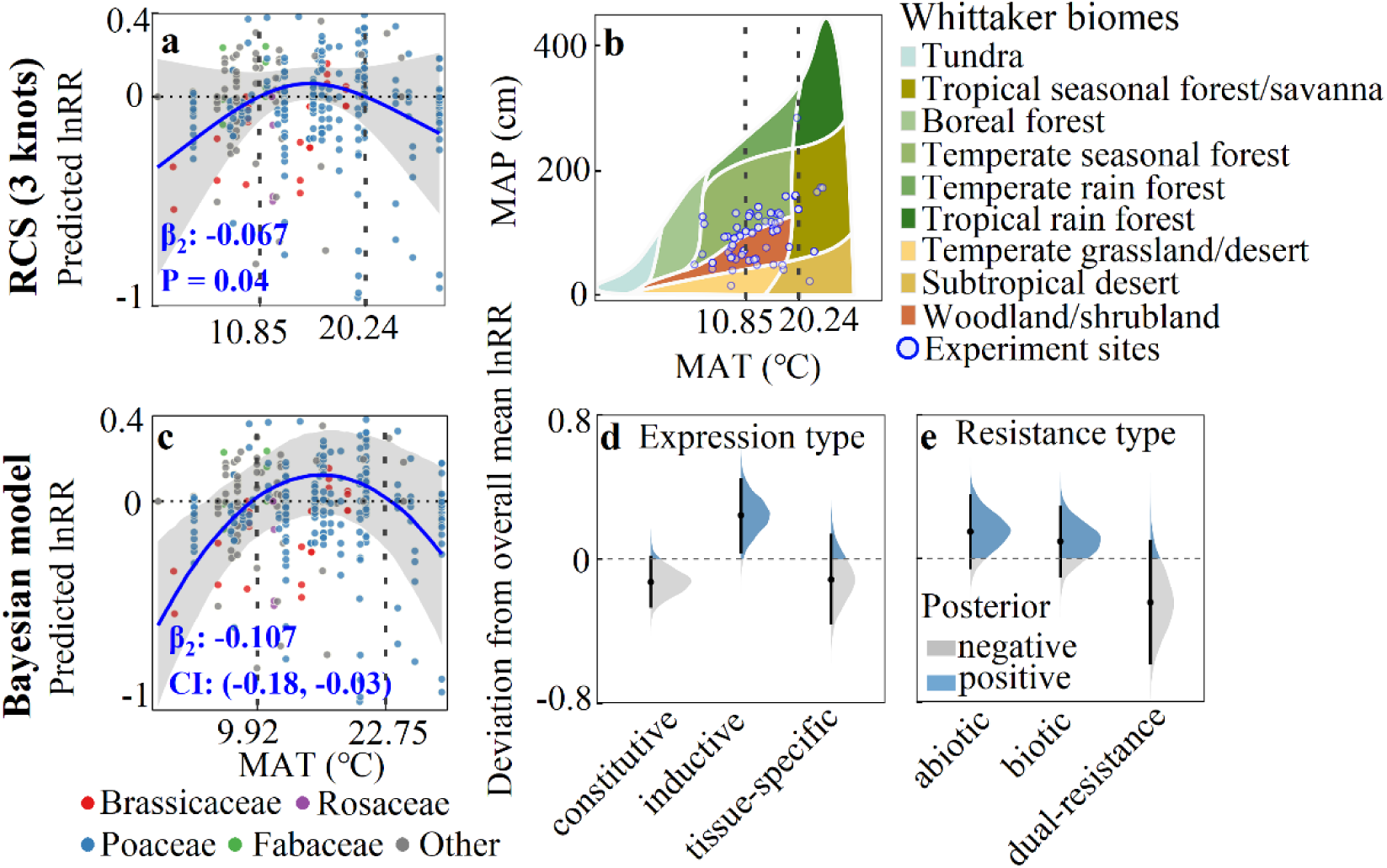
Nonlinear relationship between MAT and fitness effect (lnRR) in the field subset. **(a)** RCS shows a single-peak correlation (quadratic coefficient β₂ on standardized scale; P-value from FDR-corrected likelihood ratio test). **(b)** Global distribution of experiment sites with positive RCS predictions. **(c)** Bayesian model (506 field observations, with phylogenetic correction) confirms the single-peak relationship, with two categorical moderators accounted for: resistance type (**d**) and expression type (**e**). For categorical moderators, deviation from overall mean (sum-to-zero contrast) quantifies each level’s independent contribution; points ± bars = median ± 95% CI.

### Predictions of global ecological risk for escaped GMPs

Given the moderate explanatory power (∼53% of variance), the non-phylogenetic Bayesian model was used to predict global ecological risks for escaped GMPs under historical climate (1970–2000). Among the four transgenic designs (resistance type × expression type), GMPs with inductive abiotic resistance yielded the highest HRP, with >60% of vegetation area showing fitness neutral or benefits, constitutive expression or dual-resistance reduced HRP to below 30% (Fig. 4, Extended Data Fig. 7). Across all scenarios, cropland HRP exceeded that of natural vegetation (Extended Data Fig. 8).

**Fig. 4.**
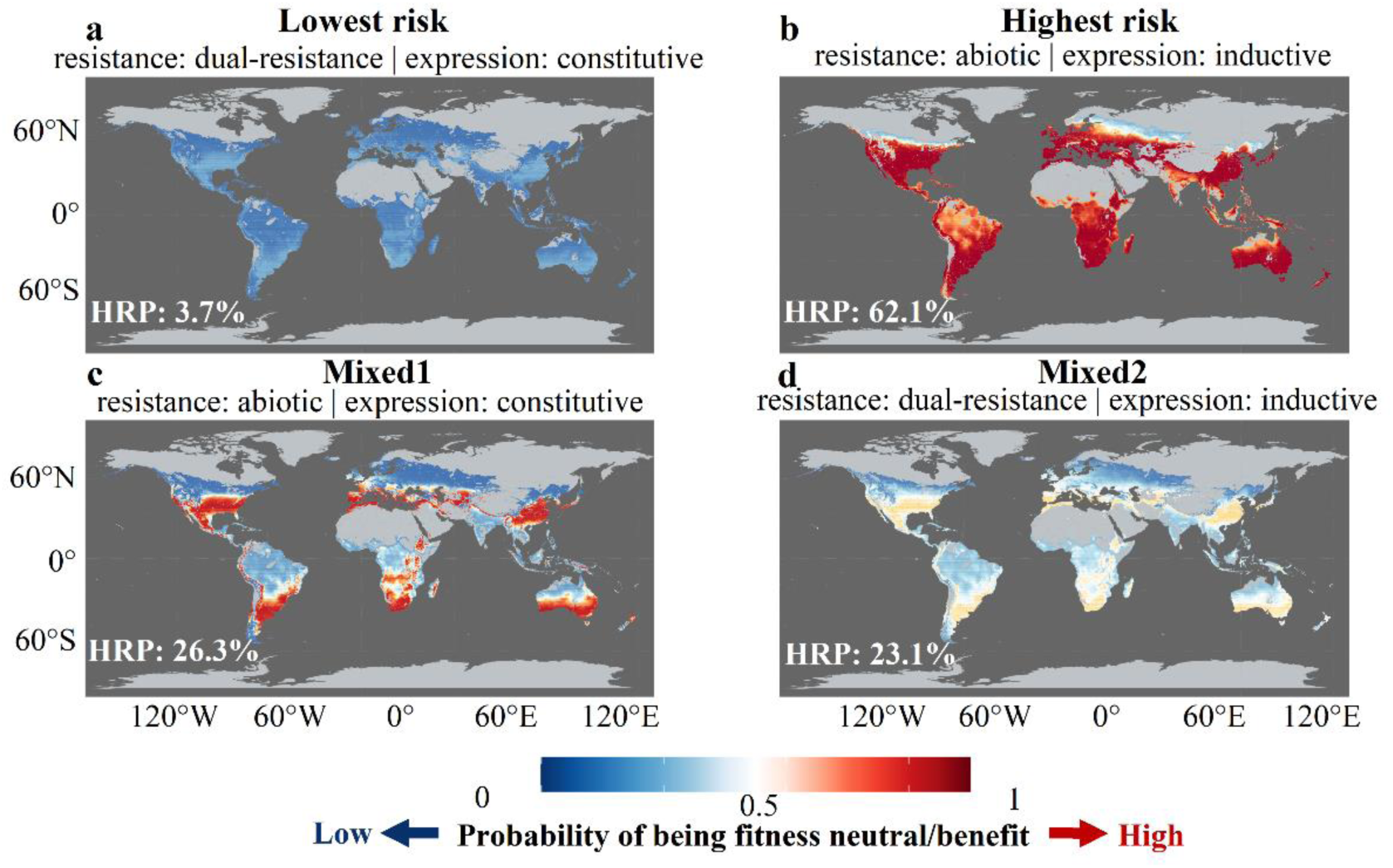
Constrained global risk maps. Colors represent the posterior probability that effect size (lnRR) ≥ 0: Blue shades (0–0.5) indicate low risk and red shades (0.5–1) indicate high risk. high-risk proportion (HRP) is the area-weighted posterior probability that effect size (lnRR) ≥ 0, measuring the area fraction with high ecological risk of escaped GMPs. Gray areas represent non-vegetated land, areas outside the training range (1.7–26.8 °C) or ocean. Four risk scenarios: **(a)** Lowest risk (dual-resistance + constitutive expression). **(b)** Highest risk (abiotic resistance + inductive expression). **(c)** Mixed1 (abiotic resistance + constitutive expression). **(d)** Mixed2 (dual-resistance + inductive expression). This constrained map provides higher confidence estimates for regions where the model has empirical support.

Future projections (2021–2100) across three Shared Socioeconomic Pathways (SSPs; SSP1-2.6: low emissions, SSP2-4.5: intermediate emissions, SSP5-8.5: high emissions) showed mixed changes in HRP for total vegetation (relative change: -1.3% to 7.3%) and natural vegetation (-3.9% to 8.3%), but cropland HRP consistently increased (0.7–18.4%) by the end of the century (Extended Data Fig. 9, Supplementary Table 16). Extrapolated predictions (temperatures outside the training range) showed no consistent trend and are presented for reference only (Extended Data Fig. 10). Consistent with historical estimates, HRP in cropland exceeded that of natural vegetation across all periods (Supplementary Tables 17, 18), and higher emission pathways amplified this disparity (Fig. 5).

**Fig. 5.**
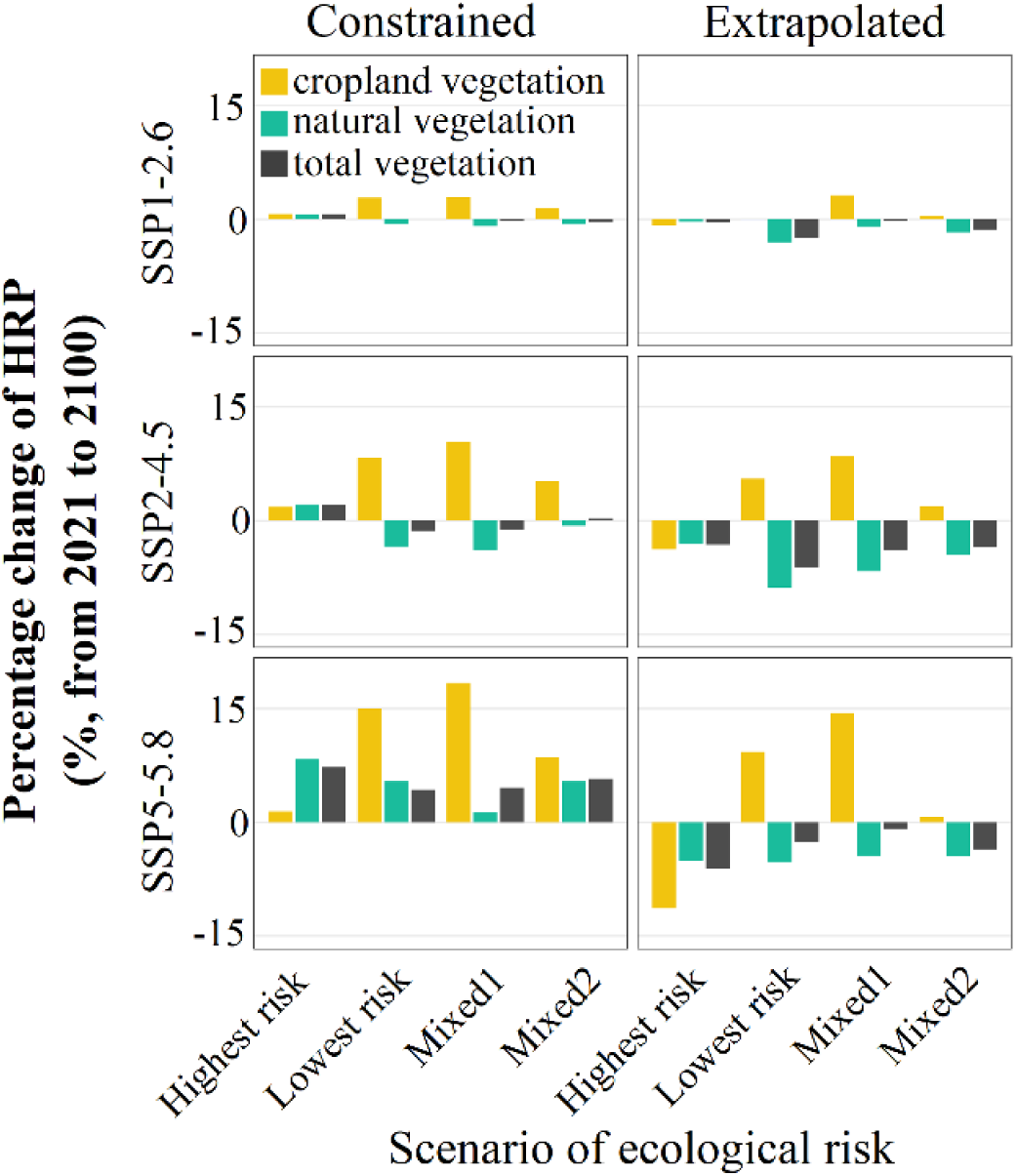
Percentage change of high-risk proportion (HRP) by 2100 relative to 2021 under three Shared Socioeconomic Pathways (SSP1-2.6, SSP2-4.5, SSP5-8.5). Predictions on constrained (within the temperature range of 1.7-26.8 °C) and extrapolated (full global range) models are shown separately. HRP represents the expected fraction of vegetated area at high ecological risk. Within each panel, four risk scenarios (defined by the combination of expression type × resistance type) are presented. As all pixel-level HRP values have extremely wide CIs globally, only the posterior medians are presented as bar charts.

## DISCUSSION

These findings resolve the long-standing ambiguity by demonstrating that cost, neutrality, and benefit are not fixed attributes of GMPs, but are environmentally contingent and modulated primarily by temperature. Globally, the fitness costs on GMPs are real (overall: ∼9%; controlled conditions: ∼13%), yet conditionally buffered in the field. This buffering is not uniform, as previously assumed^7,9,26^, but confined to a specific buffer window (MAT: 9.9–22.8 °C), within which fitness benefits can reach up to 13% and outside of which the intrinsic cost re-emerges. The prominent role of temperature in this non-monotonic pattern likely arises from two convergent mechanisms: at the molecular level, temperature mediates phytohormone signaling between the growth and defense^27,28^; at the resource level, mid-latitude temperature defines the thermal window of highest productivity^29^, where resource availability is maximal^30^, providing physiological slack to offset defense-associated costs (Growth-Differentiation Balance Hypothesis)^31,32^. This dual role may explain why temperature was the sole factor (among 21 climatic variables) exhibiting a clear relationship with fitness effects.

The key is the ascending segment (10.0–16.3 °C) of the curve: under accelerating warming^33,34^, each 1 °C warming towards the optimum (16.5 °C) translates into an ∼11% fitness gain (exp (0.11) ≈ 1.116). This sensitivity implies that progressive warming will erode the ‘safety brake’ of fitness costs in regions currently within this segment, which spatially overlap with global croplands^35^. Indeed, the projections confirm that cropland HRP consistently exceeds that of natural vegetation across all future scenarios, and this disparity increases under the high emission scenario (e.g., SSP5-8.5). This pattern is consistent with a buffering effect amplified by agricultural inputs (e.g., irrigation and fertilization)^7^, and with the warmer microclimate of croplands relative to surrounding natural vegetation^36^—escaped GMPs entering cropland may effectively encounter an artificial ‘ascending segment of the curve’ where costs are masked. These findings call for strengthened regulatory oversight of agricultural systems, where domesticated genetic backgrounds already exhibit fitness neutrality and transgene persistence is more probable (as is the case in this study, Fig. 2)^4,37^. Notably, even within the buffered window, the HRP can be halved by adopting constitutive expression or dual-resistance designs, which can increase the metabolic burden by prolonging the period of transgenic expression^7,38^. Their combination even reduces the ecological risks to negligible level (HRP < 5%). These findings therefore propose that ERA frameworks prioritize constitutive or broad-spectrum constructs, particularly in mid-latitude regions where climatic vulnerability and cropping intensity spatially converge^39^.

### Caveats and conclusions

Several frontiers remain for future research rather than constituting fatal flaws. First, the near-absence of demographic fitness measures (only 25 out of 518) severely constrains inference at population-level, calling for standardized multi-generational field experiments^40,41^. Second, the low explanatory power of individual models (random forest R²: 2–14%; meta-analysis I² > 99%), reflecting the inherent complexity of ecological data and the necessity of considering other environmental factors (e.g., soil, microbial environment) in future studies^42,43^. Despite this, the convergent directional signals across all analytical approaches substantiate the main conclusions^44^. Third, although an ISIMIP3b-constrained CMIP6 ensemble was used in the future projections, scenario-specific risk distributions retain structural uncertainty^45^; refined projections are needed as climate models evolve^5^.

Globally, transgene fitness costs are real but conditional in the field—regulated by MAT non-monotonically. The resulting risk geography (9.9–22.8°C) centers on the world’s most productive croplands^35^, where agricultural benefits and ecological risks after escape spatially coincide. As warming elevates the escape risk of GMPs^46^, croplands will increasingly become both the source of transgene escape and the environment most conducive to its persistence. These results advocate for a tiered, climate-zone-specific ERA framework. This coincidence is not an argument against transgenic technology, but a mandate for climate-smart deployment and spatially targeted monitoring.

## Supporting information

The Extended Data Figures

The Supplementary Tables

## Author Contributions

Jiangbo Xie designed the research; Hairong Qian and Zhongyuan Wang extracted data and wrote the manuscript; Jiangbo Xie provided professional advice. All authors contributed substantially to revisions.

## Conflict of interest statement

There is no conflict of interest to declare.

## Acknowledgements

This work was supported by National Natural Science Foundation of China, Grant/Award Numbers: 42330503, 32371662.

## Methods

### Data collection

We followed PRISMA guidelines^1,2^ to build a dataset on HR and LR fitness in stress-free conditions (Fig. 1a). We searched Google Scholar (https://scholar.google.com), Web of Science (https://clarivate.com/webofsciencegroup/), and PubMed (https://pubmed.ncbi.nlm.nih.gov/) until 1 January 2026 using the following search string [TS = (transgenic* OR genome edit*) AND (fitness* OR growth OR *agronomic traits* OR yield*) AND (resistance* OR resistant*)], as most relevant studies were not specifically designed to test fitness effects. Manual checks of key reviews yielded 206 additional studies. After deduplication with Zotero (www.zotero.org), we screened titles/abstracts and assessed full texts against three criteria: (1) inclusion of at least one control (LR, susceptible, or wild type) and one treatment group (HR with enhanced resistance); (2) reporting of ≥1 of 38 fitness indices (Supplementary Table 1); (3) controlled genetic background between HR and LR. Experiments with competitive environments were excluded due to unclear effects on cost estimation. Excluded literature is listed with reasons (Supplementary Table 19).

In total, we obtained 244 studies with paired observations across 48 species (28 domesticated, 20 wild; Extended Data Fig. 1, Supplementary Table 20). For papers involving multiple target genes, backgrounds, generations, or environments, each unique combination was treated as an independent experiment (518 total). We extracted mean, sample sizes (*n*), and standard deviations (SD) from HR and LR; for multiple transgenic lines, the mean effect was calculated. Data from graphs were extracted using Plot Digitizer (http://plotdigitizer.sourceforge.net/). We recorded 13 moderators across three categories: (1) **genetic features** – domestication state, life form, gene number, and homozygosity; (2) **transgenic design** – resistance type, comparison type, generation, and expression type. The historical progression of plant biotechnology was categorized into four stages (1980–1995, 1996–2005, 2006–2014, 2015–present) to account for technical evolution^3,4^; (3) **environmental context** – controlled vs. field conditions. For field sites, 21 climate variables were extracted from WorldClim v2.1 (Supplementary Table 3)^5^, with three (PDM, MAT, MAP) retained after screening (Extended Data Fig. 2).

### Imputation procedure and sensitivity analyses

To maximize data coverage, missing information was imputed rather than excluded. Missing sample sizes were conservatively set to *n* = 3^6^. For missing measurement data: if representative photos were available, we used ImageJ (https://imagej.net/ij/) to estimate traits (e.g., leaf area) with *n* = 3; if only descriptive comparisons were reported, we assigned effect sizes of -0.2 (V_HR_ < V_LR_), 0.2 (V_HR_ > V_LR_), or 0 (no difference). Correlation-matrix reports were similarly coded (0.2 for positive, -0.2 for negative correlations). Variance for imputed effect sizes was estimated from the median variance of original data (more robust to outliers)^7^. This approach (assigning ±0.2 or 0) was chosen because we focused on direction rather than magnitude^8^, and these values fall within the main distribution of original effect sizes^9^. In total, 217 of 1,256 observations were imputed. Missing SDs were converted from standard errors, coefficients of variation, P values, t values, or 95% CIs^10^. Missing climate data were extracted from WorldClim based on reported or estimated (e.g., city, region or administrative boundary) coordinates^5^. For homozygosity: <5 backcrosses = heterozygous; ≥5 backcrosses (≥72.8% homozygosity) = homozygous^11^; unspecified hybrids were conservatively treated as heterozygous.

Sensitivity analyses (Supplementary Table 11) confirmed robustness: the original analysis (n=1039) gave lnRR=-0.13 (95% CI: -0.19 to -0.06), while imputed data gave -0.10 (-0.14 to -0.05). Best/worst-case scenarios and variance inflation (2×, 5×, 10×) all yielded qualitatively identical results^12^. The field subset remained non-significant throughout. The effect-size distribution was unimodal and centered near zero, with a slightly longer left tail (Extended Data Fig. 3), confirming that imputation did not distort the underlying data structure^13^.

### Effect size calculation and overall fitness effects

We estimated the log-response ratio (lnRR) as the effect size^14^, applying a second-order Taylor series expansion (Delta method) to correct for small-sample bias (Eq. 1–4)^9^. Negative lnRR indicates fitness cost; positive indicates benefit.

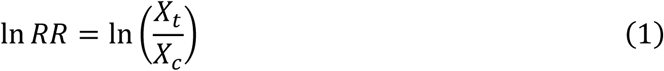

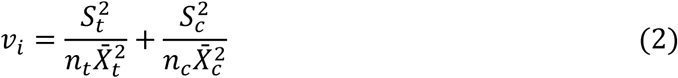

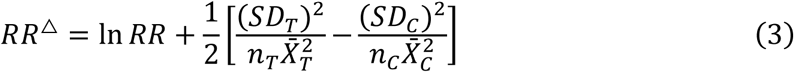

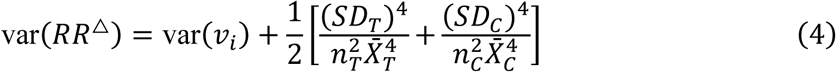

where X_T_ and X_C_ are mean fitness components of HR and LR, respectively.

To test whether enhanced resistance incurs fitness costs, we fitted a multilevel meta-analytic model (*M1*) using rma.mv in R (*metafor* package v.4.4.3)^15^, with random intercepts for Study ID, Control ID nested within Study, and Experiment ID:

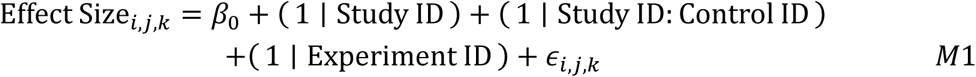

Models were fitted using the restricted maximum likelihood (REML) estimation. Heterogeneity was assessed with *Q*_t_ and *I*² (Supplementary Table 10)^16,17^.

To examine dimension-specific costs, we extended *M1* by adding (∼1|Obser_ID) as a random effect for growth (*M2a*), reproduction (*M2b*), and demography (*M2c*) separately:

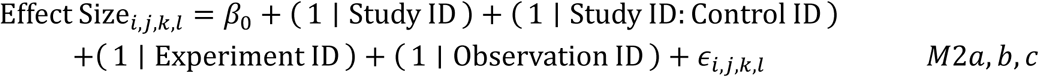

For experiments reporting both growth and reproduction (n = 224), we tested their proportional consistency via inverse-variance weighted regression, assessing whether the slope differed from 1 (Z-test)^15,18^.

### Drivers of variation in fitness effects

To disentangle the contributions of genetic features, study designs, and environmental contexts, we employed a three-tier analytical strategy. First, we used random forest (*randomForest* package v4.7-1.2, 1,000 trees) to rank moderator importance via %IncMSE, normalized to sum to 100%^19,20^.

Second, we tested linear moderating effects via multilevel meta-regressions (same structure as *M1*) with cluster-robust variance (CR2)^21,22^. This complementary approach leverages the exploratory power of random forest to identify influential predictors while the meta-regression provides formal statistical inference on their moderating effects^15,19^. For continuous climate variables (MAT, MAP, PDM; field subset only), we examined nonlinearity using restricted cubic splines (three knots at 10th, 50th, 90th percentiles), comparing linear vs. spline models via likelihood ratio tests^23,24^.

Third, for global projections, we built a Bayesian multilevel meta-regression (*brms* package v2.23.0, interfacing with Stan, *M3*) on the field subset^25,26^. The model included a quadratic term for MAT (BIO1), plus resistance type and expression type as fixed effects (sum-to-zero contrasts, appropriate when no factor level serves as a natural reference category)^27–29^, with weakly regularizing priors^30^: *β_k_* ∼ *N*(0,1), *σ* ∼ Exponential(5). The lnRR included sampling variance as a known measurement error. Convergence was confirmed (*R̂* = 1, Bulk-ESS and Tail-ESS > 1,000 for all parameters; Supplementary Table 14, 15)^31,32^.

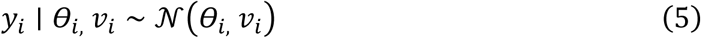

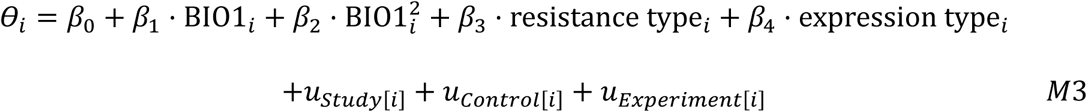

### Phylogenetic correction

We constructed a species-level phylogeny using R packages plantlist and V.phylomaker (Scenario 3for missing genera/species)^33,34^, and computed a Brownian-motion covariance matrix^35^. This matrix was converted to a correlation matrix and, where necessary, made positive definite using nearPD^36^. Phylogenetic signal in weighted mean effect sizes was weak and inconsistent (total: K = 0.278, p = 0.018; λ = 0.007, p = 0.969; field: K = 0.151, p = 0.383; λ = 0.378, p = 0.511; Supplementary Table 13)^37,38^. Including the phylogenetic covariance matrix as a random effect yielded negligible species-level variance (σ_species ≈ 0.01; 95% CrI: 0.00–0.03) with unchanged fixed estimates (Supplementary Table 15). Given the negligible phylogenetic signal and convergence issues in subsets with sparse species representation (and the random-forest algorithms do not accommodate structured random effects)^19,20^, we present the non-phylogenetic models in the main text, with full phylogenetic re-analyses provided in the Supplementary Information. The simpler Bayesian multilevel meta-regression (*M3*, without phylogenic correction) was applied in the main conclusions^39^ and generated posterior predictions (four chains, 4,000 iterations) for global risk mapping (see below).

### Global risk maps

We generated global risk maps from*M3*, using MAT (BIO1, 10-arc-min) from WorldClim v2.1^5^. Negative lnRR indicates a fitness cost (higher risk); positive indicates benefit (lower risk). Four transgenic scenarios were defined: (1) lowest risk (biotic & abiotic resistance + constitutive expression), (2) highest risk (abiotic resistance + inductive expression), (3) mixed1 (abiotic + constitutive), and (4) mixed2 (biotic & abiotic + inductive).

For each scenario, we predicted the cell-wise posterior probability of lnRR ≥ 0 for vegetated pixels (MODIS MCD12Q1 v6.1 mask; natural vegetation and cropland)^40^, under two temperature extents: (i) **conservative** – clamped to the training range (1.7–26.8 °C); and (ii) **extrapolated** – full global temperature range. Maps were reprojected to EPSG:4326 (Fig. 4, Extended Data Fig. 7). High Risk Proportion (HRP) –the area fraction with predicted lnRR ≥ 0 – was computed as the area-weighted average of cell probabilities (Eq. 6)^41^. This estimate corresponds to the posterior mean of the risk area proportion and is more robust than a median based on binary thresholding^42^:

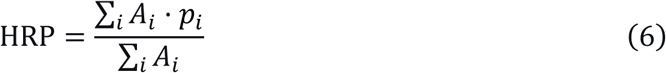

The wide 95 % CIs for HRP (e.g., 0–73.5 % for cropland) reflect extrapolation uncertainty and threshold sensitivity near lnRR = 0^43^. Despite this, the posterior median HRP consistently ranked the four scenarios (highest risk > mixed1 ≈ mixed2 > lowest risk) across both extents, with no overlap between extreme scenarios across all 500 posterior draws (full samples provided in Source Data).

Future projections used the same *M3* with multi-model mean (MMM) of BIO1 from five CMIP6 models (EC-Earth3-Veg, IPSL-CM6A-LR, MPI-ESM1-2-HR, MRI-ESM2-0, UKESM1-0-LL) following ISIMIP3b protocol (Supplementary Table 21)^44,45^. MMM data (2.5 arc-min) were bilinearly resampled to 10 arc-min to match the historical baseline. Predictions were generated for SSP1-2.6, SSP2-4.5 and SSP5-8.5 (warming: ∼0.46°C, 1.51°C, 3.47°C by 2100 relative to 2021; IPCC AR6)^46^, across four periods (2021–2040, 2041–2060, 2061–2080, 2081–2100), under both conservative and extrapolated extents (Supplementary Table 17, 18).

### Publication bias

Publication bias may arise because studies reporting negative fitness effects are more likely to be published. To mitigate this, diverse objectives (e.g., GMP performance trials) and those qualitatively reporting non-significant differences but lacking extractable data, for which effect sizes were conservatively estimated and sensitivity-tested.

We further assessed publication bias via three complementary approaches. Egger’s regression^47^ was significant for the total dataset (p = 0.006), but trim-and-fill analysis estimated zero missing studies (k₀ = 0), indicating that the negative overall effect is robust to bias^48^. Rosenberg’s fail-safe N (703) further confirmed that an implausibly large number of null studies would be required to overturn the overall negative effect^49^. Full results are provided in Supplementary Tables 22–23.

## Data availability

The source data underlying Figs. 1–5, Extended Data Figs. 1–10 and Supplementary Tables 5–18 and 22–23 are available via Figshare at. The climate data is available from the WorldClim database (worldclim.org).

## Code availability

The R scripts needed to reproduce the results are available via the Figshare repository at.

## Notes

### Competing Interest Statement

The authors have declared no competing interest.

