## Supplementary material for "Temperature governs fitness effects of transgenes on plants non-monotonically: a global meta-analysis": The Extended Data Figures

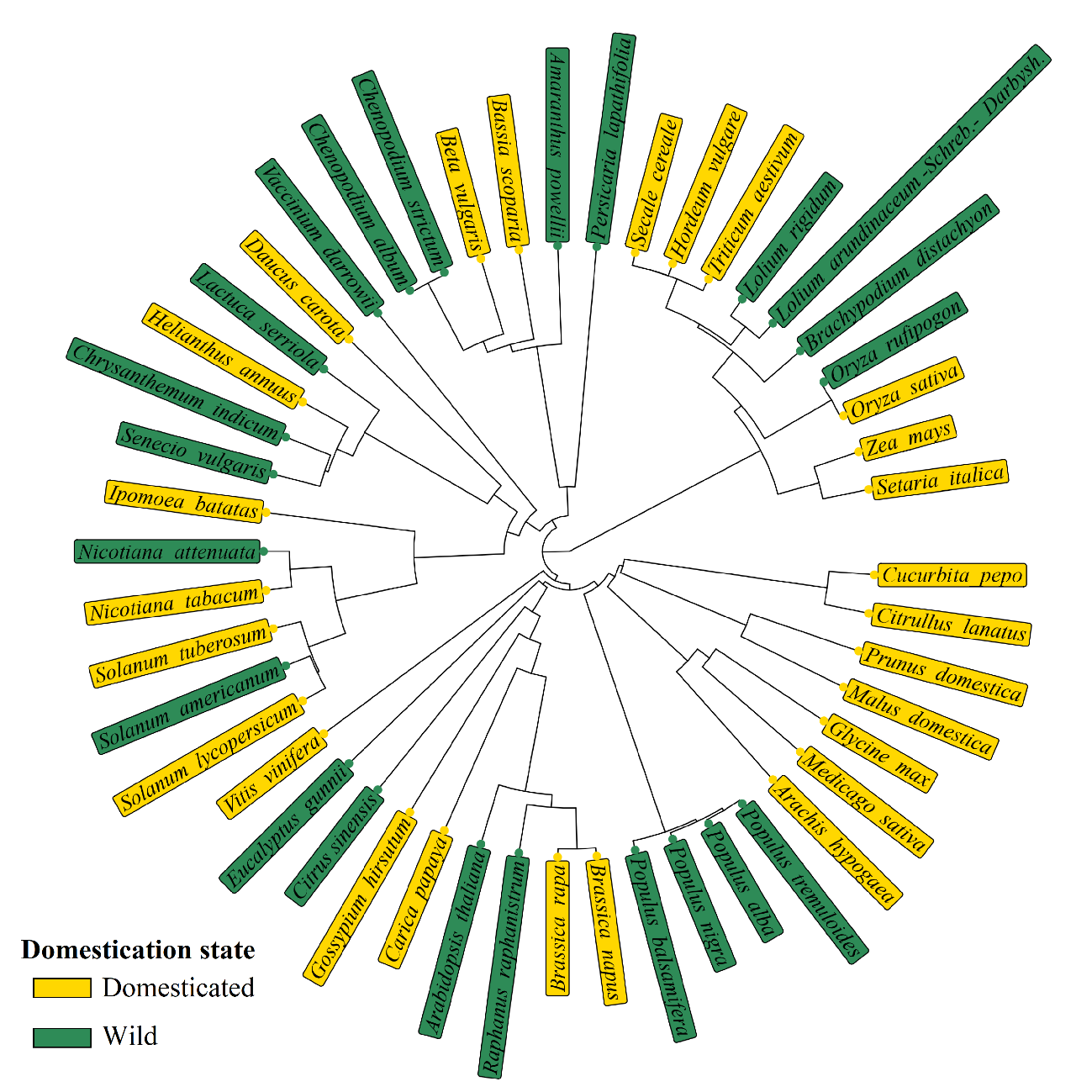
**Extended Data Fig. 1 Circular phylogenetic tree of the studied taxa.** Branch lengths represent genetic distance (scale bar is not shown). Tips are labeled with species names. Background color of each tip label indicates domestication status: yellow for domesticated taxa and green for wild taxa.

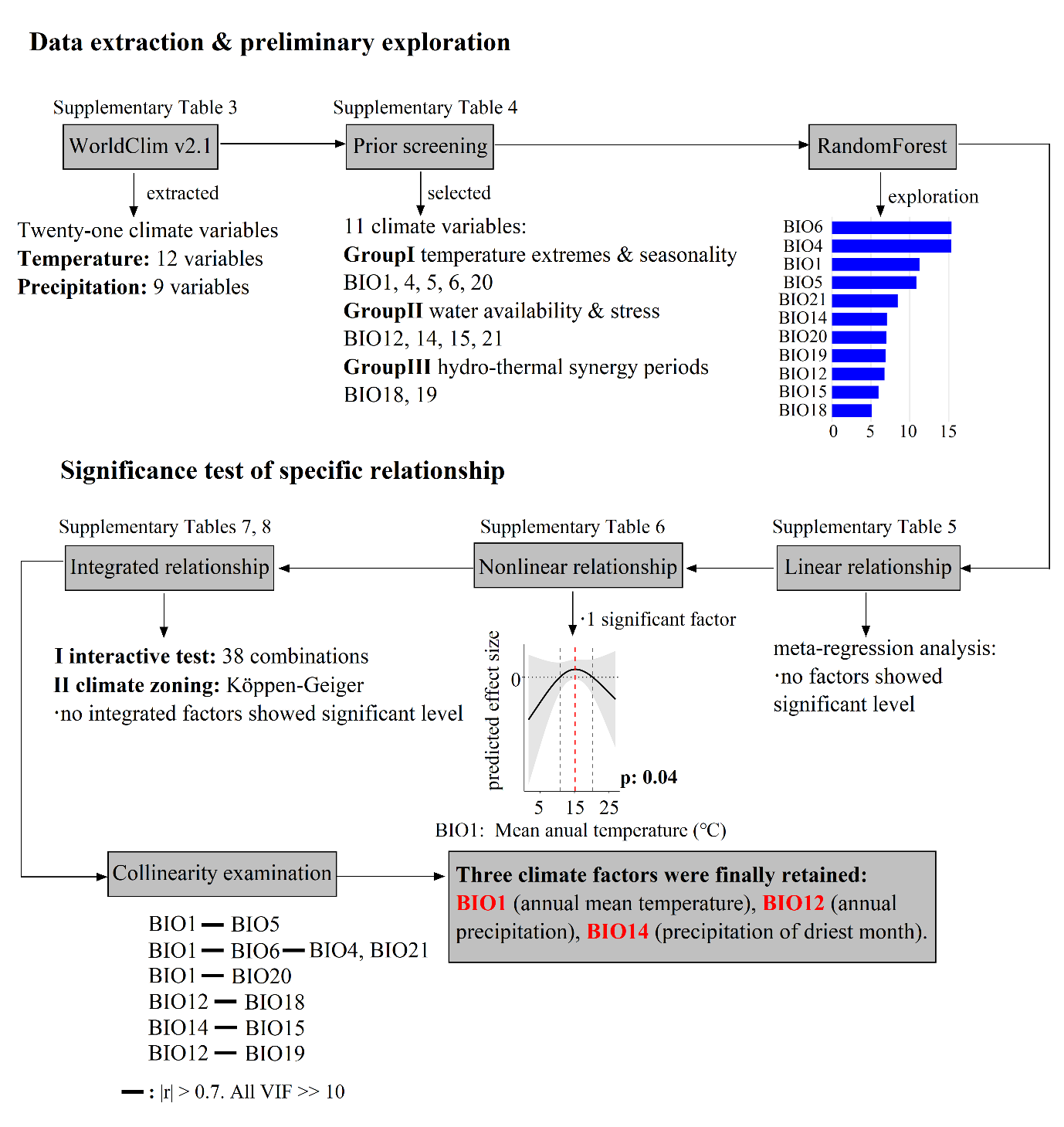

**Extended Data Fig. 2 Climate data collection for this meta-analysis.** A diagram shows the process of climate data from extraction to final including in this meta-analysis.

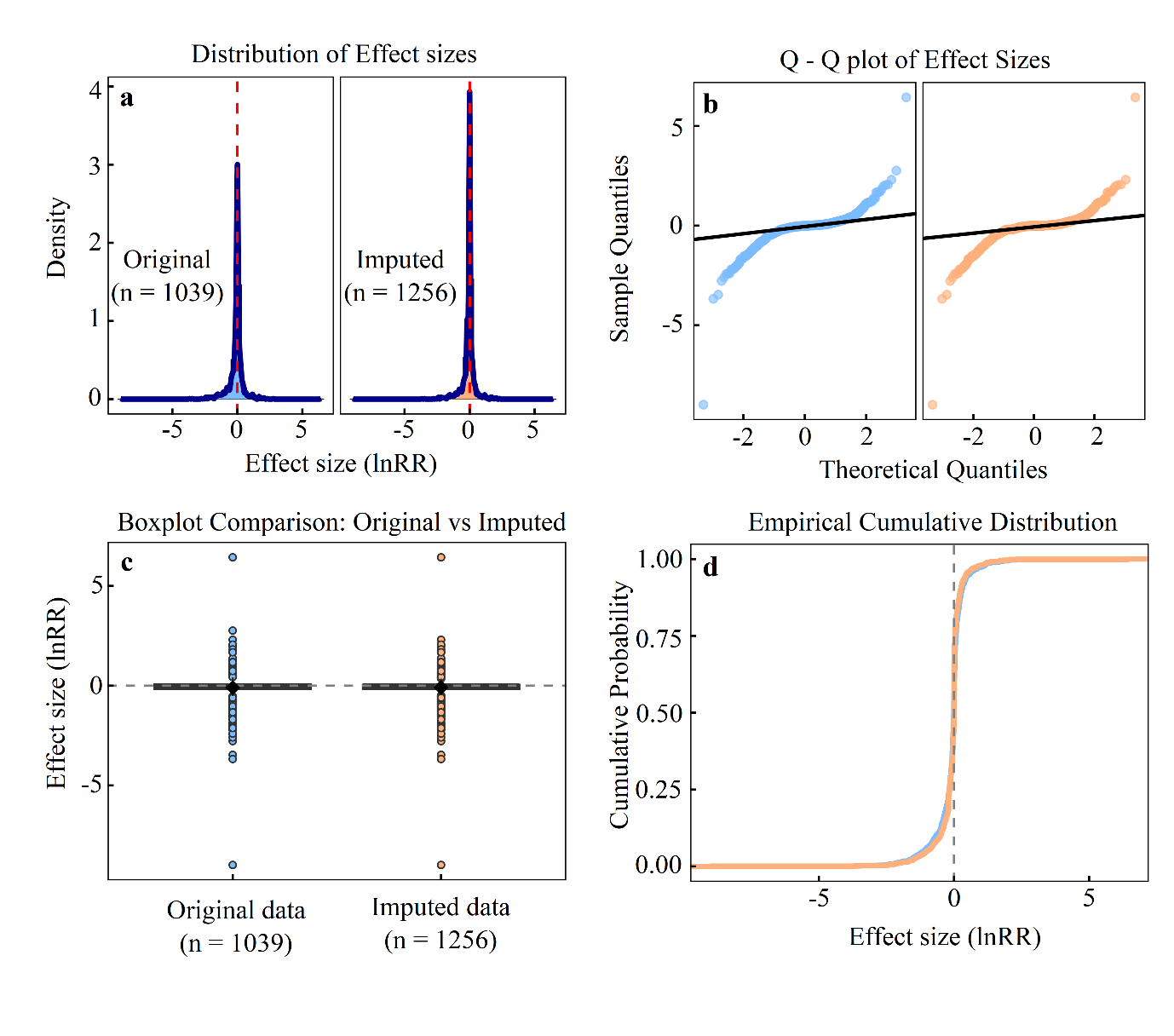
 **Extended Data Fig. 3 Distribution of effect sizes (lnRR) for complete‑case and imputed datasets. a,** Histograms with overlaid density curves showing the distribution of lnRR for the original dataset and the imputed dataset (n = 1,256; orange). **b,** Comparison of the sample quantiles against theoretical normal quantiles for the original (blue) and imputed (orange) datasets. **c,** Boxplots summarizing the central tendency and spread of effect sizes for both datasets. **d,** Empirical cumulative distribution functions (ECDFs) for the two datasets, illustrating the cumulative probability of effect sizes across the observed range. In all panels, the distributions are unimodal and centred near zero, with a slightly longer left tail that pulls the mean below the median. The imputed data closely reproduce the distributional features of the complete‑case data, confirming that the imputation procedure did not introduce systematic bias. See Methods for detailed imputation procedure.

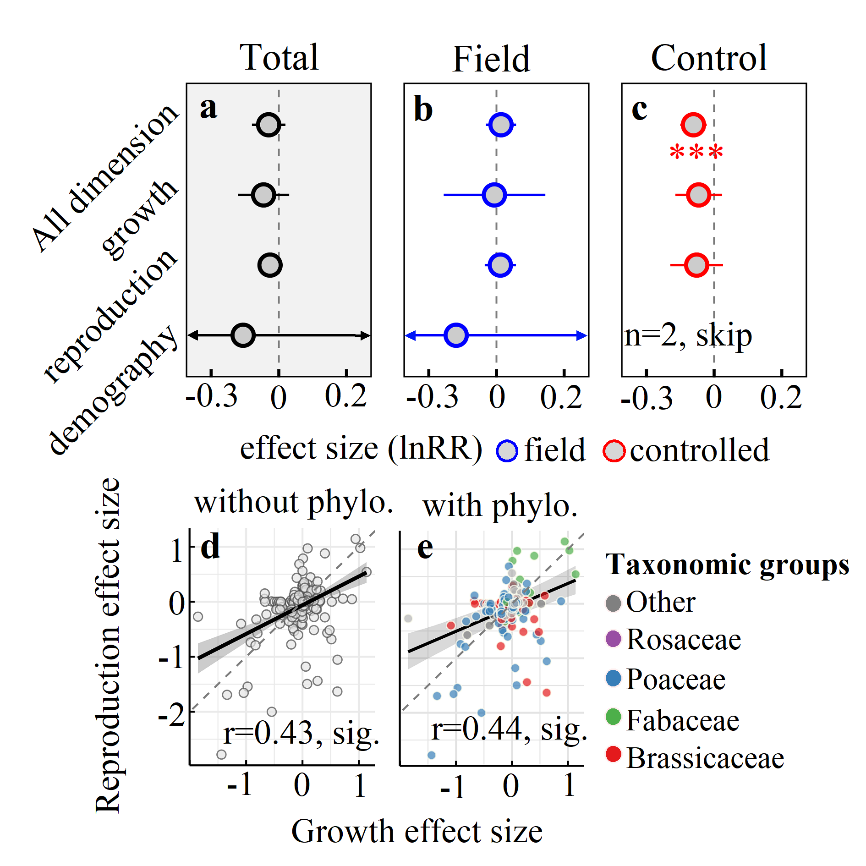

**Extended Data Fig. 4 The fitness effect of resistant transgenes across all dimensions and on different** **fitness dimensions.** For the total dataset **(a)**, field subset **(b)**, and control subset **(c)**, effect sizes (lnRR) across all dimensions and on different dimensions were estimated with phylogenetic correction. All models were constructed using a five-level random effect structure: (~ 1 | Study_ID), (~ 1 | Exp_ID), (~1|Study ID: Control ID), (~ 1 | Observation ID), (~ 1 | Species). Significance levels: *P < 0.05, **P < 0.01, ***P < 0.001. Significant linear relationships of growth and reproduction at the experiment-level (n = 221) before **(d)** and after **(e)** the correction of phylogenetic signal, with a slope approximately 0.4.

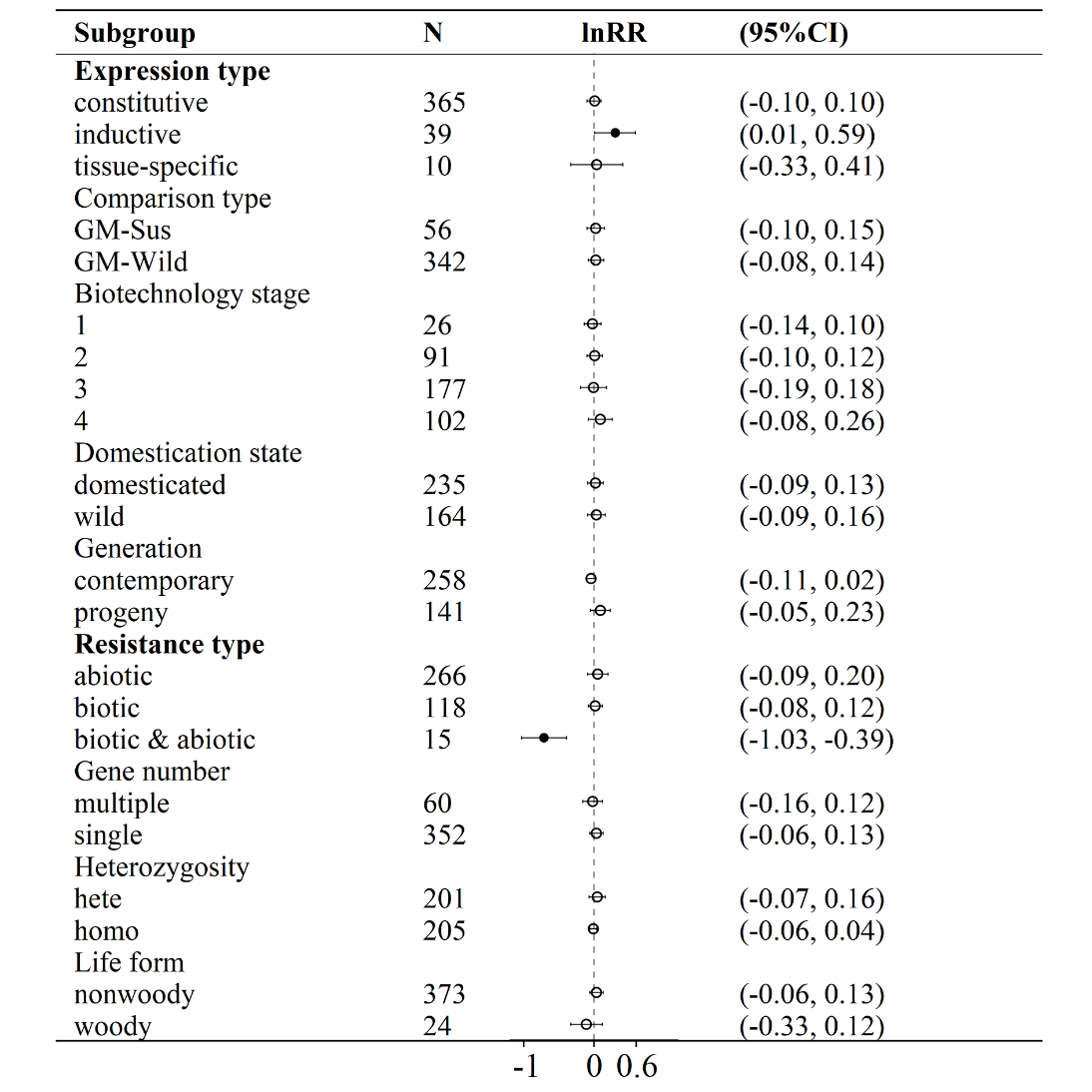

**Extended Data Fig. 5 The effect of different levels of modulators on fitness effect in the field subset.** The solid circle represents the significant subgroup levels in the meta-regression analysis, while the hollow circle represents the insignificant ones. The significant subgroups in meta analysis are marked in **bold**.

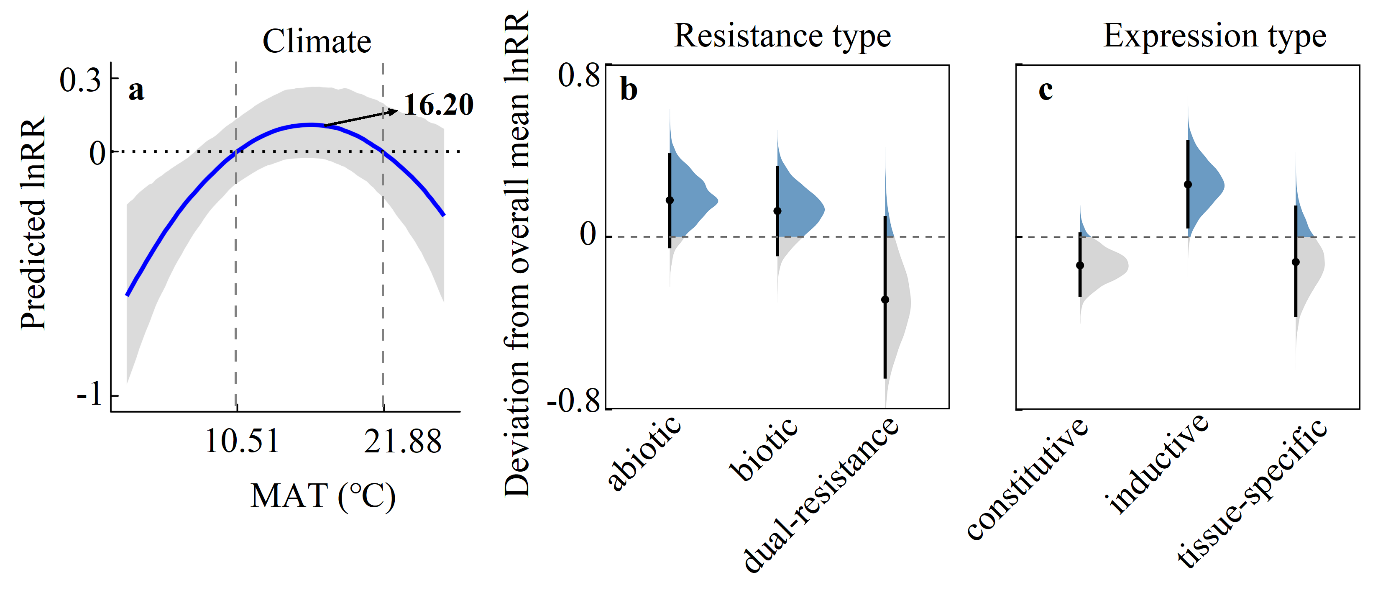

**Extended Data Fig. 6 The results of the Bayesian model for the field subset. (a)** Bayesian model (506 field observations, without phylogenetic correction) confirms the single-peak relationship, controlling two categorical moderators: resistance type **(b)** and expression type **(c)**. For categorical moderators, deviation from overall mean (sum to zero contrast) quantifies each level’s independent contribution; points ± bars = median ± 95% CI.
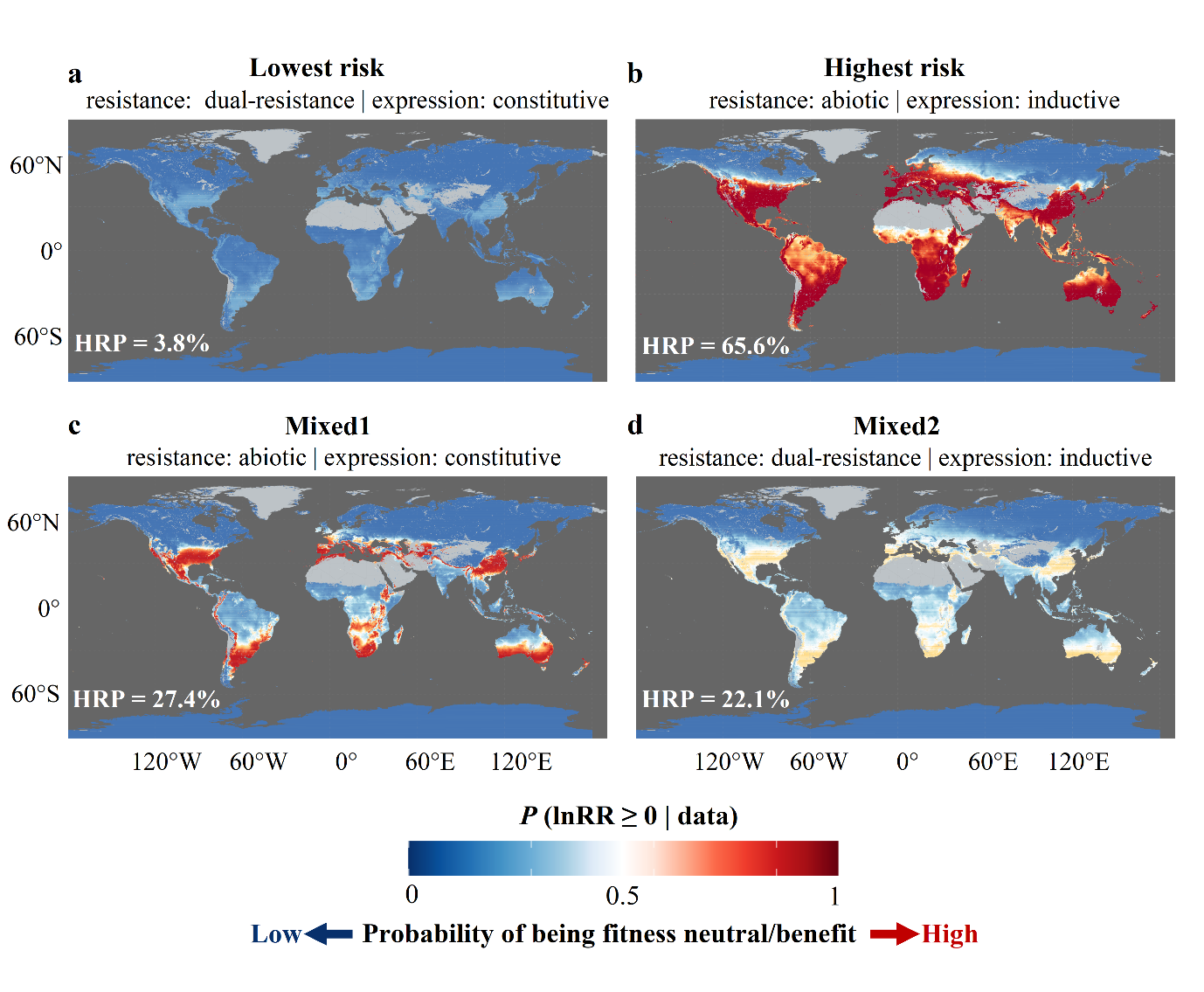
 **Extended Data Fig. 7 Extrapolated global risk maps (full temperature range).** Colors represent the posterior probability that effect size (lnRR) ≥ 0: Blue shades (0–0.5) indicate that a fitness cost is more likely (low risk), red shades (0.5–1) indicate that a fitness benefit or neutrality is more likely (high risk). HRP (High‑Risk Proportion) is the area‑weighted posterior probability that the lnRR ≥ 0, representing the expected fraction of vegetated land with high ecological risk of escaped GMPs. Gray areas represent non‑vegetated land or ocean. Predictions are extrapolated to the full global temperature range (no clamping) for four risk scenarios under vegetation mask: **(a)** Largest cost (biotic & abiotic resistance + constitutive expression). **(b)** Largest benefit (abiotic resistance + inductive expression). **(c)** Mixed1 (abiotic resistance + constitutive expression). **(d)** Mixed2 (biotic & abiotic resistance + inductive expression). Climate data: WorldClim v2.1 (BIO1, 1970–2000).
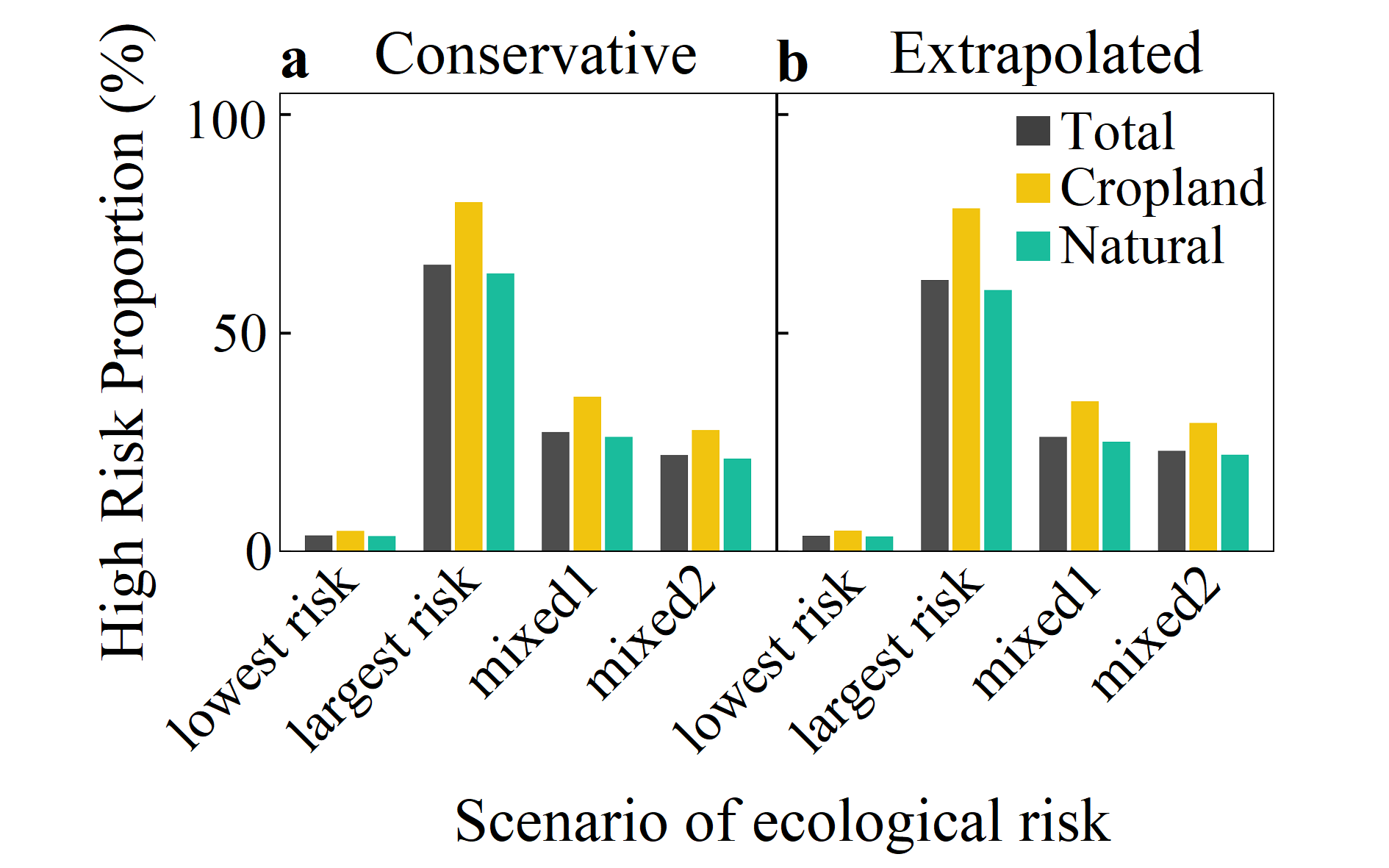

**Extended Data Fig. 8 Expected High‑Risk Proportion (HRP) for four transgenic scenarios under conservative (restricted to training temperature 1.7–26.8 °C) and extrapolated (full‑range) predictions.** HRP is the area‑weighted average of the cell‑wise probabilities that lnRR ≥ 0, representing the expected fraction of vegetated land with high ecological risk of escaped GMPs. Bars: total vegetation (black), cropland (yellow), natural vegetation (green). As all pixel-level HRP values have formed an extremely wide confidence interval globally, only bar charts are presented for their posterior median, which are sorted robustly.

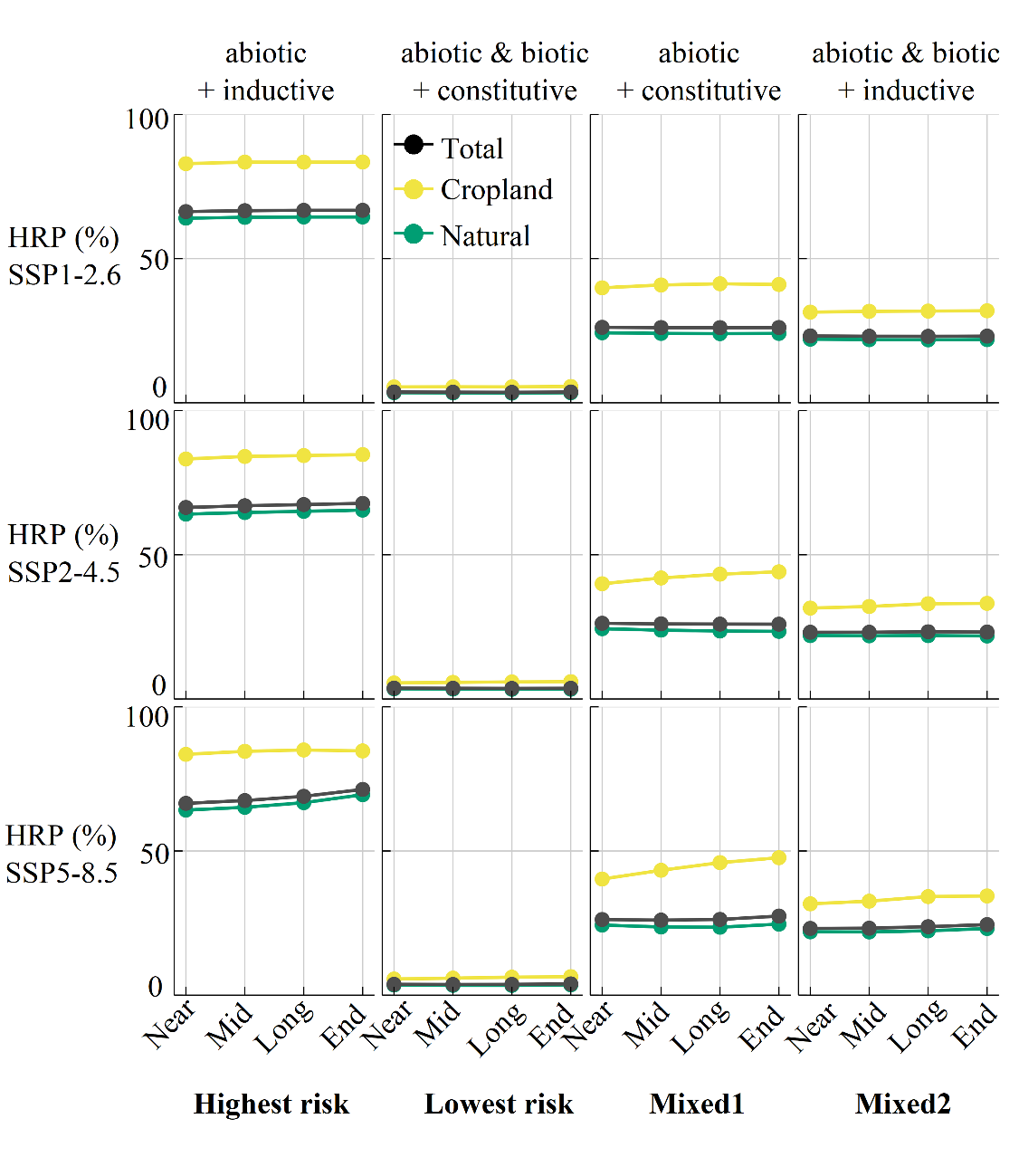

**Extended Data Fig. 9 Constrained prediction of future high‑risk area proportion (HRP) in global vegetation under four risk scenarios and three Shared Socioeconomic Pathways (SSPs).** HRP (High‑Risk Proportion) is defined as the expected fraction of vegetated land (MODIS IGBP classes 1‑12 and 14) with high ecological risk of escaped GMPs. Future climate inputs are from five CMIP6 models (ISIMIP3b subset). Four future periods (Near: 2021‑2040, Mid: 2041‑2060, Long: 2061‑2080, End: 2081‑2100) are considered under three SSPs: SSP1‑2.6 (low emissions), SSP2‑4.5 (intermediate emissions), and SSP5‑8.5 (high emissions). Predictions are constrained to the training temperature range (i.e., clipped to 1.7‑26.8 °C, no extrapolation). As all pixel-level HRP values have formed an extremely wide confidence interval globally, only bar charts are presented for their posterior median, which are sorted robustly.

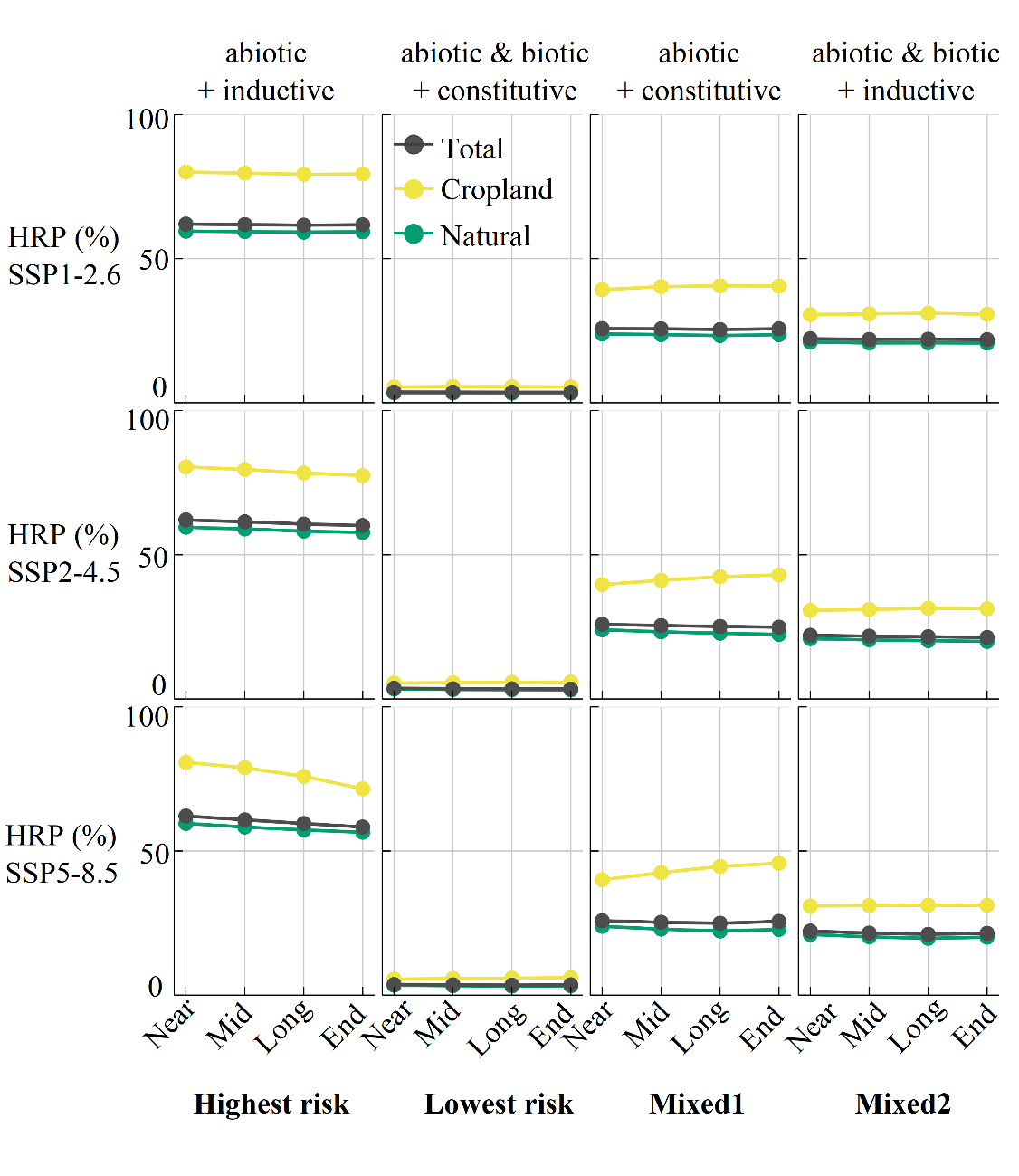

**Extended Data Fig. 10 Extrapolated prediction of future high‑risk area proportion (HRP) in global vegetation under four risk scenarios and three Shared Socioeconomic Pathways (SSPs).** Predictions are based on the Bayesian model fitted to field data (see Methods). HRP (High‑Risk Proportion) is defined as the area‑weighted posterior probability that lnRR  ≥ 0, representing the expected fraction of vegetated land (MODIS IGBP classes 1‑12 and 14) with high ecological risk of escaped GMPs. Future climate inputs are from five CMIP6 models (ISIMIP3b subset). Four future periods (Near: 2021‑2040, Mid: 2041‑2060, Long: 2061‑2080, End: 2081‑2100) are considered under three SSPs: SSP1‑2.6 (low emissions), SSP2‑4.5 (intermediate emissions), and SSP5‑8.5 (high emissions). Predictions allow full temperature extrapolation (i.e., no clipping to the training range of 1.7‑26.8 °C). As all pixel-level HRP values have formed an extremely wide confidence interval globally, only bar charts are presented for their posterior median, which are sorted robustly.
