## Supplementary material for "Temperature governs fitness effects of transgenes on plants non-monotonically: a global meta-analysis": The Supplementary Tables

**Supplementary Table 1**

**The summary and description of 38 fitness indices included.** Since the effect size adopts the response ratio (lnRR, Equation 1 in Method), the measurement units among studies were not converted.

| Dimension | Fitness index | Description |
| --- | --- | --- |
| Growth | total leaf area | The inner area of the leaf margin or using the length and/or width of the leaf. |
|  | plant length | The height above ground of a plant, excluding roots. |
|  | stem biomass | Dry weight of stems of non-woody plants or branches of woody plants. |
|  | aboveground biomass | Dry weight of aboveground parts of plants. |
|  | total biomass | The dry weight of the aboveground and underground parts of the plant, excluding seeds. |
|  | description of phenotype | Estimated according to the description given by the authors when the data cannot be obtained. |
|  | fresh weight | The fresh weight of aboveground parts of plants. |
|  | photosynthetic rate | Maximum photosynthetic rate per unit leaf area |
|  | relative growth rate | Calculated by the growth rate of dry weight, cell area or length of stem. |
|  | rosette diameter | The rosette diameter of herbaceous plants. |
|  | stem diameter | Diameter of stems of non-woody plants or branches of woody plants. |
|  | leaf number | The total number of leaves for non-woody plants |
| Reproduction | siliques per plant | The number of siliques or spikelet (for crops) on each plant. |
|  | seeds per silique | The number of seeds in each silique or spikelet (for crops). |
|  | seeds per plant | The number of seeds on each plant. |
|  | germination rate | The ratio of seeds number germinating normally in the total number of tested seeds. |
|  | pollen germination | Percentage of pollen that successfully germinates within a certain period of time. |
|  | pollen number | Number of pollen grains per unit area or per unit volume. |
|  | seed setting rate | The ratio of filled grains to the total number of flowers in cereal crops. |
|  | tillers per plant | The number of effective tillers per plant (for crops). |
|  | thousand-seed weight | The weight of 1,000 seeds from a particular batch. |
|  | seed biomass | Total seed weight per plant or hectare. |
|  | seed size | Calculated from the length and width of the seed or fruit. |
|  | fruits per plant | Number of fruits per plant. |
|  | fresh weight per fruit | Fresh weight of per fruit. |
|  | description of fertility | Estimated according to the description given by the authors when the data cannot be obtained. |
|  | pollen viability | The ability of the pollen to perform its function of delivering male gametes to the embryo sac. |
|  | spikelet fertility | The number of fertile florets in plant spikelets, assessing crop yield potential. |
|  | filling rate | The grain filling rate. |
|  | pollens per plant | Number of pollen grains per plant. |
|  | fruit biomass | The total dry weight of the fruit on each plant. |
|  | bolting rate | The proportion of crops bolting in a specific period of time. |
|  | silique length | The length of siliques or spikelets (for crops). |
|  | flower biomass | The total number or total dry weight of flowers on each plant. |
| Demography | gene frequency | Some studies also reported it using genotype frequency. |
|  | survival rate | The ratio of the number of individuals in a population to the previous year. |
|  | lambda | The ratio of the increment of population number to the original population number. |
|  | replacement capacity | Survival Rate or Establishment Rate. |

**Supplementary Table 2**

**The summary and description of 13 moderators included.** Yes or No indicates whether the moderator was included in the analysis of the data set. HR: transgenic plants with high resistance, LR: transgenic plants with low‑resistance or wild type.

| Category | Moderator | Description and different levels under subgroup | Total dataset | Control dataset |  | Field dataset |
| --- | --- | --- | --- | --- | --- | --- |
| genetic feature | Domesticated state | Transgenic host plants were domesticated/wild | Yes | Yes |  | Yes |
|  | Gene number | The number of transgenes | Yes | Yes |  | Yes |
|  | Life form | Transgenic host plants were woody/nonwoody | Yes | Yes |  | Yes |
|  | Homozygosity | Homozygosity of transgenes | Yes | Yes |  | Yes |
| study design | Resistance type | Transgenic resistance was abiotic/biotic/abiotic & biotic | Yes | Yes |  | Yes |
|  | Expression type | Transgenes in HR were constitutive/inductive/tissue-specifc expression | Yes | Yes |  | Yes |
|  | Comparison type | LR in an experiment was wild type/susceptible plants | Yes | Yes |  | Yes |
|  | Generation | Fitness indices were measured on the first generation of transgenic plants or their descendants | Yes | Yes |  | Yes |
|  | Biotechnology stage | The historical progression of plant biotechnology was delineated into four recognized stages | Yes | Yes |  | Yes |
| environmental context | Experiment condition | Fitness indices were measured in a controlled/field condition | Yes | No |  | No |
|  | PDM | Precipitation of driest month, continuous variable | No | No |  | Yes |
|  | MAT | Mean annual temperature; continuous variable | No | No |  | Yes |
|  | MAP | Mean annual precipitation; continuous variable | No | No |  | Yes |

**Supplementary Table 3**

**The summary and description of 21 climate variables extracted.** The assigned abbreviation, full name, unit, and descriptions of ecological significance of all factors are based on the original WorldClim naming.

| Category | ClimID | Abbreviation | Climate Factor | Unit | Description |
| --- | --- | --- | --- | --- | --- |
| Temperature | BIO1 | MAT | Mean Annual Temperature | °C | Overall thermal regime, fundamental for metabolic rates and species distribution. |
|  | BIO2 | MDR | Mean Diurnal Range | °C | Daily temperature fluctuation, affects physiological stress and diurnal activity. |
|  | BIO3 | ISO | Isothermality | % | Ratio of diurnal to annual temperature variation; indicates how much daily variation contributes to total annual range. |
|  | BIO4 | TSea | Temperature Seasonality | °C | Annual temperature variability; higher values indicate greater seasonal contrast. |
|  | BIO5 | MTWM | Max Temperature of Warmest Month | °C | Upper thermal extreme, important for heat stress tolerance. |
|  | BIO6 | MTCM | Min Temperature of Coldest Month | °C | Lower thermal extreme, critical for cold tolerance and winter survival. |
|  | BIO7 | TAR | Temperature Annual Range | °C | Amplitude of annual temperature extremes, influences life‑history strategies. |
|  | BIO8 | MTWetQ | Mean Temperature of Wettest Quarter | °C | Thermal conditions during the peak precipitation season. |
|  | BIO9 | MTDryQ | Mean Temperature of Driest Quarter | °C | Thermal conditions during the driest period, affects drought stress. |
|  | BIO10 | MTWarmQ | Mean Temperature of Warmest Quarter | °C | Temperature during the warmest three months, relevant for growing season. |
|  | BIO11 | MTColdQ | Mean Temperature of Coldest Quarter | °C | Temperature during the coldest three months, influences dormancy and winter hardiness. |
|  | BIO20 | SRAD | Mean Annual Solar Radiation | kJ m⁻² day⁻¹ | Energy input for photosynthesis, evapotranspiration, and heat balance. |
| Precipitation | BIO12 | MAP | Mean Annual Precipitation | cm | Total annual water input, primary determinant of water availability. |
|  | BIO13 | PWM | Precipitation of Wettest Month | mm | Intensity of the wettest period, relevant for flooding or excess moisture stress. |
|  | BIO14 | PDM | Precipitation of Driest Month | mm | Severity of the driest period, key for drought tolerance. |
|  | BIO15 | PSea | Precipitation Seasonality | % | Temporal distribution of precipitation; high values indicate strong seasonal contrasts. |
|  | BIO16 | PWQ | Precipitation of Wettest Quarter | mm | Total precipitation during the three wettest consecutive months. |
|  | BIO17 | PDQ | Precipitation of Driest Quarter | mm | Total precipitation during the three driest consecutive months. |
|  | BIO18 | PWarmQ | Precipitation of Warmest Quarter | mm | Precipitation during the warmest three months, often corresponds to growing season rainfall. |
|  | BIO19 | PColdQ | Precipitation of Coldest Quarter | mm | Precipitation during the coldest three months, influences winter water balance. |
|  | BIO21 | VAPR | Annual Mean Water Vapor Pressure | kPa | Air moisture content; higher values indicate more humid conditions, lower values suggest atmospheric dryness. |

**Supplementary Table 4**

**Prior screening of climate variables.** Based on the literature support in the "Ref." column, 11 variables that may have an impact on the trade-off between growth and defense were selected from 21 environmental variables.

| Category | Abbreviation | Moderators | Unit | Description | Ref. |
| --- | --- | --- | --- | --- | --- |
| temperature extremes & seasonality | MAT | Mean Annual Temperature | °C | Overall thermal regime, fundamental for metabolic rates and species distribution. | ^1^ |
|  | TSea | Temperature Seasonality | °C | Annual temperature variability; higher values indicate greater seasonal contrast. | ^2,3^ |
|  | MTWM | Max Temperature of Warmest Month | °C | Upper thermal extreme, important for heat stress tolerance. | ^4^ |
|  | MTCM | Min Temperature of Coldest Month | °C | Lower thermal extreme, critical for cold tolerance and winter survival. | ^5,6^ |
|  | SRAD | Mean Annual Solar Radiation | kJ m⁻² day⁻¹ | Energy input for photosynthesis, evapotranspiration, and heat balance. | ^7^ |
| water availability & stress | MAP | Mean Annual Precipitation | cm | Total annual water input, primary determinant of water availability. | ^8,9^ |
|  | PDM | Precipitation of Driest Month | mm | Severity of the driest period, key for drought tolerance. | ^10^ |
|  | PSea | Precipitation Seasonality | % | Temporal distribution of precipitation; high values indicate strong seasonal contrasts. | ^3^ |
|  | VAPR | Annual Mean Water Vapor Pressure | kPa | Air moisture content; higher values indicate more humid conditions, lower values suggest atmospheric dryness. | ^11^ |
| hydro-thermal synergy periods | PWarmQ | Precipitation of Warmest Quarter | mm | Precipitation during the warmest three months, often corresponds to growing season rainfall. | ^12^ |
|  | PColdQ | Precipitation of Coldest Quarter | mm | Precipitation during the coldest three months, influencing winter water balance. | ^13^ |

**Supplementary Table 5**

**The results of meta-regression analysis of 11 climate variables.** The random-effect of analysis was ~Study ID, ~Experiment ID, ~Control ID, which is consistent with the structure of other meta-analysis in this study. The full name of variables was provided in Supplementary Table 3.

| Variables | beta | SE | t_stat | p_val | q_fdr |
| --- | --- | --- | --- | --- | --- |
| BIO1 | 0.0116026 | 0.0106581 | 1.0886128 | 0.2861672 | 0.6478638 |
| BIO4 | -0.000289 | 0.000236 | -1.223579 | 0.2334411 | 0.6478638 |
| BIO5 | 0.0094662 | 0.0129367 | 0.7317301 | 0.4728383 | 0.6478638 |
| BIO6 | 0.010994 | 0.0082384 | 1.3344834 | 0.1929784 | 0.6478638 |
| BIO20 | 7.52E-06 | 1.72E-05 | 0.4379183 | 0.6684313 | 0.6684313 |
| BIO12 | 4.99E-05 | 8.57E-05 | 0.5821877 | 0.5676575 | 0.6478638 |
| BIO14 | -0.00127 | 0.0020393 | -0.622921 | 0.5400753 | 0.6478638 |
| BIO15 | 0.0013274 | 0.0015575 | 0.8522846 | 0.4039931 | 0.6478638 |
| BIO21 | 0.1396437 | 0.1079007 | 1.2941866 | 0.2081432 | 0.6478638 |
| BIO18 | -9.88E-05 | 0.0001801 | -0.548915 | 0.588967 | 0.6478638 |
| BIO19 | 0.0001784 | 0.0003025 | 0.5897992 | 0.5663204 | 0.6478638 |

**Supplementary Table 6**

**Maximum Likelihood Ratios (LRT) of Restricted Cubic Spline Analysis for 11 climatic variables and their FDR corrected results.** The red font represents a variable that has a significant nonlinear relationship with the effect size. The full name of variables was provided in Supplementary Table 3.

| Variables | Variables | ClimID | LRT_pval | FDR_pval |
| --- | --- | --- | --- | --- |
| Mean Annual Temperature | MAT | BIO1 | 0.0412 | 0.0412 |
| Min Temperature of Coldest Month | MTCM | BIO6 | 0.1325 | 0.2915 |
| Mean Annual Radiation | SRAD | BIO20 | 0.16 | 0.2933 |
| Max Temperature of Warmest Month | MTWM | BIO5 | 0.2287 | 0.3494 |
| Precipitation Seasonality | PSea | BIO15 | 0.3521 | 0.4577 |
| Annual Mean Water Vapor Pressure | VAPR | BIO21 | 0.4847 | 0.5332 |
| Precipitation of Coldest Quarter | PColdQ | BIO19 | 0.5032 | 0.5332 |
| Precipitation of Warmest Quarter | PWarmQ | BIO18 | 0.5161 | 0.5332 |
| Mean Annual Precipitation | MAP | BIO12 | 0.7498 | 0.7498 |
| Precipitation of Driest Month | PDM | BIO14 | 0.9912 | 0.9912 |
| Temperature Seasonality | TSea | BIO4 | 0.4119 | 0.5134 |

**Supplementary Table 7**

**Test for all interaction terms of climate variables in three categories.** The random-effect of analysis was ~Study ID, ~Experiment ID, ~Control ID, which is consistent with the structure of other meta-analysis in this study. The category of climate variables was provided in Supplementary Table 3.

| Interaction | beta | SE | t_stat | p_value | q_fdr |
| --- | --- | --- | --- | --- | --- |
| BIO1:BIO12 | 2.91E-05 | 3.25E-05 | 0.8948787 | 0.3830296 | 0.6328315 |
| BIO1:BIO14 | 0.0007162 | 0.000566 | 1.2653814 | 0.2281286 | 0.6316829 |
| BIO1:BIO15 | -0.000575 | 0.0002759 | -2.085099 | 0.093179 | 0.6316829 |
| BIO1:BIO21 | -0.020269 | 0.0175159 | -1.157188 | 0.2722341 | 0.6316829 |
| BIO4:BIO12 | -3.47E-07 | 3.01E-07 | -1.151744 | 0.3185932 | 0.6316829 |
| BIO4:BIO14 | -1.29E-05 | 1.03E-05 | -1.253166 | 0.2621167 | 0.6316829 |
| BIO4:BIO15 | 8.06E-06 | 7.53E-06 | 1.0708014 | 0.2989093 | 0.6316829 |
| BIO4:BIO21 | 0.0004748 | 0.0004832 | 0.9826399 | 0.3461143 | 0.6316829 |
| BIO5:BIO12 | 1.76E-05 | 3.59E-05 | 0.4902763 | 0.6406715 | 0.798415 |
| BIO5:BIO14 | 0.0004428 | 0.0006832 | 0.648087 | 0.5298007 | 0.7743241 |
| BIO5:BIO15 | -0.000519 | 0.0002916 | -1.778744 | 0.1542729 | 0.6316829 |
| BIO5:BIO21 | -0.041339 | 0.0349385 | -1.183182 | 0.274779 | 0.6316829 |
| BIO6:BIO12 | 1.82E-05 | 1.54E-05 | 1.1809301 | 0.2643677 | 0.6316829 |
| BIO6:BIO14 | 0.0005562 | 0.0003727 | 1.4922582 | 0.1687964 | 0.6316829 |
| BIO6:BIO15 | -0.000449 | 0.0002533 | -1.770977 | 0.1023811 | 0.6316829 |
| BIO6:BIO21 | -0.01176 | 0.0125349 | -0.93817 | 0.3657112 | 0.6316829 |
| BIO20:BIO12 | 2.97E-08 | 6.44E-08 | 0.4611187 | 0.6513385 | 0.798415 |
| BIO20:BIO14 | 1.09E-06 | 1.12E-06 | 0.9675746 | 0.3534468 | 0.6316829 |
| BIO20:BIO15 | -7.51E-07 | 3.47E-07 | -2.164025 | 0.0825747 | 0.6316829 |
| BIO20:BIO21 | -9.56E-05 | 6.17E-05 | -1.548894 | 0.1460168 | 0.6316829 |
| BIO1:BIO18 | 6.80E-05 | 6.74E-05 | 1.0081647 | 0.33496 | 0.6316829 |
| BIO1:BIO19 | 5.97E-05 | 0.000104 | 0.5744681 | 0.5775955 | 0.7822215 |
| BIO4:BIO18 | -1.80E-07 | 6.33E-07 | -0.283861 | 0.7841391 | 0.916045 |
| BIO4:BIO19 | -1.63E-06 | 1.40E-06 | -1.170828 | 0.2956994 | 0.6316829 |
| BIO5:BIO18 | 0.0001368 | 0.0001166 | 1.1735829 | 0.2628569 | 0.6316829 |
| BIO5:BIO19 | 1.69E-05 | 0.0001137 | 0.1483608 | 0.8856821 | 0.916045 |
| BIO6:BIO18 | 1.66E-05 | 3.05E-05 | 0.5455975 | 0.5969585 | 0.7822215 |
| BIO6:BIO19 | 6.63E-05 | 6.23E-05 | 1.0635618 | 0.3269414 | 0.6316829 |
| BIO20:BIO18 | 2.27E-07 | 1.48E-07 | 1.5352575 | 0.1448902 | 0.6316829 |
| BIO20:BIO19 | 5.80E-09 | 1.65E-07 | 0.0351058 | 0.9729005 | 0.9729005 |
| BIO12:BIO18 | -4.67E-07 | 4.46E-07 | -1.048396 | 0.3276504 | 0.6316829 |
| BIO12:BIO19 | 1.33E-07 | 4.84E-07 | 0.2755513 | 0.8044513 | 0.916045 |
| BIO14:BIO18 | -1.78E-06 | 1.26E-05 | -0.141713 | 0.8919385 | 0.916045 |
| BIO14:BIO19 | 1.75E-06 | 7.81E-06 | 0.2240961 | 0.8318628 | 0.916045 |
| BIO15:BIO18 | 6.84E-06 | 1.03E-05 | 0.6625957 | 0.5188681 | 0.7743241 |
| BIO15:BIO19 | 7.81E-06 | 1.06E-05 | 0.7359535 | 0.4738325 | 0.7502348 |
| BIO21:BIO18 | -0.00026 | 0.0004373 | -0.594462 | 0.5797109 | 0.7822215 |
| BIO21:BIO19 | 0.0001692 | 0.0009118 | 0.185591 | 0.8573495 | 0.916045 |

**Supplementary Table 8**

**Meta‑regression results for resistance cost across Köppen‑Geiger climate zones (CR2 robust standard errors).** Overall test for climate zone effect: QM = 1.54, df = 3, p = 0.6725. Robust standard errors are cluster‑corrected (CR2) at the random-effect of analysis was ~Study ID, ~Experiment ID, ~Control ID, which is consistent with the structure of other meta-analysis in this study. N represents the number of experiments.

| Climate zone | N | Estimate (lnRR) | SE (CR2) | 95% CI | p‑value |
| --- | --- | --- | --- | --- | --- |
| Temperate (C) | 361 | 0.066 | 0.044 | (-0.020, 0.151) | 0.131 |
| Continental (D) | 103 | -0.055 | 0.115 | (-0.281, 0.172) | 0.635 |
| Arid (B) | 39 | -0.019 | 0.118 | (-0.249, 0.212) | 0.873 |
| Tropical (A) | 3 | 0.025 | 1.162 | (-2.252, 2.302) | 0.983 |

**Supplementary Table 9**

**The number of observations of different magnitudes of fitness effect across three datasets.** Expensive cost: ≤ -0.5; cost: > -0.5 & < -0.01; neutrual: ≥-0.01 & ＜0.01 -; benefit: ≥ 0.01 & ≤ 0.5; substantial benefits: > 0.5.

|  |  | **Number of observations** | | | | | |
| --- | --- | --- | --- | --- | --- | --- | --- |
|  | Subset | Expensive cost | Cost | Neutral | Benefit | Substantial benefit | In total |
| Classified by experiment condition | Control | 91 | 269 | 180 | 183 | 27 | 750 |
|  | Field | 36 | 169 | 94 | 177 | 30 | 506 |
| Classified by fitness dimension | Growth | 51 | 185 | 129 | 138 | 13 | 516 |
|  | Reproduction | 71 | 248 | 139 | 218 | 39 | 715 |
|  | Demography | 5 | 5 | 6 | 4 | 5 | 25 |

**Supplementary Table 10**

**The fitness effect of resistant transgenes on GMPs in stress-free environments before phylogenetic correction.** The demographics dimension was only involved in two experiments in the controlled condition, which did not meet the requirements for conducting a meta-analysis. *M1*, *M2a*, *M2b*, *M2c* represent multilevel meta-analysis on all dimensions, growth-dimension, reproduction-dimension, and demography-dimension, respectively. GMPs, genetically modified plants.

| Dimension | Model | Dataset | Estimate | SE | CI_lower | CI_upper | p_value | QE | I² (%) | n_exp |
| --- | --- | --- | --- | --- | --- | --- | --- | --- | --- | --- |
| Combined | *M1* | Total | -0.095 | 0.025 | -0.144 | -0.047 | 1.45E-04 | 149910.800 | 99.86 | 518 |
|  | *M1* | Field | 0.027 | 0.044 | -0.060 | 0.115 | 0.534 | 49156.985 | 99.89 | 179 |
|  | *M1* | Control | -0.135 | 0.028 | -0.190 | -0.081 | 2.19E-06 | 48843.940 | 99.84 | 339 |
| Growth | *M2a* | Total | -0.139 | 0.026 | -0.190 | -0.087 | 2.95E-07 | 39538.207 | 99.77 | 381 |
|  | *M2a* | Field | -0.021 | 0.049 | -0.119 | 0.078 | 0.676 | 8906.333 | 99.87 | 102 |
|  | *M2a* | Control | -0.162 | 0.030 | -0.220 | -0.103 | 1.54E-07 | 28751.622 | 99.60 | 279 |
| Reproduction | *M2b* | Total | -0.054 | 0.032 | -0.118 | 0.010 | 0.099 | 104774.725 | 99.90 | 342 |
|  | *M2b* | Field | 0.019 | 0.046 | -0.073 | 0.110 | 0.681 | 35579.287 | 99.86 | 152 |
|  | *M2b* | Control | -0.095 | 0.042 | -0.179 | -0.012 | 0.026 | 68777.564 | 99.92 | 190 |
| Demography | *M2c* | Total | -0.211 | 0.339 | -1.040 | 0.618 | 0.557 | 4361.659 | 99.86 | 20 |
|  | *M2c* | Field | -0.212 | 0.376 | -1.178 | 0.754 | 0.597 | 4301.452 | 99.87 | 18 |
|  | *M2c* | Control | - | - | - | - | - | - | - | 2 |

**Supplementary Table 11**

**Sensitivity analyses for the overall effect size under different missing‑data handling methods.** “Best‑case”, “worst‑case” and Variance inflation analysis were applied to the 217 imputed observations.

| Sensitivity analysis | lnRR | SE | 95% CI | p‑value | Remarks |
| --- | --- | --- | --- | --- | --- |
| Imputed (main analysis) | –0.09504 | 0.02462 | [–0.14353, –0.04655] | 0.0001449 | Imputed dataset (n = 1256) |
| Complete‑case | –0.12579 | 0.03311 | [–0.19111, –0.06048] | 0.0001956 | Only original complete observations (n = 1039) |
| Best‑case (revised) | –0.02107 | 0.02531 | [–0.07101, 0.02887] | 0.4056 | Positive imputed → 0.5; negative/zero → 0.2 |
| Worst‑case (revised) | –0.16304 | 0.02535 | [–0.21297, –0.11311] | 6.65E-10 | Positive imputed → –0.2; negative → –0.5; zero → –0.2 |
| Variance × 2 | –0.09576 | 0.02473 | [–0.14448, –0.04705] | 0.0001388 | Imputed variances multiplied by 2 |
| Variance × 5 | –0.09759 | 0.02498 | [–0.14678, –0.04839] | 0.0001211 | Imputed variances multiplied by 5 |
| Variance × 10 | –0.09979 | 0.02528 | [–0.14960, –0.04999] | 0.0001036 | Imputed variances multiplied by 10 |

**Supplementary Table 12**

**The fitness effect of resistant transgenes on GMPs in stress-free environments with phylogenetic correction.** The demographics dimension was only involved in two experiments in the controlled condition, which did not meet the requirements for conducting a meta-analysis. *M1*, *M2a*, *M2b*, *M2c* represent multilevel meta-analysis on all dimensions, growth-dimension, reproduction-dimension, and demography-dimension, respectively. GMPs, genetically modified plants. Var_species (%) represents the species‑level variance component in meta-analysis.

| Dimension | Model | Dataset | Estimate | SE | CI_lower | CI_upper | p_value | QE | Var_species (%) | I² (%) | n_exp |
| --- | --- | --- | --- | --- | --- | --- | --- | --- | --- | --- | --- |
| Combined | *M1* | Total | -0.060 | 0.050 | -0.159 | 0.039 | 0.232 | 149860.913 | 5.4340 | 99.87 | 518 |
|  | *M1* | Field | 0.027 | 0.045 | -0.062 | 0.116 | 0.549 | 48843.838 | 0.3260 | 99.89 | 179 |
|  | *M1* | Control | -0.122 | 0.040 | -0.200 | -0.044 | 0.002 | 101014.489 | 1.6665 | 99.85 | 339 |
| Growth | *M2a* | Total | -0.090 | 0.077 | -0.241 | 0.061 | 0.244 | 39465.732 | 0.0303 | 99.78 | 381 |
|  | *M2a* | Field | -0.012 | 0.153 | -0.312 | 0.289 | 0.939 | 8877.933 | 0.1193 | 99.93 | 102 |
|  | *M2a* | Control | -0.091 | 0.070 | -0.229 | 0.047 | 0.194 | 28700.716 | 0.0212 | 99.61 | 279 |
| Reproduction | *M2b* | Total | -0.052 | 0.033 | -0.116 | 0.013 | 0.116 | 104679.988 | 1.45E-09 | 99.90 | 342 |
|  | *M2b* | Field | 0.023 | 0.047 | -0.069 | 0.115 | 0.620 | 35550.741 | 2.03E-09 | 99.87 | 152 |
|  | *M2b* | Control | -0.103 | 0.080 | -0.259 | 0.053 | 0.194 | 68673.623 | 0.0221 | 99.93 | 190 |
| Demography | *M2c* | Total | -0.211 | 0.339 | -0.875 | 0.453 | 0.534 | 4361.659 | 3.05E-09 | 99.86 | 20 |
|  | *M2c* | Field | -0.238 | 0.424 | -1.068 | 0.592 | 0.574 | 4301.424 | 1.31E-08 | 99.88 | 18 |
|  | *M2c* | Control | - | - | - | - | - | - | - | - | 2 |

**Supplementary Table 13**

**Phylogenetic signal tests for the total dataset and its subsets.** K and λ were calculated using weighted mean effect sizes per species (inverse‑variance weights). P‑values for K were obtained from 999 randomizations; p‑values for λ are from likelihood‑ratio tests. Significant phylogenetic signal (p < 0.05) was only observed for Blomberg's K in the total dataset, but not for Pagel's λ in any dataset, indicating a weak and inconsistent signal.

| Dataset | Blomberg's K | p-value (K) | Pagel's λ | p-value (λ) |
| --- | --- | --- | --- | --- |
| Total dataset | 0.2784 | 0.018 | 0.0068 | 0.969 |
| Control subset | 0.3085 | 0.067 | 0.5795 | 0.051 |
| Field subset | 0.1506 | 0.383 | 0.3781 | 0.511 |

**Supplementary Table 14**

**Bayesian multilevel meta-regression results for effect size (lnRR) in field trials before phylogenetic correction.** For deviation‑coded coefficients of categorical moderators, Bulk_ESS was omitted as they are linear combinations of the original model parameters; all original parameters had Rhat = 1 and Bulk/Tail‑ESS > 1000. The residual standard deviation (σ = 0.35) measures the magnitude of the remaining variation between the observed effect size (lnRR) and the predicted values by the model.

| **Parameter** | **Estimate** | **Est.Error** | **95% CI** | **P(β<0)** | **P(β>0)** | **Rhat** | **Bulk_ESS** |
| --- | --- | --- | --- | --- | --- | --- | --- |
| **Fixed effects (continuous)** | | | | | | | |
| BIO1 (raw) | 0.11 | 0.03 | [0.040, 0.170] | 0.002 | 0.999 | 1 | 4790 |
| BIO1² (raw) | -0.003 | 0.001 | [-0.005, -0.0003] | 0.997 | 0.003 | 1 | 4817 |
| BIO1 (scaled) | 0.61 | 0.038 | [0.219, 0.985] | 0.002 | 0.999 | 1 | 4790 |
| BIO1² (scaled) | -0.107 | 0.001 | [-0.180, -0.031] | 0.997 | 0.003 | 1 | 4817 |
| **Fixed effects (categorical, deviation coding)** | | | | | | | |
| Resistance type: abiotic | 0.149 | 0.097 | [-0.046, 0.339] | 0.07 | 0.93 | 1 | — |
| Resistance type: biotic | 0.104 | 0.093 | [-0.079, 0.287] | 0.139 | 0.861 | 1 | — |
| Resistance type: biotic & abiotic | -0.252 | 0.167 | [-0.575, 0.085] | 0.93 | 0.07 | 1 | — |
| Expression type: constitutive | -0.114 | 0.067 | [-0.242, 0.019] | 0.956 | 0.044 | 1 | — |
| Expression type: inductive | 0.214 | 0.091 | [0.035, 0.394] | 0.01 | 0.99 | 1 | — |
| Expression type: tissue‑specific | -0.1 | 0.115 | [-0.324, 0.128] | 0.81 | 0.19 | 1 | — |
| **Random effects (SD)** | | | | | | | |
| σ (Control_ID) | 0.29 | 0.04 | [0.20, 0.38] | — | — | 1.01 | 1551 |
| σ (Exp_ID) | 0.06 | 0.04 | [0.00, 0.15] | — | — | 1 | 1245 |
| σ (Paper_ID) | 0.24 | 0.06 | [0.12, 0.35] | — | — | 1 | 1426 |
| σ (residual) | 0.35 | 0.01 | [0.32, 0.38] | — | — | 1 | 5509 |

**Supplementary Table 15**

**Bayesian multilevel meta-regression results for effect size (lnRR) in field trials after phylogenetic correction.** For deviation‑coded coefficients of categorical moderators, Bulk_ESS was omitted as they are linear combinations of the original model parameters; all original parameters had Rhat = 1 and Bulk/Tail‑ESS > 1000.

| **Parameter** | **Estimate** | **Est.Error** | **95% CI** | **P(β<0)** | **P(β>0)** | **Rhat** | **Bulk_ESS** |
| --- | --- | --- | --- | --- | --- | --- | --- |
| **Fixed effects (continuous)** | | | | | | | |
| BIO1 (raw) | 0.11 | 0.04 | [0.040, 0.180] | 0.002 | 0.999 | 1 | 4790 |
| BIO1² (raw) | -0.003 | 0.001 | [-0.010, -0.000] | 0.997 | 0.003 | 1 | 4817 |
| BIO1 (scaled) | 0.619 | 0.199 | [0.224, 1.015] | 0.002 | 0.999 | 1 | 4790 |
| BIO1² (scaled) | -0.108 | 0.038 | [-0.183, -0.033] | 0.997 | 0.003 | 1 | 4817 |
| **Fixed effects (categorical, deviation coding)** | | | | | | | |
| Resistance type: abiotic | 0.13 | 0.093 | [-0.052, 0.313] | 0.083 | 0.917 | 1 | — |
| Resistance type: biotic | 0.083 | 0.09 | [-0.095, 0.257] | 0.179 | 0.821 | 1 | — |
| Resistance type: biotic & abiotic | -0.212 | 0.154 | [-0.515, 0.092] | 0.915 | 0.085 | 1 | — |
| Expression type: constitutive | -0.112 | 0.064 | [-0.237, 0.016] | 0.955 | 0.045 | 1 | — |
| Expression type: inductive | 0.211 | 0.093 | [0.025, 0.392] | 0.013 | 0.987 | 1 | — |
| Expression type: tissue‑specific | -0.1 | 0.113 | [-0.318, 0.123] | 0.813 | 0.187 | 1 | — |
| **Random effects (SD)** | | | | | | | |
| σ (Control_ID) | 0.29 | 0.04 | [0.20, 0.38] | — | — | 1 | 1290 |
| σ (Exp_ID) | 0.06 | 0.04 | [0.00, 0.15] | — | — | 1 | 1092 |
| σ (Paper_ID) | 0.23 | 0.06 | [0.10, 0.34] | — | — | 1 | 892 |
| σ (species) | 0.01 | 0.01 | [0.00, 0.03] | — | — | 1 | 1485 |
| σ (residual) | 0.35 | 0.01 | [0.32, 0.38] | — | — | 1 | 5509 |

**Supplementary Table 16**

**The predicted relative percentage change in the High-Risk area Proportion (HRP) of global vegetation by the end of this century.** The climate data used in the Bayesian risk assessment model is derived from global temperature models (WorldClim v2.1) for three Shared Socioeconomic Pathways (SSPs). Change of HRP is calculated by The HRP value at the end value (2081-2100) minus the recent value (2021-2040), then divided by the recent value. Constrained predictions are clipped to the training temperature range (1.7–26.8 °C); extrapolated predictions allow full temperature extrapolation beyond this range. Total vegetation: gray, Cropland vegetation: yellow, Natural vegetation: green.

| **SSP** | **Scenario** | **Vegetation** | **Constrained change (%)** | **Extrapolated change (%)** |
| --- | --- | --- | --- | --- |
| 1‑2.6 | Highest risk | Total | 0.67 | -0.39 |
|  |  | Cropland | 0.7 | -0.85 |
|  |  | Natural | 0.66 | -0.3 |
|  | Lowest risk | Total | -0.02 | -2.47 |
|  |  | Cropland | 2.84 | 0.04 |
|  |  | Natural | -0.64 | -3.02 |
|  | mixed1 | Total | -0.21 | -0.21 |
|  |  | Cropland | 2.9 | 3.17 |
|  |  | Natural | -0.91 | -0.98 |
|  | mixed2 | Total | -0.33 | -1.37 |
|  |  | Cropland | 1.5 | 0.45 |
|  |  | Natural | -0.68 | -1.73 |
| 2‑4.5 | Highest risk | Total | 2.13 | -3.14 |
|  |  | Cropland | 1.83 | -3.75 |
|  |  | Natural | 2.18 | -3.02 |
|  | Lowest risk | Total | -1.33 | -6.2 |
|  |  | Cropland | 8.29 | 5.55 |
|  |  | Natural | -3.41 | -8.78 |
|  | mixed1 | Total | -1.21 | -3.85 |
|  |  | Cropland | 10.38 | 8.5 |
|  |  | Natural | -3.86 | -6.65 |
|  | mixed2 | Total | 0.32 | -3.44 |
|  |  | Cropland | 5.21 | 1.89 |
|  |  | Natural | -0.65 | -4.52 |
| 5‑8.5 | Highest risk | Total | 7.25 | -6.11 |
|  |  | Cropland | 1.41 | -11.36 |
|  |  | Natural | 8.3 | -5.13 |
|  | Lowest risk | Total | 4.29 | -2.61 |
|  |  | Cropland | 14.94 | 9.28 |
|  |  | Natural | 5.41 | -5.23 |
|  | mixed1 | Total | 4.52 | -0.9 |
|  |  | Cropland | 18.37 | 14.45 |
|  |  | Natural | 1.37 | -4.44 |
|  | mixed2 | Total | 5.74 | -3.6 |
|  |  | Cropland | 8.62 | 0.74 |
|  |  | Natural | 5.42 | -4.47 |

**Supplementary Table 17**

**High-Risk area Proportion (HRP) for temperature-restricted in five periods.** HRP (High‑Risk Proportion) is defined as the area‑weighted average of the cell‑wise probabilities that lnRR  ≥ 0, representing the expected fraction of vegetated land (MODIS IGBP classes 1‑12 and 14) with high ecological risk of escaped GMPs. The Bayesian model constructed in this study was based on History-period data (1970-2000). Future climate inputs are from five CMIP6 models (ISIMIP3b subset). Four future periods (Near: 2021‑2040, Mid: 2041‑2060, Long: 2061‑2080, End: 2081‑2100) are considered under three Shared Socioeconomic Pathways (SSPs): SSP1‑2.6 (low emissions), SSP2‑4.5 (intermediate emissions), and SSP5‑8.5 (high emissions). Predictions are constrained to the training temperature range (i.e., clipped to 1.7‑26.8 °C, no extrapolation). dual-resistance: abiotic & biotic resistance.

|  |  |  |  | **High-Risk area Proportion (%)** | | |
| --- | --- | --- | --- | --- | --- | --- |
| Period | Time | Scenario | Description | cropland | natural | Total vegetation |
| History | 1970-2000 | Highest risk | abiotic resistance + inductive expression | 79.9538 | 63.67467619 | 65.642448 |
|  |  | Lowest risk | dual-resistance + constitutive expression | 4.83506 | 3.650649453 | 3.7938177 |
|  |  | mixed1 | abiotic resistance + constitutive expression | 35.4671 | 26.31955306 | 27.425277 |
|  |  | mixed2 | dual-resistance + inductive expression | 27.8798 | 21.33931209 | 22.129903 |
| ssp126_Near | 2021-2040 | Highest risk | abiotic resistance + inductive expression | 82.9206 | 64.0228 | 66.3082 |
|  |  | Lowest risk | dual-resistance + constitutive expression | 5.5136 | 3.5048 | 3.7478 |
|  |  | mixed1 | abiotic resistance + constitutive expression | 39.8663 | 24.2836 | 26.1681 |
|  |  | mixed2 | dual-resistance + inductive expression | 31.4394 | 22.0696 | 23.2028 |
| ssp245_Near | 2021-2040 | Highest risk | abiotic resistance + inductive expression | 83.2311 | 64.1049 | 66.4179 |
|  |  | Lowest risk | dual-resistance + constitutive expression | 5.5591 | 3.5235 | 3.7697 |
|  |  | mixed3 | abiotic resistance + constitutive expression | 39.9568 | 24.3734 | 26.258 |
|  |  | mixed4 | dual-resistance + inductive expression | 31.5205 | 21.9929 | 23.1451 |
| ssp585_Near | 2021-2040 | Highest risk | abiotic resistance + inductive expression | 83.5147 | 64.1866 | 66.524 |
|  |  | Lowest risk | dual-resistance + constitutive expression | 5.5977 | 3.5148 | 3.7667 |
|  |  | mixed3 | abiotic resistance + constitutive expression | 40.272 | 24.3072 | 26.2379 |
|  |  | mixed4 | dual-resistance + inductive expression | 31.6808 | 21.9804 | 23.1535 |
| ssp126_Mid | 2041-2060 | Highest risk | abiotic resistance + inductive expression | 83.4697 | 64.3085 | 66.6257 |
|  |  | Lowest risk | dual-resistance + constitutive expression | 5.5427 | 3.4384 | 3.6928 |
|  |  | mixed5 | abiotic resistance + constitutive expression | 40.856 | 24.0652 | 26.0957 |
|  |  | mixed6 | dual-resistance + inductive expression | 31.7258 | 21.8949 | 23.0838 |
| ssp245_Mid | 2041-2060 | Highest risk | abiotic resistance + inductive expression | 84.1038 | 64.661 | 67.0123 |
|  |  | Lowest risk | dual-resistance + constitutive expression | 5.7684 | 3.4464 | 3.7272 |
|  |  | mixed5 | abiotic resistance + constitutive expression | 41.9662 | 23.8645 | 26.0536 |
|  |  | mixed6 | dual-resistance + inductive expression | 32.0792 | 21.9203 | 23.1488 |
| ssp585_Mid | 2041-2060 | Highest risk | abiotic resistance + inductive expression | 84.5585 | 65.144 | 67.4919 |
|  |  | Lowest risk | dual-resistance + constitutive expression | 5.9091 | 3.4194 | 3.7205 |
|  |  | mixed7 | abiotic resistance + constitutive expression | 43.3124 | 23.6563 | 26.0334 |
|  |  | mixed8 | dual-resistance + inductive expression | 32.5974 | 21.937 | 23.2262 |
| ssp126_Long | 2061-2080 | Highest risk | abiotic resistance + inductive expression | 83.465 | 64.4431 | 66.7434 |
|  |  | Lowest risk | dual-resistance + constitutive expression | 5.5186 | 3.3803 | 3.6389 |
|  |  | mixed7 | abiotic resistance + constitutive expression | 41.2856 | 23.9828 | 26.0753 |
|  |  | mixed8 | dual-resistance + inductive expression | 31.8097 | 21.8357 | 23.0419 |
| ssp245_Long | 2061-2080 | Highest risk | abiotic resistance + inductive expression | 84.4116 | 65.0877 | 67.4246 |
|  |  | Lowest risk | dual-resistance + constitutive expression | 5.9083 | 3.37 | 3.677 |
|  |  | mixed9 | abiotic resistance + constitutive expression | 43.2832 | 23.5879 | 25.9697 |
|  |  | mixed10 | dual-resistance + inductive expression | 33.035 | 21.9847 | 23.321 |
| ssp585_Long | 2061-2080 | Highest risk | abiotic resistance + inductive expression | 85.0318 | 66.7451 | 68.9566 |
|  |  | Lowest risk | dual-resistance + constitutive expression | 6.2432 | 3.3988 | 3.7428 |
|  |  | mixed9 | abiotic resistance + constitutive expression | 45.9672 | 23.5613 | 26.2709 |
|  |  | mixed10 | dual-resistance + inductive expression | 34.1842 | 22.3215 | 23.7561 |
| ssp126_End | 2081-2100 | Highest risk | abiotic resistance + inductive expression | 83.5008 | 64.4457 | 66.7501 |
|  |  | Lowest risk | dual-resistance + constitutive expression | 5.6703 | 3.4824 | 3.747 |
|  |  | mixed11 | abiotic resistance + constitutive expression | 41.0208 | 24.0615 | 26.1125 |
|  |  | mixed12 | dual-resistance + inductive expression | 31.9114 | 21.919 | 23.1274 |
| ssp245_End | 2081-2100 | Highest risk | abiotic resistance + inductive expression | 84.752 | 65.5032 | 67.831 |
|  |  | Lowest risk | dual-resistance + constitutive expression | 6.0196 | 3.4033 | 3.7197 |
|  |  | mixed11 | abiotic resistance + constitutive expression | 44.1068 | 23.4429 | 25.9418 |
|  |  | mixed12 | dual-resistance + inductive expression | 33.1617 | 21.8507 | 23.2186 |
| ssp585_End | 2081-2100 | Highest risk | abiotic resistance + inductive expression | 84.6933 | 69.5127 | 71.3485 |
|  |  | Lowest risk | dual-resistance + constitutive expression | 6.434 | 3.5826 | 3.9275 |
|  |  | mixed13 | abiotic resistance + constitutive expression | 47.6681 | 24.6394 | 27.4244 |
|  |  | mixed14 | dual-resistance + inductive expression | 34.4131 | 23.114 | 24.4804 |

**Supplementary Table 18**

**High-Risk area Proportion (HRP) for extrapolated temperature-extent in five periods.** HRP (High‑Risk Proportion) is defined as the area‑weighted posterior probability that lnRR  ≥ 0, representing the expected fraction of vegetated land (MODIS IGBP classes 1‑12 and 14) with high ecological risk of escaped GMPs. The Bayesian model constructed in this study was based on History-period data (1970-2000). Future climate inputs are from five CMIP6 models (ISIMIP3b subset). Four future periods (Near: 2021‑2040, Mid: 2041‑2060, Long: 2061‑2080, End: 2081‑2100) are considered under three Shared Socioeconomic Pathways (SSPs): SSP1‑2.6 (low emissions), SSP2‑4.5 (intermediate emissions), and SSP5‑8.5 (high emissions). Predictions allow full temperature extrapolation (i.e., no clipping to the training range of 1.7‑26.8 °C). dual-resistance: abiotic & biotic resistance.

|  |  |  |  | **High-Risk area Proportion (%)** | | |
| --- | --- | --- | --- | --- | --- | --- |
| Period | Time | Scenario | Description | cropland | natural | Total vegetation |
| History | 1970-2000 | Highest risk | abiotic resistance + inductive expression | 78.5194 | 59.88551052 | 62.137912 |
|  |  | Lowest risk | dual-resistance + constitutive expression | 4.86162 | 3.543185889 | 3.702554 |
|  |  | mixed1 | abiotic resistance + constitutive expression | 34.4343 | 25.19441185 | 26.311302 |
|  |  | mixed2 | dual-resistance + inductive expression | 29.4878 | 22.20406854 | 23.084504 |
| ssp126_Near | 2021-2040 | Highest risk | abiotic resistance + inductive expression | 80.0142 | 59.4953 | 61.9767 |
|  |  | Lowest risk | dual-resistance + constitutive expression | 5.4892 | 3.4674 | 3.7119 |
|  |  | mixed1 | abiotic resistance + constitutive expression | 39.2181 | 23.8691 | 25.7253 |
|  |  | mixed2 | dual-resistance + inductive expression | 30.5804 | 21.0663 | 22.2169 |
| ssp245_Near | 2021-2040 | Highest risk | abiotic resistance + inductive expression | 80.4779 | 59.5715 | 62.0998 |
|  |  | Lowest risk | dual-resistance + constitutive expression | 5.5512 | 3.4698 | 3.7215 |
|  |  | mixed3 | abiotic resistance + constitutive expression | 39.6572 | 24.0052 | 25.898 |
|  |  | mixed4 | dual-resistance + inductive expression | 30.7028 | 20.9059 | 22.0906 |
| ssp585_Near | 2021-2040 | Highest risk | abiotic resistance + inductive expression | 80.709 | 59.5554 | 62.1136 |
|  |  | Lowest risk | dual-resistance + constitutive expression | 5.5371 | 3.4479 | 3.7006 |
|  |  | mixed3 | abiotic resistance + constitutive expression | 40.0024 | 23.8611 | 25.8131 |
|  |  | mixed4 | dual-resistance + inductive expression | 30.9415 | 21.048 | 22.2445 |
| ssp126_Mid | 2041-2060 | Highest risk | abiotic resistance + inductive expression | 79.6431 | 59.3649 | 61.8172 |
|  |  | Lowest risk | dual-resistance + constitutive expression | 5.5837 | 3.4154 | 3.6776 |
|  |  | mixed5 | abiotic resistance + constitutive expression | 40.2706 | 23.6522 | 25.6619 |
|  |  | mixed6 | dual-resistance + inductive expression | 30.8342 | 20.7541 | 21.9731 |
| ssp245_Mid | 2041-2060 | Highest risk | abiotic resistance + inductive expression | 79.5781 | 58.9614 | 61.4547 |
|  |  | Lowest risk | dual-resistance + constitutive expression | 5.6014 | 3.277 | 3.5581 |
|  |  | mixed5 | abiotic resistance + constitutive expression | 41.1507 | 23.2971 | 25.4562 |
|  |  | mixed6 | dual-resistance + inductive expression | 31.0452 | 20.5098 | 21.7838 |
| ssp585_Mid | 2041-2060 | Highest risk | abiotic resistance + inductive expression | 78.8565 | 58.3142 | 60.7984 |
|  |  | Lowest risk | dual-resistance + constitutive expression | 5.7792 | 3.2871 | 3.5885 |
|  |  | mixed7 | abiotic resistance + constitutive expression | 42.498 | 22.9079 | 25.277 |
|  |  | mixed8 | dual-resistance + inductive expression | 31.0742 | 20.2129 | 21.5263 |
| ssp126_Long | 2061-2080 | Highest risk | abiotic resistance + inductive expression | 79.2392 | 59.1845 | 61.6098 |
|  |  | Lowest risk | dual-resistance + constitutive expression | 5.5463 | 3.3715 | 3.6345 |
|  |  | mixed7 | abiotic resistance + constitutive expression | 40.562 | 23.3219 | 25.4068 |
|  |  | mixed8 | dual-resistance + inductive expression | 31.0267 | 20.816 | 22.0508 |
| ssp245_Long | 2061-2080 | Highest risk | abiotic resistance + inductive expression | 78.374 | 58.226 | 60.6625 |
|  |  | Lowest risk | dual-resistance + constitutive expression | 5.7731 | 3.2421 | 3.5482 |
|  |  | mixed9 | abiotic resistance + constitutive expression | 42.391 | 22.7978 | 25.1673 |
|  |  | mixed10 | dual-resistance + inductive expression | 31.4359 | 20.2341 | 21.5888 |
| ssp585_Long | 2061-2080 | Highest risk | abiotic resistance + inductive expression | 75.8794 | 57.2856 | 59.5342 |
|  |  | Lowest risk | dual-resistance + constitutive expression | 5.8934 | 3.1417 | 3.4745 |
|  |  | mixed9 | abiotic resistance + constitutive expression | 44.6332 | 22.2396 | 24.9477 |
|  |  | mixed10 | dual-resistance + inductive expression | 31.2039 | 19.7188 | 21.1077 |
| ssp126_End | 2081-2100 | Highest risk | abiotic resistance + inductive expression | 79.3323 | 59.3145 | 61.7353 |
|  |  | Lowest risk | dual-resistance + constitutive expression | 5.4913 | 3.3626 | 3.62 |
|  |  | mixed11 | abiotic resistance + constitutive expression | 40.4595 | 23.6357 | 25.6703 |
|  |  | mixed12 | dual-resistance + inductive expression | 30.7181 | 20.7021 | 21.9134 |
| ssp245_End | 2081-2100 | Highest risk | abiotic resistance + inductive expression | 77.4613 | 57.7698 | 60.1512 |
|  |  | Lowest risk | dual-resistance + constitutive expression | 5.8595 | 3.165 | 3.4909 |
|  |  | mixed11 | abiotic resistance + constitutive expression | 43.0262 | 22.408 | 24.9014 |
|  |  | mixed12 | dual-resistance + inductive expression | 31.2819 | 19.9605 | 21.3297 |
| ssp585_End | 2081-2100 | Highest risk | abiotic resistance + inductive expression | 71.5432 | 56.4989 | 58.3182 |
|  |  | Lowest risk | dual-resistance + constitutive expression | 6.0506 | 3.2676 | 3.6042 |
|  |  | mixed13 | abiotic resistance + constitutive expression | 45.7821 | 22.8013 | 25.5804 |
|  |  | mixed14 | dual-resistance + inductive expression | 31.17 | 20.1071 | 21.445 |

**Supplementary Table 19**

**List of publications excluded from the dataset of this study with the reasons for exclusion.**

| **No.** | **Publication** |
| --- | --- |
| **Reason: the fitness effects involve mixed results (arising from differences in measurement time-point, gene expression levels, individuals, condition setting, etc.) and lacks mean assessment.** | |
| 1 | Ceccoli, R.D. et al. (2012) Flavodoxin displays dose‑dependent effects on photosynthesis and stress tolerance when expressed in transgenic tobacco plants. Planta, 236, 1447‑1458. |
| 2 | Brown, R.H. et al. (2013) Unintended consequences: high phosphinothricin acetyltransferase activity related to reduced fitness in barley. In Vitro Cellular & Developmental Biology‑Plant, 49, 240‑247. |
| 3 | Bhatnagar-Mathur, P. et al. (2007) Stress-inducible expression of At DREB1A in transgenic peanut (Arachis hypogaea L.) increases transpiration efficiency under water-limiting conditions. Plant Cell Reports, 26, 2071-2082. |
| 4 | Behnke, K. et al. (2012) Isoprene emission-free poplars--a chance to reduce the impact from poplar plantations on the atmosphere. The New Phytologist, 194, 70-82. |
| 5 | Chen, S. et al. (2021) Knockout of the entire family of AITR genes in Arabidopsis leads to enhanced drought and salinity tolerance without fitness costs. BMC Plant Biology, 21, 137. |
| 6 | Cipollini, D. (2007) Consequences of the overproduction of methyl jasmonate on seed production, tolerance to defoliation and competitive effect and response of Arabidopsis thaliana. New Phytologist, 173, 146-153. |
| 7 | Cipollini, D. (2010) Constitutive expression of methyl jasmonate-inducible responses delays reproduction and constrains fitness responses to nutrients in Arabidopsis thaliana. Evolutionary Ecology, 24, 59-68. |
| 8 | Cosseboom, S.D. et al. (2023) CRISPR-enabled investigation of fitness costs associated with the E198A mutation in β-tubulin of Colletotrichum siamense. Frontiers in Plant Science, 14. |
| 9 | Fladung, M. et al. (1993) Constitutive or light-regulated expression of the rolC gene in transgenic potato plants has different effects on yield attributes and tuber carbohydrate composition. Plant Molecular Biology, 23, 749-757. |
| 10 | Fu, J. et al. (2018) Fitness Cost of Transgenic cry1Ab/c Rice Under Saline-Alkaline Soil Condition. Frontiers in Plant Science, 9. |
| 11 | Fu, J. et al. (2021) Fitness of Insect-resistant transgenic rice T1C-19 under four growing conditions combining land use and weed competition. GM Crops & Food, 12, 328-341. |
| 12 | Fu, J. and Liu, B. (2022) Individual and combined effects of land use and weeds on Cry1Ab/c protein expression and yield of transgenic cry1Ab/c rice. GM Crops & Food, 13, 156-170. |
| 13 | Fu, J. and Liu, B. (2020) Enhanced yield performance of transgenic cry1C* rice in saline-alkaline soil. GM Crops & Food, 11, 97-112. |
| 14 | Heidel, A.J. and Dong, X. (2006) Fitness benefits of systemic acquired resistance during Hyaloperonospora parasitica infection in Arabidopsis thaliana. Genetics, 173, 1621-1628. |
| 15 | Hyun, T.K. et al. (2015) The Arabidopsis PLAT domain protein1 promotes abiotic stress tolerance and growth in tobacco. Transgenic Research, 24, 651-663. |
| 16 | Jiang, X.-Q. et al. (2021) Increased Longevity and Dormancy of Soil-Buried Seeds from Advanced Crop-Wild Rice Hybrids Overexpressing the EPSPS Transgene. Biology, 10, 562. |
| 17 | Jørgensen, R. et al. (1997) Introgression of crop genes from oilseed rape (Brassica napus) to related wild species-an avenue for the escape of engineered genes. 8th International Symposium on the Biosafety of Genetically Modified Organisms, 211-218. |
| 18 | Kang, I.-H. et al. (2023) A novel mechanism of herbicide action through disruption of pyrimidine biosynthesis. Proceedings of the National Academy of Sciences, 120, e2313197120. |
| 19 | Khan, M.H. et al. (2021) Targeted mutagenesis of EOD3 gene in Brassica napus L. regulates seed production. Journal of Cellular Physiology, 236, 1996-2007. |
| 20 | Koller, T. et al. (2019) Field grown transgenic Pm3e wheat lines show powdery mildew resistance and no fitness costs associated with high transgene expression. Transgenic Research, 28, 9-20. |
| 21 | Li, Z. et al. (2019) Overexpression of Arabidopsis Nucleotide-Binding and Leucine-Rich Repeat Genes RPS2 and RPM1(D505V) Confers Broad-Spectrum Disease Resistance in Rice. Frontiers in Plant Science, 10. |
| 22 | Li, R. et al. (2024) Less is more: CRISPR/Cas9-based mutations in DND1 gene enhance tomato resistance to powdery mildew with low fitness costs. BMC Plant Biology, 24, 763. |
| 23 | Li, S. et al. (2009) Functional analysis of an Arabidopsis transcription factor WRKY25 in heat stress. Plant Cell Reports, 28, 683-693. |
| 24 | Linder, C.R. and Schmitt, J. (1995) Potential persistence of escaped transgenes: performance of transgenic, oil-modified Brassica seeds and seedlings. Ecological Applications, 5, 1056-1068. |
| 25 | Schuman, M.C. et al. (2014) Ectopic terpene synthase expression enhances sesquiterpene emission in Nicotiana attenuata without altering defense or development of transgenic plants or neighbors. Plant Physiology, 166, 779-797. |
| 26 | Xu, J. et al. (2015) Transgenic Arabidopsis plants expressing tomato glutathione S-transferase showed enhanced resistance to salt and drought stress. PLoS ONE, 10, e0136960. |
| 27 | Yang, C. et al. (2014) Segregation distortion affected by transgenes in early generations of rice crop-weed hybrid progeny: Implications for assessing potential evolutionary impacts from transgene flow into wild relatives. Journal of Systematics and Evolution, 52, 466-476. |
| 28 | Song, X. et al. (2010) Potential gene flow of two herbicide-tolerant transgenes from oilseed rape to wild B. juncea var. gracilis. Theoretical and Applied Genetics, 120, 1501-1510. |
| 29 | Sun, X. et al. (2015) A 14-3-3 Family Protein from Wild Soybean (Glycine soja) Regulates ABA Sensitivity in Arabidopsis. PLoS ONE, 10, e0146163. |
| **Reason: without a stress-free condition setting.** | |
| 1 | Pandolfo, C.E. et al. (2018) Transgene escape and persistence in an agroecosystem: the case of glyphosate-resistant *Brassica rapa* L. in central Argentina. *Environmental Science and Pollution Research*, 25, 6251-6264. |
| 2 | Al-Ahmad, H. et al. (2004) Tandem constructs to mitigate transgene persistence: tobacco as a model. *Molecular Ecology*, 13, 697-710. |
| 3 | Al-Ahmad, H. and Gressel, J. (2006) Mitigation using a tandem construct containing a selectively unfit gene precludes establishment of *Brassica napus* transgenes in hybrids and backcrosses with weedy *Brassica rapa*. *Plant Biotechnology Journal*, 4, 23-33. |
| 4 | Benevenuto, R.F. et al. (2021) Proteomic profile of glyphosate-resistant soybean under combined herbicide and drought stress conditions. *Plants*, 10, 2381. |
| 5 | Bigelow, P.J. et al. (2018) Influence of intergenotypic competition on multigenerational persistence of abiotic stress resistance transgenes in populations of *Arabidopsis thaliana*. *Evolutionary Applications*, 11, 950-962. |
| 6 | Chen, L.-Y. et al. (2006) Effects of insect-resistance transgenes on fecundity in rice (*Oryza sativa*, Poaceae): a test for underlying costs. *American Journal of Botany*, 93, 94-101. |
| 7 | Cowgill, S.E. et al. (2002) Transgenic potatoes with enhanced levels of nematode resistance do not have altered susceptibility to nontarget aphids. *Molecular Ecology*, 11, 821-827. |
| 8 | Di, K. et al. (2009) Fitness and maternal effects in hybrids formed between transgenic oilseed rape (*Brassica napus* L.) and wild brown mustard [*B. juncea* (L.) Czern et Coss.] in the field. *Pest Management Science*, 65, 753-760. |
| 9 | Dixit, P. et al. (2011) Glutathione transferase from *Trichoderma virens* enhances cadmium tolerance without enhancing its accumulation in transgenic *Nicotiana tabacum*. *PLoS ONE*, 6, e16360. |
| 10 | Dong, S.s. et al. (2017) Persistence of transgenes in wild rice populations depends on the interaction between genetic background of recipients and environmental conditions. *Annals of Applied Biology*, 171, 202-213. |
| 11 | Fen, L.H. et al. (2002) Maize transformation of cry1Ac3 gene and insect resistance of their transgenic plants. *Journal of Integrative Plant Biology*, 44, 684. |
| 12 | Fillatti, J.J. et al. (1987) Efficient transfer of a glyphosate tolerance gene into tomato using a binary *Agrobacterium tumefaciens* vector. *Bio/technology*, 5, 726-730. |
| 13 | Foresi, N. et al. (2015) Expression of the tetrahydrofolate-dependent nitric oxide synthase from the green alga *Ostreococcus tauri* increases tolerance to abiotic stresses and influences stomatal development in *Arabidopsis*. *The Plant Journal*, 82, 806-821. |
| 14 | FU, J.-m. et al. (2015) Ecological Fitness of Transgenic cry1Ab/c Rice in Nutrient-Deficient Soils. *Journal of Ecology and Rural Environment*, 31, 528-533. |
| 15 | Gaxiola, R.A. et al. (2001) Drought- and salt-tolerant plants result from overexpression of the AVP1 H+ -pump. *Proceedings of the National Academy of Sciences*, 98, 11444-11449. |
| 16 | Hails, R.S. et al. (1997) Burial and seed survival in *Brassica napus* subsp. *oleifera* and *Sinapis arvensis* including a comparison of transgenic and non-transgenic lines of the crop. *Proceedings of the Royal Society B: Biological Sciences*. |
| 17 | Harris, C.J. et al. (2013) Stepwise artificial evolution of a plant disease resistance gene. *Proceedings of the National Academy of Sciences*, 110, 21189-21194. |
| 18 | Harth, J.E. et al. (2016) Effects of virus infection on pollen production and pollen performance: Implications for the spread of resistance alleles. *American Journal of Botany*, 103, 577-583. |
| 19 | Harth, J.E. et al. (2018) Zucchini Yellow Mosaic Virus Infection Limits Establishment and Severity of Powdery Mildew in Wild Populations of *Cucurbita pepo*. *Frontiers in Plant Science*, 9. |
| 20 | Hui Xia et al. (2010) Yield benefit and underlying cost of insect-resistance transgenic rice: Implication in breeding and deploying transgenic crops. *Field Crops Research*, 118, 215-220. |
| 21 | Igarashi, D. et al. (2012) The peptide growth factor, phytosulfokine, attenuates pattern-triggered immunity. *The Plant Journal*, 71, 194-204. |
| 22 | Jin, C.Y. et al. (2013) Fitness cost and competitive ability of transgenic herbicide-tolerant rice expressing a protoporphyrinogen oxidase gene. *Journal of Ecology and Environment*, 36, 39-47. |
| 23 | Johannessen, M.M. et al. (2006) Competition affects gene flow from oilseed rape (female symbol) to *Brassica rapa* (male symbol). *Heredity*, 96, 360-367. |
| 24 | Jørgensen, R.B. et al. (2009) The variability of processes involved in transgene dispersal-case studies from *Brassica* and related genera. *Environmental Science and Pollution Research*, 16, 389-395. |
| 25 | Ju, Y. et al. (2017) Overexpression of OsHSP18.0-CI enhances resistance to bacterial leaf streak in rice. *Rice*, 10, 1-11. |
| 26 | Kalinina, O. et al. (2011) Competitive Performance of Transgenic Wheat Resistant to Powdery Mildew. *PLoS ONE*, 6, e28091. |
| 27 | Kalinina, O. et al. (2015) Persistence of seeds, seedlings and plants, performance of transgenic wheat in weed communities in the field and effects on fallow weed diversity. *Perspectives in Plant Ecology, Evolution and Systematics*, 17, 421-433. |
| 28 | Karavangeli, M. et al. (2005) Development of transgenic tobacco plants overexpressing maize glutathione S-transferase I for chloroacetanilide herbicides phytoremediation. *Biomolecular Engineering*, 22, 121-128. |
| 29 | Letourneau, D.K. and Hagen, J.A. (2009) Plant fitness assessment for wild relatives of insect resistant crops. *Environmental Biosafety Research*, 8, 45-55. |
| 30 | Li, S. et al. (2022) Elevated CO2 and high endogenous ABA level alleviate PEG-induced short-term osmotic stress in tomato plants. *Environmental and Experimental Botany*, 194, 104763. |
| 31 | Liu, L. et al. (2020) Expression of Bt Protein in Transgenic Bt Cotton Plants and Ecological Fitness of These Plants in Different Habitats. *Frontiers in Plant Science*, 11. |
| 32 | Liu, Y. et al. (2015) Characterization of competitive interactions in the coexistence of Bt-transgenic and conventional rice. *BMC Biotechnology*, 15, 27. |
| 33 | Liu, M. et al. (2019) Inducible overexpression of Ideal Plant Architecture1 improves both yield and disease resistance in rice. *Nature Plants*, 5, 389-400. |
| 34 | Liu, Y. et al. (2015) The presence of Bt-transgenic oilseed rape in wild mustard populations affects plant growth. *Transgenic Research*, 24, 1043-1053. |
| 35 | Liu, Y. et al. (2013) Spread of introgressed insect-resistance genes in wild populations of *Brassica juncea*: a simulated in-vivo approach. *Transgenic Research*, 22, 747-756. |
| 36 | Londo, J.P. et al. (2010) Glyphosate drift promotes changes in fitness and transgene gene flow in canola (*Brassica napus*) and hybrids. *Annals of Botany*, 106, 957-965. |
| 37 | Lu, H.-P. et al. (2017) CRISPR-S: an active interference element for a rapid and inexpensive selection of genome-edited, transgene-free rice plants. *Plant Biotechnology Journal*, 15, 1371-1373. |
| 38 | Luo, J.Y. et al. (2016) Ecological fitness of transgenic GAFP cotton and its effects on the field insect community. *Journal of Applied Ecology*, 27, 3675-3681. |
| 39 | Mannaa, M. et al. (2024) Exploring the comparative genome of rice pathogen *Burkholderia plantarii*: unveiling virulence, fitness traits, and a potential type III secretion system effector. *Frontiers in Plant Science*, 15. |
| 40 | Metz, T.D. et al. (1995) Transgenic broccoli expressing a *Bacillus thuringiensis* insecticidal crystal protein: implications for pest resistance management strategies. *Molecular Breeding*, 1, 309-317. |
| 41 | Millwood, R.J. (2011) Consequences of gene flow and transgene introgression in hybrids between transgenic *Brassica napus* and its weedy wild relative *Brassica rapa*. |
| 42 | Nawaz, G. and Kang, H. (2019) Rice OsRH58, a chloroplast DEAD-box RNA helicase, improves salt or drought stress tolerance in *Arabidopsis* by affecting chloroplast translation. *BMC Plant Biology*, 19, 1-11. |
| 43 | Panda, S. et al. (2025) Jasmonate Responsive SlnsLTP Confers Resistance Against *Botrytis cinerea* and *Verticillium dahliae* in Tomato. *Journal of Plant Growth Regulation*. |
| 44 | Peiró, A. et al. (2014) The movement protein (NSm) of *Tomato spotted wilt virus* is the avirulence determinant in the tomato Sw-5 gene-based resistance. *Molecular Plant Pathology*, 15, 802-813. |
| 45 | Pessel, D. et al. (2001) Persistence of oilseed rape (*Brassica napus* L.) outside of cultivated fields. *Theoretical and Applied Genetics*, 102, 841-846. |
| 46 | Qi, M. et al. (2019) QQS orphan gene and its interactor NF-YC4 reduce susceptibility to pathogens and pests. *Plant Biotechnology Journal*, 17, 252-263. |
| 47 | Qian, Q. et al. (2014) Enhanced resistance to blast fungus in rice (*Oryza sativa* L.) by expressing the ribosome-inactivating protein alpha-momorcharin. *Plant Science*, 217, 1-7. |
| 48 | Qiao, D. et al. (2023) Optimized prime editing efficiently generates heritable mutations in maize. *Journal of Integrative Plant Biology*, 65, 900-906. |
| 49 | Qin, X. and Zeevaart, J.A.D. (2002) Overexpression of a 9-cis-epoxycarotenoid dioxygenase gene in *Nicotiana plumbaginifolia* increases abscisic acid and phaseic acid levels and enhances drought tolerance. *Plant Physiology*, 128, 544-551. |
| 50 | Qiu, D. et al. (2008) Rice gene network inferred from expression profiling of plants overexpressing OsWRKY13, a positive regulator of disease resistance. *Molecular Plant*, 1, 538-551. |
| 51 | Rahnamaeian, M. and Vilcinskas, A. (2012) Defense gene expression is potentiated in transgenic barley expressing antifungal peptide metchnikowin throughout powdery mildew challenge. *Journal of Plant Research*, 125, 115-124. |
| 52 | Ramachandran, S. et al. (2000) Intraspecific Competition of an Insect-Resistant Transgenic Canola in Seed Mixtures. *Agronomy Journal*, 92, 368-374. |
| 53 | Rees, J.D. et al. (2009) Relative contributions of nine genes in the pathway of histidine biosynthesis to the control of free histidine concentrations in *Arabidopsis thaliana*. *Plant Biotechnology Journal*, 7, 499-511. |
| 54 | Santamaria, M.E. et al. (2018) Overexpression of HvIcy6 in barley enhances resistance against *Tetranychus urticae* and entails partial transcriptomic reprogramming. *International Journal of Molecular Sciences*, 19, 697. |
| 55 | Schwachtje, J. et al. (2008) Reverse genetics in ecological research. *PLoS ONE*, 3, e1543. |
| 56 | Tian, B. et al. (2019) Host-derived gene silencing of parasite fitness genes improves resistance to soybean cyst nematodes in stable transgenic soybean. *Theoretical and Applied Genetics*, 132, 2651-2662. |
| 57 | Wang, G.-L. et al. (1996) The cloned gene, Xa21, confers resistance to multiple *Xanthomonas oryzae* pv. *oryzae* isolates in transgenic plants. *Molecular Plant-Microbe Interactions*, 9, 850-855. |
| 58 | Xia, H. et al. (2011) Enhanced yield performance of Bt rice under target-insect attacks: implications for field insect management. *Transgenic Research*, 20, 655-664. |
| 59 | Yang, X. et al. (2015) Efficacy of insect-resistance Bt/CpTI transgenes in F5-F7 generations of rice crop-weed hybrid progeny: implications for assessing ecological impact of transgene flow. *Science Bulletin*, 60, 1563-1571. |
| 60 | Yang, X. et al. (2012) Limited fitness advantages of crop-weed hybrid progeny containing insect-resistant transgenes (Bt/CpTI) in transgenic rice field. *PLoS ONE*, 7, e41220. |
| 61 | Zhang, F. et al. (2012) Differences in Ecological Fitness Between Bt Transgenic Rice and Regular Rice Under Different Insect-infestation Pressures. *Chinese Journal of Applied Environmental Biology*, 18, 35. |
| 62 | Simard, M.-J. et al. (2025) Competitive fitness of bird rape mustard (*Brassica rapa* L.) that integrated transgenes conferring glyphosate resistance in commercial fields. *Weed Research*, 65, e70003. |
| 63 | Srinivasan, T. (2021) Overexpression of Trypsin Protease Inhibitor in Transgenic Tobacco Plants Exhibit Oxidative Stress Tolerance and Fungal Resistance. *Annals of the Romanian Society for Cell Biology*, 25, 7629-7634. |
| 64 | Stewart, C.N. et al. (1997) Increased fitness of transgenic insecticidal rapeseed under insect selection pressure. *Molecular Ecology*, 6, 773-779. |
| 65 | Sun, K. et al. (2016) Down-regulation of *Arabidopsis* DND1 orthologs in potato and tomato leads to broad-spectrum resistance to late blight and powdery mildew. *Transgenic Research*, 25, 123-138. |
| 66 | Turrà, D. et al. (2020) Heterologous Expression of PKPI and Pin1 Proteinase Inhibitors Enhances Plant Fitness and Broad-Spectrum Resistance to Biotic Threats. *Frontiers in Plant Science*, 11, 461. |
| 67 | Wei, J. et al. (2011) Ecological trade-offs between jasmonic acid-dependent direct and indirect plant defences in tritrophic interactions. *New Phytologist*, 189, 557-567. |
| 68 | Wu, C. et al. Limited fitness costs of herbicide-resistance traits in *Amaranthus tuberculatus* facilitate resistance evolution. |
| 69 | Yanniccari, M. et al. (2016) Glyphosate resistance in perennial ryegrass (*Lolium perenne* L.) is associated with a fitness penalty. *Weed Science*, 64, 71-79. |
| 70 | Zhu, J.-Q. et al. (2012) Improvement of pest resistance in transgenic tobacco plants expressing dsRNA of an insect-associated gene EcR. *PLoS ONE*, 7, e38572. |
| **Reason: without increasing resistance (abiotic or biotic) in HR.** | |
| 1 | Ahlawat, Y.K. et al. (2024) Heterologous expression of *Arabidopsis* laccase2, laccase4 and peroxidase52 driven under developing xylem specific promoter DX15 improves saccharification in populus. *Biotechnology for Biofuels and Bioproducts*, 17, 5. |
| 2 | Aly, R. et al. (2019) Use of a visible reporter marker- myb-related gene in crop plants to minimize herbicide usage against weeds. *Plant Signaling & Behavior*, 14, e1581558. |
| 3 | Anstead, J.A. et al. (2012) *Arabidopsis* P-protein filament formation requires both AtSEOR1 and AtSEOR2. *Plant and Cell Physiology*, 53, 1033-1042. |
| 4 | Arriola, P.E. and Ellstrand, N.C. (1997) Fitness of interspecific hybrids in the genus *Sorghum*: persistence of crop genes in wild populations. *Ecological Applications*, 7, 512-518. |
| 5 | Augustin, J.M. et al. (2019) Field performance of terpene-producing *Camelina sativa*. *Industrial Crops and Products*, 136, 50-58. |
| 6 | Austin, S. et al. (1995) Production and field performance of transgenic alfalfa (*Medicago sativa* L.) expressing alpha-amylase and manganese-dependent lignin peroxidase. 381-393. |
| 7 | Baack, E.J. et al. (2008) Selection on domestication traits and quantitative trait loci in crop-wild sunflower hybrids. *Molecular Ecology*, 17, 666-677. |
| 8 | Baldwin, I.T. (1996) Methyl jasmonate-induced nicotine production in *Nicotiana attenuata*: inducing defenses in the field without wounding. 213-220. |
| 9 | Baldwin, I.T. (1999) Inducible nicotine production in native *Nicotiana* as an example of adaptive phenotypic plasticity. *Journal of Chemical Ecology*, 25, 3-30. |
| 10 | Baxter, H.L. et al. (2015) Field evaluation of transgenic switchgrass plants overexpressing PvMYB4 for reduced biomass recalcitrance. *BioEnergy Research*, 8, 910-921. |
| 11 | Beltramino, M. et al. (2018) Robust increase of leaf size by *Arabidopsis thaliana* GRF3-like transcription factors under different growth conditions. *Scientific Reports*, 8, 13447. |
| 12 | Benedetti, M. et al. (2020) Expression of a hyperthermophilic cellobiohydrolase in transgenic *Nicotiana tabacum* by protein storage vacuole targeting. *Plants*, 9, 1799. |
| 13 | Bhatia, R. et al. (2023) Transgenic ZmMYB167 *Miscanthus sinensis* with increased lignin to boost bioenergy generation for the bioeconomy. *Biotechnology for Biofuels and Bioproducts*, 16, 29. |
| 14 | Bovy, A.G. et al. (1999) Heterologous expression of the *Arabidopsis* etr1-1 allele inhibits the senescence of carnation flowers. *Molecular Breeding*, 5, 301-308. |
| 15 | Buyel, J.F. and Fischer, R. (2012) Predictive models for transient protein expression in tobacco (*Nicotiana tabacum* L.) can optimize process time, yield, and downstream costs. *Biotechnology and Bioengineering*, 109, 2575-2588. |
| 16 | Callahan, H.S. et al. (2005) Plasticity genes and plasticity costs: a new approach using an *Arabidopsis* recombinant inbred population. *New Phytologist*, 166, 129-139. |
| 17 | Casler, M.D. et al. (2002) Genetic modification of lignin concentration affects fitness of perennial herbaceous plants. *Theoretical and Applied Genetics*, 104, 127-131. |
| 18 | Casler, M.D. and Jung, H.-J.G. (2021) Agronomic fitness of three temperate forage grasses divergently selected for lignin concentration or ferulate cross-linking. *Euphytica*, 217. |
| 19 | Chahardoli, M. et al. (2018) Recombinant expression of LFchimera antimicrobial peptide in a plant-based expression system and its antimicrobial activity against clinical and phytopathogenic bacteria. *Biotechnology & Biotechnological Equipment*, 32, 714-723. |
| 20 | Chehab, E.W. et al. (2008) Distinct roles of jasmonates and aldehydes in plant-defense responses. *PLoS ONE*, 3, e1904. |
| 21 | Chen, L. et al. (2003) Improved forage digestibility of tall fescue (*Festuca arundinacea*) by transgenic down-regulation of cinnamyl alcohol dehydrogenase. *Plant Biotechnology Journal*, 1, 437-449. |
| 22 | Cheng, M. et al. (1996) Production of fertile transgenic peanut (*Arachis hypogaea* L.) plants using *Agrobacterium tumefaciens*. *Plant Cell Reports*, 15, 653-657. |
| 23 | Cheng, Q. et al. (2022) Oilseed rape MPK1 mediates reactive oxygen species-dependent cell death and jasmonic acid-induced leaf senescence. *Environmental and Experimental Botany*, 202, 105028. |
| 24 | Constabel, C.P. et al. (1993) Transgenic potato plants overexpressing the pathogenesis-related STH-2 gene show unaltered susceptibility to *Phytophthora infestans* and potato virus X. *Plant Molecular Biology*, 22, 775-782. |
| 25 | Desbiez, C. et al. (2003) Increase in *Zucchini yellow mosaic virus* symptom severity in tolerant zucchini cultivars is related to a point mutation in P3 protein and is associated with a loss of relative fitness on susceptible plants. *Phytopathology*, 93, 1478-1484. |
| 26 | Ding, B. (2023) The roles of R2R3-MYBs in regulating complex pigmentation patterns in flowers. *Horticultural Plant Journal*, 9, 1067-1078. |
| 27 | Dong, J.-Z. and McHughen, A. (1993) Transgenic flax plants from *Agrobacterium* mediated transformation: incidence of chimeric regenerants and inheritance of transgenic plants. *Plant Science*, 91, 139-148. |
| 28 | Doty, S.L. et al. (2007) Enhanced phytoremediation of volatile environmental pollutants with transgenic trees. *Proceedings of the National Academy of Sciences*, 104, 16816-16821. |
| 29 | Frenkel, M. et al. (2007) Hierarchy amongst photosynthetic acclimation responses for plant fitness. *Physiologia Plantarum*, 129, 455-459. |
| 30 | Ganeteg, U. et al. (2004) Is Each Light-Harvesting Complex Protein Important for Plant Fitness? *Plant Physiology*, 134, 502-509. |
| 31 | Gao, Y. et al. (2024) Optimization and establishment of *Agrobacterium*-mediated transformation in alfalfa (*Medicago sativa* L.) using eGFP as a visual reporter. *Plant Cell, Tissue and Organ Culture (PCTOC)*, 156, 7. |
| 32 | García-Almodóvar, R.C. et al. (2017) Production of transgenic diploid *Cucumis melo* plants. *Plant Cell, Tissue and Organ Culture (PCTOC)*, 130, 323-333. |
| 33 | Garner, C.M. et al. (2021) Opposing functions of the plant TOPLESS gene family during SNC1-mediated autoimmunity. *PLoS Genetics*, 17, e1009026. |
| 34 | Gase, K. et al. (2011) Efficient screening of transgenic plant lines for ecological research. *Molecular Ecology Resources*, 11, 890-902. |
| 35 | Godfree, R.C. et al. (2004) Growth, fecundity and competitive ability of transgenic *Trifolium subterraneum* subsp. *subterraneum* cv. Leura expressing a sunflower seed albumin gene. *Hereditas*, 140, 229-244. |
| 36 | Gutierrez, A. et al. (2011) Persistence of sunflower crop traits and fitness in *Helianthus petiolaris* populations. *Plant Biology*, 13, 821-830. |
| 37 | Harper, B.K. et al. (1999) Green fluorescent protein as a marker for expression of a second gene in transgenic plants. *Nature Biotechnology*, 17, 1125-1129. |
| 38 | Hettwer, K. et al. (2016) Dynamic metabolic changes in seeds and seedlings of *Brassica napus* (oilseed rape) suppressing UGT84A9 reveal plasticity and molecular regulation of the phenylpropanoid pathway. *Phytochemistry*, 124, 46-57. |
| 39 | Hiruma, K. et al. (2016) Root endophyte *Colletotrichum tofieldiae* confers plant fitness benefits that are phosphate status dependent. *Cell*, 165, 464-474. |
| 40 | Hoffman, L.M. et al. (1987) Synthesis and protein body deposition of maize 15-kd zein in transgenic tobacco seeds. *The EMBO Journal*, 6, 3213-3221. |
| 41 | Howe, A. et al. (2006) Rapid and reproducible *Agrobacterium*-mediated transformation of sorghum. *Plant Cell Reports*, 25, 784-791. |
| 42 | Huang, X. et al. (2023) Transgene-free genome editing of vegetatively propagated and perennial plant species in the T0 generation via a co-editing strategy. *Nature Plants*, 9, 1591-1597. |
| 43 | Hühns, M. et al. (2008) Plastid targeting strategies for cyanophycin synthetase to achieve high-level polymer accumulation in *Nicotiana tabacum*. *Plant Biotechnology Journal*, 6, 321-336. |
| 44 | Hühns, M. et al. (2009) Tuber-specific cphA expression to enhance cyanophycin production in potatoes. *Plant Biotechnology Journal*, 7, 883-898. |
| 45 | Janacek, S.H. et al. (2009) Photosynthesis in cells around veins of the C(3) plant *Arabidopsis thaliana* is important for both the shikimate pathway and leaf senescence as well as contributing to plant fitness. *The Plant Journal*, 59, 329-343. |
| 46 | Jänkänpää, H.J. and Jansson, S. (2012) How to Grow Transgenic *Arabidopsis* in the Field. *Transgenic Plants*, 483-494. |
| 47 | Jung, H.-J.G. et al. (2012) Modifying crops to increase cell wall digestibility. *Plant Science*, 185, 65-77. |
| 48 | Jung, H.-J.G. and Ni, W. (1998) Lignification of plant cell walls: impact of genetic manipulation. *Proceedings of the National Academy of Sciences*, 95, 12742-12743. |
| 49 | Kang, J.-H. et al. (2006) Silencing threonine deaminase and JAR4 in *Nicotiana attenuata* impairs jasmonic acid-isoleucine-mediated defenses against *Manduca sexta*. *The Plant Cell*, 18, 3303-3320. |
| 50 | Kessler, D. et al. (2008) Field experiments with transformed plants reveal the sense of floral scents. *Science*, 321, 1200-1202. |
| 51 | Komarnytsky, S. et al. (2006) Cosecretion of protease inhibitor stabilizes antibodies produced by plant roots. *Plant Physiology*, 141, 1185-1193. |
| 52 | Lal, S.K. et al. (2023) Concurrent Overexpression of Rice GS1;1 and GS2 Genes to Enhance the Nitrogen Use Efficiency (NUE) in Transgenic Rice. *Journal of Plant Growth Regulation*, 42, 6699-6720. |
| 53 | Lee, J.-H. et al. (2022) Phytochrome B Conveys Low Ambient Temperature Cues to the Ethylene-Mediated Leaf Senescence in *Arabidopsis*. *Plant & Cell Physiology*, 63, 326-339. |
| 54 | Lee, K.P. et al. (2020) PLANT NATRIURETIC PEPTIDE A and Its Putative Receptor PNP-R2 Antagonize Salicylic Acid-Mediated Signaling and Cell Death. *The Plant Cell*, 32, 2237-2250. |
| 55 | Li, Y. et al. (2011) MOS1 epigenetically regulates the expression of plant Resistance gene SNC1. *Plant Signaling & Behavior*, 6, 434-436. |
| 56 | Li, P. et al. (2014) Multiple FLC haplotypes defined by independent cis-regulatory variation underpin life history diversity in *Arabidopsis thaliana*. *Genes & Development*, 28, 1635-1640. |
| 57 | Li, S. et al. (2024) Field-work reveals a novel function for MAX2 in a native tobacco's high-light adaptions. *Plant, Cell & Environment*, 47, 230-245. |
| 58 | Li, Y.-M. et al. (2024) Expression and function identification of senescence-associated genes under continuous drought treatment in grapevine (*Vitis vinifera* L.) leaves. *Physiology and Molecular Biology of Plants*, 30, 877-891. |
| 59 | Liang, H. et al. (2008) Improved sugar release from lignocellulosic material by introducing a tyrosine-rich cell wall peptide gene in poplar. *Clean–Soil, Air, Water*, 36, 662-668. |
| 60 | Lindgren, L.O. et al. (2003) Seed-specific overexpression of an endogenous *Arabidopsis* phytoene synthase gene results in delayed germination and increased levels of carotenoids, chlorophyll, and abscisic acid. *Plant Physiology*, 132, 779-785. |
| 61 | Liu, J. et al. (2020) Fitness benefit plays a vital role in the retention of the Pi-ta susceptible alleles. *bioRxiv*. |
| 62 | Liu, G.-F. et al. (2018) Implementation of CsLIS/NES in linalool biosynthesis involves transcript splicing regulation in *Camellia sinensis*. *Plant, Cell & Environment*, 41, 176-186. |
| 63 | Luis, R.J. and others (2021) Production of volatile moth sex pheromones in transgenic *Nicotiana benthamiana* plants. *Biodesign Research*. |
| 64 | Machado, R.A.R. et al. (2013) Leaf-herbivore attack reduces carbon reserves and regrowth from the roots via jasmonate and auxin signaling. *New Phytologist*, 200, 1234-1246. |
| 65 | MacLean, A.M. et al. (2014) Phytoplasma Effector SAP54 Hijacks Plant Reproduction by Degrading MADS-box Proteins and Promotes Insect Colonization in a RAD23-Dependent Manner. *PLoS Biology*, 12, e1001835. |
| 66 | Malik, Z.I. (2014) Genes that underlie natural variation in growth rate and flowering time in local accessions of *Arabidopsis thaliana*. |
| 67 | McCabe, M.S. et al. (2001) Effects of PSAG12-IPT gene expression on development and senescence in transgenic lettuce. *Plant Physiology*, 127, 505-516. |
| 68 | Meldau, S. et al. (2012) MAPK-dependent JA and SA signalling in *Nicotiana attenuata* affects plant growth and fitness during competition with conspecifics. *BMC Plant Biology*, 12, 213. |
| 69 | Miao, J. et al. (2020) OsPP2C09, a negative regulatory factor in abscisic acid signalling, plays an essential role in balancing plant growth and drought tolerance in rice. *New Phytologist*, 227, 1417-1433. |
| 70 | Minina, E.A. et al. (2018) Transcriptional stimulation of rate-limiting components of the autophagic pathway improves plant fitness. *Journal of Experimental Botany*, 69, 1415-1432. |
| 71 | Moreno, J.C. et al. (2016) Increased *Nicotiana tabacum* fitness through positive regulation of carotenoid, gibberellin and chlorophyll pathways promoted by *Daucus carota* lycopene beta-cyclase (Dclcyb1) expression. *Journal of Experimental Botany*, 67, 2325-2338. |
| 72 | Munoz-Bertomeu, J. et al. (2006) Up-regulation of 1-deoxy-D-xylulose-5-phosphate synthase enhances production of essential oils in transgenic spike lavender. *Plant Physiology*, 142, 890-900. |
| 73 | Negin, B. and Moshelion, M. (2020) Remember where you came from: ABA insensitivity is epigenetically inherited in mesophyll, but not seeds. *Plant Science*, 295. |
| 74 | Neugart, S. et al. (2020) Ultraviolet-B radiation exposure lowers the antioxidant capacity in the *Arabidopsis thaliana* pdx1.3-1 mutant and leads to glucosinolate biosynthesis alteration in both wild type and mutant. *Photochemical & Photobiological Sciences*, 19, 217-228. |
| 75 | Palmer-Young, E.C. et al. (2015) The sesquiterpenes (E)-ß-farnesene and (E)-alpha-bergamotene quench ozone but fail to protect the wild tobacco *Nicotiana attenuata* from ozone, UVB, and drought stresses. *PLoS ONE*, 10, e0127296. |
| 76 | Pedersen, J.F. et al. (2005) Impact of reduced lignin on plant fitness. *Crop Science*, 45, 812-819. |
| 77 | Pierik, R. et al. (2004) Density-induced plant size reduction and size inequalities in ethylene-sensing and ethylene-insensitive tobacco. *Plant Biology*, 6, 201-205. |
| 78 | Pierik, R. et al. (2003) Ethylene is required in tobacco to successfully compete with proximate neighbours. *Plant, Cell & Environment*, 26, 1229-1234. |
| 79 | Pilate, G. et al. (2002) Field and pulping performances of transgenic trees with altered lignification. *Nature Biotechnology*, 20, 607-612. |
| 80 | Pogorelko, G. et al. (2011) Post-synthetic modification of plant cell walls by expression of microbial hydrolases in the apoplast. *Plant Molecular Biology*, 77, 433-445. |
| 81 | Preis, I. et al. (2018) The modification of cell wall properties by expression of recombinant resilin in transgenic plants. *Molecular Biotechnology*, 60, 310-318. |
| 82 | Quilis, J. et al. (2008) The *Arabidopsis* AtNPR1 inversely modulates defense responses against fungal, bacterial, or viral pathogens while conferring hypersensitivity to abiotic stresses in transgenic rice. *Molecular Plant-Microbe Interactions*, 21, 1215-1231. |
| 83 | Ramos-Alvelo, M. et al. (2024) The SlDLK2 receptor, involved in the control of arbuscular mycorrhizal symbiosis, regulates hormonal balance in roots. *Frontiers in Microbiology*, 15, 1472449. |
| 84 | Ravet, K. et al. (2012) Iron and ROS control of the DownSTream mRNA decay pathway is essential for plant fitness. *The EMBO Journal*, 31, 175-186. |
| 85 | Rutter, M.T. et al. (2019) Distributed phenomics with the unPAK project reveals the effects of mutations. *The Plant Journal*, 100, 199-211. |
| 86 | Sattler, S.E. et al. (2013) Registration of N614, A3N615, N616, and N617 Shattercane Genetic Stocks with Cytoplasmic or Nuclear Male Sterility and Juicy or Dry Midribs. *Journal of Plant Registrations*, 7, 245-249. |
| 87 | Schaeffer, A. et al. (2001) The ratio of campesterol to sitosterol that modulates growth in *Arabidopsis* is controlled by STEROL METHYLTRANSFERASE 2; 1. *The Plant Journal*, 25, 605-615. |
| 88 | Schmalenbach, I. et al. (2014) Functional analysis of the Landsberg erecta allele of FRIGIDA. *BMC Plant Biology*, 14, 218. |
| 89 | Schulze, J. et al. (2013) Reduced clonal reproduction indicates low potential for establishment of hybrids between wild and cultivated strawberries (*Fragaria vesca* x *F. x ananassa*). *Ecological Research*, 28, 43-52. |
| 90 | Takita, E. et al. (2021) Development of the binary vector pTACAtg1 for stable gene expression in plant: Reduction of gene silencing in transgenic plants carrying the target gene with long flanking sequences. *Plant Biotechnology*, 38, 391-400. |
| 91 | Thakur, A.K. et al. (2012) Genetic modification of lignin biosynthetic pathway in *Populus ciliata* Wall. via *Agrobacterium*-mediated antisense CAD Gene transfer for quality paper production. *National Academy Science Letters*, 35, 79-84. |
| 92 | Tian, J. et al. (2012) The Mechanism of Antifungal Action of Essential Oil from Dill (*Anethum graveolens* L.) on *Aspergillus flavus*. *PLoS ONE*, 7, e30147. |
| 93 | Umnajkitikorn, K. et al. (2020) Silencing of OsCV (chloroplast vesiculation) maintained photorespiration and N assimilation in rice plants grown under elevated CO2. *Plant, Cell & Environment*, 43, 920-933. |
| 94 | Xu, M. et al. (2016) Developmental functions of miR156-regulated SQUAMOSA PROMOTER BINDING PROTEIN-LIKE (SPL) genes in *Arabidopsis thaliana*. *PLoS Genetics*, 12, e1006263. |
| 95 | Zafirov, D. et al. (2023) *Arabidopsis* eIF4E1 protects the translational machinery during TuMV infection and restricts virus accumulation. *PLoS Pathogens*, 19, e1011417. |
| 96 | Zhang, R. et al. (2017) Transgenic *Arabidopsis thaliana* containing increased levels of ATP and sucrose is more susceptible to *Pseudomonas syringae*. *PLoS ONE*, 12, e0171040. |
| 97 | Zhang, X. et al. (2020) Overexpression of a maize BR transcription factor ZmBZR1 in *Arabidopsis* enlarges organ and seed size of the transgenic plants. *Plant Science*, 292, 110378. |
| 98 | Sim, S.A. et al. (2019) FLOWERING HTH1 is involved in CONSTANS-mediated flowering regulation in *Arabidopsis*. *Applied Biological Chemistry*, 62. |
| 99 | Song, Z. et al. (2024) Comparative transcriptome analysis reveals nicotine metabolism is a critical component for enhancing stress response intensity of innate immunity system in tobacco. *Frontiers in Plant Science*, 15. |
| 100 | Stitz, M. et al. (2011) Diverting the flux of the JA pathway in *Nicotiana attenuata* compromises the plant's defense metabolism and fitness in nature and glasshouse. *PLoS ONE*, 6, e25925. |
| 101 | Stolze, A. et al. (2017) Development of rubber-enriched dandelion varieties by metabolic engineering of the inulin pathway. *Plant Biotechnology Journal*, 15, 740-753. |
| 102 | Straub, C.T. et al. (2020) Use of the lignocellulose-degrading bacterium *Caldicellulosiruptor bescii* to assess recalcitrance and conversion of wild-type and transgenic poplar. *Biotechnology for Biofuels*, 13, 1-10. |
| 103 | Tomassetti, S. et al. (2015) Controlled expression of pectic enzymes in *Arabidopsis thaliana* enhances biomass conversion without adverse effects on growth. *Phytochemistry*, 112, 221-230. |
| 104 | Tu, Y. et al. (2010) Functional analyses of caffeic acid O-Methyltransferase and Cinnamoyl-CoA-reductase genes from perennial ryegrass (*Lolium perenne*). *The Plant Cell*, 22, 3357-3373. |
| 105 | Tyagi, S. et al. (2019) SUPPRESSOR OF OVEREXPRESSION OF CONSTANS1 influences flowering time, lateral branching, oil quality, and seed yield in *Brassica juncea* cv. Varuna. *Functional & Integrative Genomics*, 19, 43-60. |
| 106 | von Bismarck, T. et al. (2023) Light acclimation interacts with thylakoid ion transport to govern the dynamics of photosynthesis in *Arabidopsis*. *New Phytologist*, 237, 160-176. |
| 107 | Wang, H. et al. (2022) STRIPE3, encoding a human dNTPase SAMHD1 homolog, regulates chloroplast development in rice. *Plant Science*, 323. |
| 108 | Wang, M. et al. (2018) *Nicotiana attenuata*'s capacity to interact with arbuscular mycorrhiza alters its competitive ability and elicits major changes in the leaf transcriptome. *Journal of Integrative Plant Biology*, 60, 242-261. |
| 109 | Wang, D. et al. (2023) Suppression of ETI by PTI priming to balance plant growth and defense through an MPK3/MPK6-WRKYs-PP2Cs module. *Molecular Plant*, 16, 903-918. |
| 110 | Weinig, C. et al. (2004) Testing adaptive plasticity to UV: costs and benefits of stem elongation and light-induced phenolics. *Evolution*, 58, 2645-2656. |
| 111 | Weiste, C. et al. (2017) The *Arabidopsis* bZIP11 transcription factor links low-energy signalling to auxin-mediated control of primary root growth. *PLoS Genetics*, 13. |
| 112 | Worakan, P. et al. (2022) Stable and reproducible expression of bacterial ipt gene under the control of SAM-specific promoter (pKNOX1) with interference of developmental patterns in transgenic *Peperomia pellucida* plants. *Frontiers in Plant Science*, 13, 984716. |
| 113 | Yoon, H.-K. et al. (2008) Regulation of Leaf Senescence by NTL9-mediated Osmotic Stress Signaling in *Arabidopsis*. *Molecules and Cells*, 25, 438-445. |
| 114 | Yu, X. et al. (2020) Wheat PP2C-a10 regulates seed germination and drought tolerance in transgenic *Arabidopsis*. *Plant Cell Reports*, 39, 635-651. |
| 115 | Yuan, X. et al. (2021) The APETALA2 homolog CaFFN regulates flowering time in pepper. *Horticulture Research*, 8. |
| 116 | Zhang RenShan et al. (2017) Transgenic *Arabidopsis thaliana* containing increased levels of ATP and sucrose is more susceptible to *Pseudomonas syringae*. *PLoS ONE*, 12, e0171040. |
| 117 | Zhou, F.-Y. et al. (2022) Overexpression of *Polypogon fugax* Type I-Like MADS-Box Gene PfAGL28 Affects Flowering Time and Pod Formation in Transgenic *Arabidopsis*. *Plant Molecular Biology Reporter*, 40, 188-196. |
| 118 | Zhou, F.-Y. et al. (2020) StMADS11 Subfamily Gene PfMADS16 From *Polypogon fugax* Regulates Early Flowering and Seed Development. *Frontiers in Plant Science*, 11. |
| 119 | Züst, T. et al. (2011) Using knockout mutants to reveal the growth costs of defensive traits. *Proceedings of the Royal Society B: Biological Sciences*, 278, 2598-2603. |
| **Reason: without data of fitness measurement of interest from LR/HR, or no setting of LR/HR plant.** | |
| 1 | J. H. Jørgensen and H. P. Jensen (1990) Effect of 'unnecessary' powdery mildew resistance genes on agronomic properties of spring barley. *Norsk Landbruksforsking*, 125-130. |
| 2 | Adugna, A. and Bekele, E. (2013) Morphology and fitness components of wild×crop F1 hybrids of *Sorghum bicolor* (L.) in Ethiopia: Implications for survival and introgression of crop genes in the wild pool. *Plant Genetic Resources*, 11, 196-205. |
| 3 | Adugna, A. and Bekele, E. (2017) Indirect estimates reveal the potential of transgene flow in the crop-wild-weed *Sorghum bicolor* complex in its centre of origin, Ethiopia. *Plant Genetic Resources*, 15, 496-505. |
| 4 | Albacete, A. et al. (2015) Ectopic overexpression of the cell wall invertase gene CIN1 leads to dehydration avoidance in tomato. *Journal of Experimental Botany*, 66, 863-878. |
| 5 | Aono, M. et al. (2006) Detection of feral transgenic oilseed rape with multiple-herbicide resistance in Japan. *Environmental Biosafety Research*, 5, 77-87. |
| 6 | Bai, Y. et al. (2008) Naturally occurring broad-spectrum powdery mildew resistance in a Central American tomato accession is caused by loss of Mlo function. *Molecular Plant-Microbe Interactions*, 21, 30-39. |
| 7 | Bartsch, D. (1999) Risk Analysis on the Ecological Impact of Gene Flow to Wild Relatives: The Transgenic Sugar Beet Example. *Nature Biotechnology*, 17, 22-22. |
| 8 | Baucom, R.S. and Mauricio, R. (2004) Fitness costs and benefits of novel herbicide tolerance in a noxious weed. *Proceedings of the National Academy of Sciences*, 101, 13386-13390. |
| 9 | Brar, G.S. et al. (1994) Recovery of transgenic peanut (*Arachis hypogaea* L.) plants from elite cultivars utilizing ACCELL technology. *The Plant Journal*, 5, 745-753. |
| 10 | Busi, R. and Powles, S.B. (2016) Transgenic glyphosate-resistant canola (*Brassica napus*) can persist outside agricultural fields in Australia. *Agriculture, Ecosystems & Environment*, 220, 28-34. |
| 11 | Chai, T.-T. et al. (2010) Alternative oxidase, a determinant of plant gametophyte fitness and fecundity. *Plant Signaling & Behavior*, 5, 604-606. |
| 12 | Chakrabarti, S.K. et al. (2006) Expression of the cry9Aa2 Bt gene in tobacco chloroplasts confers resistance to potato tuber moth. *Transgenic Research*, 15, 481-488. |
| 13 | Chekan, J.R. et al. (2019) Molecular basis for enantioselective herbicide degradation imparted by aryloxyalkanoate dioxygenases in transgenic plants. *Proceedings of the National Academy of Sciences*, 116, 13299-13304. |
| 14 | Chen, S.-F. et al. (2016) Ectopic expression of a tobacco vacuolar invertase inhibitor in guard cells confers drought tolerance in *Arabidopsis*. *Journal of Enzyme Inhibition and Medicinal Chemistry*, 31, 1381-1385. |
| 15 | Chow, H.T. and Ng, D.W.-K. (2017) Regulation of miR163 and its targets in defense against *Pseudomonas syringae* in *Arabidopsis thaliana*. *Scientific Reports*, 7, 46433. |
| 16 | Cooper, J.I. (2004) The hazard of ecological release in wild relatives of transgenic TuMV-tolerant brassicas. |
| 17 | Corrado, G. et al. (2011) Systemin-inducible defence against pests is costly in tomato. *Biologia Plantarum*, 55, 305-311. |
| 18 | Corrado, G. et al. (2007) Systemin regulates both systemic and volatile signaling in tomato plants. *Journal of Chemical Ecology*, 33, 669-681. |
| 19 | Cosa, B.D. et al. (2001) Overexpression of the Bt cry2Aa2 operon in chloroplasts leads to formation of insecticidal crystals. *Nature Biotechnology*, 19, 71-74. |
| 20 | Dahal, K. et al. (2012) The effects of phenotypic plasticity on photosynthetic performance in winter rye, winter wheat and *Brassica napus*. *Physiologia Plantarum*, 144, 169-188. |
| 21 | Darmency, H. (1994) The impact of hybrids between genetically modified crop plants and their related species: introgression and weediness. *Molecular Ecology*, 3, 37-40. |
| 22 | Doebley, J. (1990) Molecular evidence for gene flow among *Zea* species. *Bioscience*, 40, 443-448. |
| 23 | Eckes, P. et al. (1989) Overproduction of alfalfa glutamine synthetase in transgenic tobacco plants. *Molecular and General Genetics MGG*, 217, 263-268. |
| 24 | Endo, T. et al. (2020) Fast-track breeding system to introduce CTV resistance of trifoliate orange into citrus germplasm, by integrating early flowering transgenic plants with marker-assisted selection. *BMC Plant Biology*, 20, 1-16. |
| 25 | Farnham, G. and Baulcombe, D.C. (2006) Artificial evolution extends the spectrum of viruses that are targeted by a disease-resistance gene from potato. *Proceedings of the National Academy of Sciences*, 103, 18828-18833. |
| 26 | Fartyal, D. et al. (2018) Developing dual herbicide tolerant transgenic rice plants for sustainable weed management. *Scientific Reports*, 8, 11598. |
| 27 | Feechan, A. et al. (2015) Strategies for RUN1 Deployment Using RUN2 and REN2 to Manage Grapevine Powdery Mildew Informed by Studies of Race Specificity. *Phytopathology*, 105, 1104-1113. |
| 28 | Fiorani, F. et al. (2005) The alternative oxidase of plant mitochondria is involved in the acclimation of shoot growth at low temperature. A study of *Arabidopsis* AOX1a transgenic plants. *Plant Physiology*, 139, 1795-1805. |
| 29 | Franceschetti, M. et al. (2004) Expression proteomics identifies biochemical adaptations and defense responses in transgenic plants with perturbed polyamine metabolism. *FEBS Letters*, 576, 477-480. |
| 30 | Fritz, M.L. et al. (2020) Mutations in a Novel Cadherin Gene Associated with Bt Resistance in *Helicoverpa zea*. *G3: Genes, Genomes, Genetics*, 10, 1563-1574. |
| 31 | Gao, J.-J. et al. (2011) Forced expression of Mdmyb10, a myb transcription factor gene from apple, enhances tolerance to osmotic stress in transgenic *Arabidopsis*. *Molecular Biology Reports*, 38, 205-211. |
| 32 | Gassmann, A.J. and Futuyma, D.J. (2005) Consequence of herbivory for the fitness cost of herbicide resistance: photosynthetic variation in the context of plant–herbivore interactions. *Journal of Evolutionary Biology*, 18, 447-454. |
| 33 | Giacomelli, L. et al. (2023) Simultaneous editing of two DMR6 genes in grapevine results in reduced susceptibility to downy mildew. *Frontiers in Plant Science*, 14, 1242240. |
| 34 | Grand, X. et al. (2012) Identification of positive and negative regulators of disease resistance to rice blast fungus using constitutive gene expression patterns. *Plant Biotechnology Journal*, 10, 840-850. |
| 35 | Gu, X. et al. (2017) RdreB1BI enhances drought tolerance by activating AQP-related genes in transgenic strawberry. *Plant Physiology and Biochemistry*, 119, 33-42. |
| 36 | Gutierrez-Campos, R. et al. (1999) The use of cysteine proteinase inhibitors to engineer resistance against potyviruses in transgenic tobacco plants. *Nature Biotechnology*, 17, 1223-1226. |
| 37 | Hails, R.S. (2006) The potential invasiveness of insect-resistant transgenic plants. |
| 38 | Himanen, S.J. et al. (2008) Interactions of elevated carbon dioxide and temperature with aphid feeding on transgenic oilseed rape: Are *Bacillus thuringiensis* (Bt) plants more susceptible to nontarget herbivores in future climate? *Global Change Biology*, 14, 1437-1454. |
| 39 | Horsman, J. et al. (2007) Picloram resistance in transgenic tobacco expressing an anti-picloram scFv antibody is due to reduced translocation. *Journal of Agricultural and Food Chemistry*, 55, 106-112. |
| 40 | Hu, L. et al. (2016) OsWRKY53, a versatile switch in regulating herbivore-induced defense responses in rice. *Plant Signaling & Behavior*, 11, e1169357. |
| 41 | Huang, F. (2021) Resistance of the fall armyworm, *Spodoptera frugiperda*, to transgenic *Bacillus thuringiensis* Cry1F corn in the Americas: lessons and implications for Bt corn IRM in China. *Insect Science*, 28, 574-589. |
| 42 | Jiang, Y. et al. (2022) Optimized prime editing efficiently generates glyphosate-resistant rice plants carrying homozygous TAP-IVS mutation in EPSPS. *Molecular Plant*, 15, 1646-1649. |
| 43 | Jin, L. et al. (2015) Large-scale test of the natural refuge strategy for delaying insect resistance to transgenic Bt crops. *Nature Biotechnology*, 33, 169-174. |
| 44 | Jin, S.-B. et al. (1999) Construction of Citrus Transgenic Plant with Fatty Acid Desaturase Gene. *Journal of Applied Biological Chemistry*, 42, 113-118. |
| 45 | Jung, H.W. et al. (2020) Pathogen-Associated Molecular Pattern-Triggered Immunity Involves Proteolytic Degradation of Core Nonsense-Mediated mRNA Decay Factors During the Early Defense Response. *The Plant Cell*, 32, 1081-1101. |
| 46 | Kappers, I.F. et al. (2005) Genetic engineering of terpenoid metabolism attracts bodyguards to *Arabidopsis*. *Science*, 309, 2070-2072. |
| 47 | Kelly, C.K. et al. (2005) An analytical model assessing the potential threat to natural habitats from insect resistance transgenes. *Proceedings of the Royal Society B: Biological Sciences*, 272, 1759-1767. |
| 48 | Kim, D.Y. et al. (2021) Natural hybridization between transgenic and wild soybean genotypes. *Plant Biotechnology Reports*, 15, 299-308. |
| 49 | Klińska-Bąchor, S. et al. (2023) Phospholipid:diacylglycerol acyltransferase1-overexpression stimulates lipid turnover, oil production and fitness in cold-grown plants. *BMC Plant Biology*, 23, 370. |
| 50 | Knight, K.M. et al. (2021) Successful development and implementation of a practical proactive resistance management plan for Bt cotton in Australia. *Pest Management Science*, 77, 4262-4273. |
| 51 | Knox, O.G.G. et al. (2007) Constitutive expression of Cry proteins in roots and border cells of transgenic cotton. *Euphytica*, 154, 83-90. |
| 52 | Koseoglou, E. et al. (2023) Inactivation of tomato WAT1 leads to reduced susceptibility to *Clavibacter michiganensis* through downregulation of bacterial virulence factors. *Frontiers in Plant Science*, 14, 1082094. |
| 53 | Kruger, M. et al. (2014) No fitness costs associated with resistance of *Busseola fusca* (Lepidoptera: Noctuidae) to genetically modified Bt maize. *Crop Protection*, 55, 1-6. |
| 54 | La Paz, J.L. et al. (2014) The Use of Massive Sequencing to Detect Differences between Immature Embryos of MON810 and a Comparable Non-GM Maize Variety. *PLoS ONE*, 9, e100895. |
| 55 | Lai, Y. et al. (2020) The *Arabidopsis* PHD-finger protein EDM2 has multiple roles in balancing NLR immune receptor gene expression. *PLoS Genetics*, 16, e1008993. |
| 56 | Laine, A.-L. (2016) Disease resistance: not so costly after all. *Nature Plants*, 2, 1-2. |
| 57 | Leach, J.E. et al. (2001) Pathogen fitness penalty as a predictor of durability of disease resistance genes. *Annual Review of Phytopathology*, 39, 187-224. |
| 58 | Lee, H.-H. et al. (2018) Root-specific expression of defensin in transgenic tobacco results in enhanced resistance against *Phytophthora parasitica* var. *nicotianae*. *European Journal of Plant Pathology*, 151, 811-823. |
| 59 | Lee, H.G. et al. (2016) Increased STM expression is associated with drought tolerance in *Arabidopsis*. *Journal of Plant Physiology*, 201, 79-84. |
| 60 | Légère, A. (2005) Risks and consequences of gene flow from herbicide-resistant crops: canola (*Brassica napus* L) as a case study. *Pest Management Science*, 61, 292-300. |
| 61 | Lei, L. et al. (2011) Dynamic expression of green fluorescent protein and *Bacillus thuringiensis* Cry1Ac endotaxin in interspecific hybrids and successive backcross generations (BC1 and BC2) between transgenic *Brassica napus* crop and wild *Brassica juncea*. *Annals of Applied Biology*, 159, 212-219. |
| 62 | Li, Y. et al. (2010) Regulation of the Expression of Plant Resistance Gene SNC1 by a Protein with a Conserved BAT2 Domain. *Plant Physiology*, 153, 1425-1434. |
| 63 | Liu, Y. et al. (2013) Consequences of gene flow between oilseed rape (*Brassica napus*) and its relatives. *Plant Science*, 211, 42-51. |
| 64 | Liu, Y. et al. (2024) Overriding Mendelian inheritance in *Arabidopsis* with a CRISPR toxin–antidote gene drive that impairs pollen germination. *Nature Plants*, 10, 910-922. |
| 65 | Liu, X.Q. et al. (2005) OsWRKY03, a rice transcriptional activator that functions in defense signaling pathway upstream of OsNPR1. *Cell Research*, 15, 593-603. |
| 66 | Liu, Z. et al. (2011) Overexpression of a resveratrol synthase gene (PcRS) from *Polygonum cuspidatum* in transgenic *Arabidopsis* causes the accumulation of trans-piceid with antifungal activity. *Plant Cell Reports*, 30, 2027-2036. |
| 67 | Lohn, A.F. et al. (2021) Transgene behavior in genetically modified teosinte hybrid plants: transcriptome expression, insecticidal protein production and bioactivity against a target insect pest. *Environmental Sciences Europe*, 33, 67. |
| 68 | Lu, C. et al. (2002) Destiny of a transgene escape from *Brassica napus* into *Brassica rapa*. *Theoretical and Applied Genetics*, 105, 78-84. |
| 69 | Luan, H. et al. (2020) Transgenic plant generated by RNAi-mediated knocking down of soybean Vma12 and soybean mosaic virus resistance evaluation. *AMB Express*, 10, 1-10. |
| 70 | Malabadi, R.B. and Nataraja, K. (2007) Production of transgenic plants via *Agrobacterium tumefaciens* mediated genetic transformation in *Pinus wallichiana* (Himalayan blue pine). *Transgenic Plant Journal*, 1, 376-383. |
| 71 | Manikandan, R. et al. (2016) Transformation of tobacco (*Nicotiana tabaccum*) with cry2AX1 gene and analysis of transgenic plants. |
| 72 | Mark Tepfer (2000) Potential risks associated with virus-resistant transgenic plants. *The Biosafety of Genetically Modified Organisms*, 89. |
| 73 | Marvier, M.A. et al. (1999) How do the design of monitoring and control strategies affect the chance of detecting and containing transgenic weeds? *Methods for Risk Assessment of Transgenic Plants: III*, 109-122. |
| 74 | Massange-Sánchez, J.A. et al. (2016) Overexpression of Grain Amaranth (*Amaranthus hypochondriacus*) AhERF or AhDOF Transcription Factors in *Arabidopsis thaliana* Increases Water Deficit- and Salt-Stress Tolerance, Respectively, via Contrasting Stress-Amelioration Mechanisms. *PLoS ONE*, 11, e0164280. |
| 75 | Meng, F.-Z. et al. (2023) The velvet family proteins mediate low resistance to isoprothiolane in *Magnaporthe oryzae*. *PLoS Pathogens*, 19, e1011011. |
| 76 | Merotto, A. et al. (2016) Evolutionary and social consequences of introgression of nontransgenic herbicide resistance from rice to weedy rice in Brazil. *Evolutionary Applications*, 9, 837-846. |
| 77 | Messeguer, J. et al. (2001) Field assessments of gene flow from transgenic to cultivated rice (*Oryza sativa* L.) using a herbicide resistance gene as tracer marker. *Theoretical and Applied Genetics*, 103, 1151-1159. |
| 78 | Miaw, C.S.W. et al. (2017) Single-laboratory validation of a method for detection of Roundup Ready soy in soybeans: application of new strategies for qualitative validation. *Quality Assurance and Safety of Crops & Foods*, 9, 105-114. |
| 79 | Mimura, M. et al. (2008) Impact of environmental stress-tolerant transgenic potato on genotypic diversity of microbial communities and soil enzyme activities under stress conditions. *Microbes and Environments*, 23, 221-228. |
| 80 | Mitra, S. et al. (2021) Negative regulation of plastidial isoprenoid pathway by herbivore-induced beta-cyclocitral in *Arabidopsis thaliana*. *Proceedings of the National Academy of Sciences*, 118, e2008747118. |
| 81 | Moon, H.S. (2011) Developing biocontainment strategies to suppress transgene escape via pollen dispersal from transgenic plants. |
| 82 | Morris, W.F. et al. (1994) Do barren zones and pollen traps reduce gene escape from transgenic crops? *Ecological Applications*, 4, 157-165. |
| 83 | Morroni, M. et al. (2013) Deep Sequencing of Recombinant Virus Populations in Transgenic and Nontransgenic Plants Infected with *Cucumber mosaic virus*. *Molecular Plant-Microbe Interactions*, 26, 801-811. |
| 84 | Mwimba, M. et al. (2018) Daily humidity oscillation regulates the circadian clock to influence plant physiology. *Nature Communications*, 9, 4290. |
| 85 | Nadal, A. et al. (2012) Constitutive expression of transgenes encoding derivatives of the synthetic antimicrobial peptide BP100: impact on rice host plant fitness. *BMC Plant Biology*, 12, 159. |
| 86 | Nagadhara, D. et al. (2004) Transgenic rice plants expressing the snowdrop lectin gene (gna) exhibit high-level resistance to the whitebacked planthopper (*Sogatella furcifera*). *Theoretical and Applied Genetics*, 109, 1399-1405. |
| 87 | Nam, K.-H. and Han, S.M. (2020) Seed germination of sunflower as a case study for the risk assessment and management of transgenic plants used for environmental remediation in South Korea. *Sustainability*, 12, 10110. |
| 88 | Neubauer, M. and Innes, R.W. (2020) Loss of the Acetyltransferase NAA50 Induces Endoplasmic Reticulum Stress and Immune Responses and Suppresses Growth. *Plant Physiology*, 183, 1838-1854. |
| 89 | Ni, M. et al. (2017) Next-generation transgenic cotton: pyramiding RNAi and Bt counters insect resistance. *Plant Biotechnology Journal*, 15, 1204-1213. |
| 90 | Nir, I.D.O. et al. (2014) The *Arabidopsis* GIBBERELLIN METHYL TRANSFERASE 1 suppresses gibberellin activity, reduces whole-plant transpiration and promotes drought tolerance in transgenic tomato. *Plant, Cell & Environment*, 37, 113-123. |
| 91 | Oger, P. et al. (1997) Genetically engineered plants producing opines alter their biological environment. *Nature Biotechnology*, 15, 369-372. |
| 92 | Orgil, U. et al. (2007) Intraspecific genetic variations, fitness cost and benefit of RPW8, a disease resistance locus in *Arabidopsis thaliana*. *Genetics*, 176, 2317-2333. |
| 93 | Oswald, K.J. et al. (2012) Assessment of fitness costs in Cry3Bb1-resistant and susceptible western corn rootworm (Coleoptera: Chrysomelidae) laboratory colonies. *Journal of Applied Entomology*, 136, 730-740. |
| 94 | Pan, X. (2019) Determining Pollen-Mediated Gene Flow in Transgenic Cotton. *Transgenic Cotton: Methods and Protocols*, 309-321. |
| 95 | Pan, X. et al. (2019) Quantification of root anatomical traits in RGP transgenic maize plants based on Micro-CT. *Computer and Computing Technologies in Agriculture XI*, 340-346. |
| 96 | Park, J.-E. et al. (2007) GH3-mediated auxin homeostasis links growth regulation with stress adaptation response in *Arabidopsis*. *Journal of Biological Chemistry*, 282, 10036-10046. |
| 97 | Parsons, P.A. (1992) Fluctuating asymmetry: a biological monitor of environmental and genomic stress. *Heredity*, 68, 361-364. |
| 98 | Paschold, A. et al. (2006) Using 'mute' plants to translate volatile signals. *The Plant Journal*, 45, 275-291. |
| 99 | Pavei, D. et al. (2016) Response to water stress in transgenic ('p5cs' gene) wheat plants ('*Triticum aestivum*' L.). *Australian Journal of Crop Science*, 10, 776-783. |
| 100 | Pérez-Díaz, J. et al. (2021) *Prunus* Hexokinase 3 genes alter primary C-metabolism and promote drought and salt stress tolerance in *Arabidopsis* transgenic plants. *Scientific Reports*, 11, 7098. |
| 101 | Phung, T.-H. et al. (2011) Porphyrin biosynthesis control under water stress: sustained porphyrin status correlates with drought tolerance in transgenic rice. *Plant Physiology*, 157, 1746-1764. |
| 102 | Pickett, J. (2012) Indirect routes to reproductive success. *eLife*, 1, e00240. |
| 103 | Porri, A. et al. (2023) Inhibition profile of trifludimoxazin towards PPO2 target site mutations. *Pest Management Science*, 79, 507-519. |
| 104 | Prändl, R. et al. (1998) HSF3, a new heat shock factor from *Arabidopsis thaliana*, derepresses the heat shock response and confers thermotolerance when overexpressed in transgenic plants. *Molecular and General Genetics MGG*, 258, 269-278. |
| 105 | Prendeville, H.R. et al. (2012) Virus infections in wild plant populations are both frequent and often unapparent. *American Journal of Botany*, 99, 1033-1042. |
| 106 | Quilis, J. et al. (2014) Inducible expression of a fusion gene encoding two proteinase inhibitors leads to insect and pathogen resistance in transgenic rice. *Plant Biotechnology Journal*, 12, 367-377. |
| 107 | Rabelo, M.M. et al. (2020) Demographic Performance of *Helicoverpa zea* Populations on Dual and Triple-Gene Bt Cotton. *Toxins*, 12, 551. |
| 108 | Ramos, P.L. et al. (2004) Identification of the minimal sequence required for vascular-specific activity of *Tomato mottle Taino virus* replication-associated protein promoter in transgenic plants. *Virus Research*, 102, 125-132. |
| 109 | Ramstein, G.P. and Buckler, E.S. (2022) Prediction of evolutionary constraint by genomic annotations improves functional prioritization of genomic variants in maize. *Genome Biology*, 23, 183. |
| 110 | Raybould, A.F. et al. (1999) The prevalence and spatial distribution of viruses in natural populations of *Brassica oleracea*. *New Phytologist*, 141, 265-275. |
| 111 | Raybould, A. and Cooper, I. (2005) Tiered tests to assess the environmental risk of fitness changes in hybrids between transgenic crops and wild relatives: the example of virus resistant *Brassica napus*. *Environmental Biosafety Research*, 4, 127-140. |
| 112 | Reichman, J.R. et al. (2006) Establishment of transgenic herbicide-resistant creeping bentgrass (*Agrostis stolonifera* L.) in nonagronomic habitats. *Molecular Ecology*, 15, 4243-4255. |
| 113 | Reyes-Rosales, A. et al. (2023) Identification of genetic and biochemical mechanisms associated with heat shock and heat stress adaptation in grain amaranths. *Frontiers in Plant Science*, 14, 1101375. |
| 114 | Richter, K.S. et al. (2014) Somatic homologous recombination in plants is promoted by a geminivirus in a tissue-selective manner. *Virology*, 452-453, 287-296. |
| 115 | Rohini, V.K. and Rao, K.S. (2000) Transformation of peanut (*Arachis hypogaea* L.): a non-tissue culture based approach for generating transgenic plants. *Plant Science*, 150, 41-49. |
| 116 | Roux, F. and Reboud, X. (2007) Herbicide resistance dynamics in a spatially heterogeneous environment. *Crop Protection*, 26, 335-341. |
| 117 | Runyon, J.B. and Birdsall, J.L. (2016) Costs of induced defenses for the invasive plant houndstongue (*Cynoglossum officinale* L.) and the potential importance for weed biocontrol. *Arthropod-Plant Interactions*, 10, 383-391. |
| 118 | Ryder, L.S. et al. (2012) Saprotrophic competitiveness and biocontrol fitness of a genetically modified strain of the plant-growth-promoting fungus *Trichoderma hamatum* GD12. *Microbiology-SGM*, 158, 84-97. |
| 119 | Ryu, C. et al. (2007) A two-strain mixture of rhizobacteria elicits induction of systemic resistance against *Pseudomonas syringae* and *Cucumber mosaic virus* coupled to promotion of plant growth on *Arabidopsis thaliana*. *Journal of Microbiology and Biotechnology*, 17, 280. |
| 120 | Saji, H. et al. (2005) Monitoring the escape of transgenic oilseed rape around Japanese ports and roadsides. *Environmental Biosafety Research*, 4, 217-222. |
| 121 | Samantsidis, G.-R. et al. (2020) 'What I cannot create, I do not understand': functionally validated synergism of metabolic and target site insecticide resistance. *Proceedings of the Royal Society B: Biological Sciences*, 287, 20200838. |
| 122 | Sane, V.A. et al. (1997) Development of insect-resistant transgenic plants using plant genes: Expression of cowpea trypsin inhibitor in transgenic tobacco plants. *Current Science*, 72, 741-747. |
| 123 | Santos-Amaya, O.F. et al. (2015) Resistance to dual-gene Bt maize in *Spodoptera frugiperda*: selection, inheritance and cross-resistance to other transgenic events. *Scientific Reports*, 5, 18243. |
| 124 | Schöffl, F. et al. (1987) Constitutive transcription of a soybean heat-shock gene by a cauliflower mosaic virus promoter in transgenic tobacco plants. *Developmental Genetics*, 8, 365-374. |
| 125 | Schouten, H.J. et al. (2017) Re-sequencing transgenic plants revealed rearrangements at T-DNA inserts, and integration of a short T-DNA fragment, but no increase of small mutations elsewhere. *Plant Cell Reports*, 36, 493-504. |
| 126 | Schubert, J. et al. (2004) Transgenic potatoes with resistance to *Potato virus Y*-results of several years' field trials for studying stability and biosafety aspects. |
| 127 | Tan, L. et al. (2014) The *Xanthomonas* campestris effector protein XopDXcc8004 triggers plant disease tolerance by targeting DELLA proteins. *New Phytologist*, 204, 595-608. |
| 128 | Thakur, N. et al. (2014) Enhanced whitefly resistance in transgenic tobacco plants expressing double stranded RNA of v-ATPase A gene. *PLoS ONE*, 9, e87235. |
| 129 | Tian, J.-C. et al. (2012) Transgenic Cry1Ab rice does not impact ecological fitness and predation of a generalist spider. *PLoS ONE*, 7, e35164. |
| 130 | Vazquez-Barrios, V. et al. (2021) Ongoing ecological and evolutionary consequences by the presence of transgenes in a wild cotton population. *Scientific Reports*, 11, 1959. |
| 131 | Wang, X. et al. (2024) Inheritance and ecological effects of exogenous genes from transgenic *Brassica napus* to *Brassica juncea* hybrids. *Plant Science*, 349, 112245. |
| 132 | Xu, R. et al. (2021) Identification of herbicide resistance OsACC1 mutations via in planta prime-editing-library screening in rice. *Nature Plants*, 7, 888-892. |
| 133 | Xu, H.-X. et al. (2017) Effects of Transgenic Rice Infected with SRBSDV on Bt expression and the Ecological Fitness of Non-vector Brown Planthopper *Nilaparvata lugens*. *Scientific Reports*, 7, 6328. |
| 134 | Xu, L. et al. (2017) Overexpression of *Arabidopsis* DREB1A gene in transgenic *Poa pratensis*: Impacts on osmotic adjustment and hormone metabolism under drought. *International Turfgrass Society Research Journal*, 13, 527-536. |
| 135 | Yadav, S. et al. (2024) Cry2Aa Delta-Endotoxin Confers Strong Resistance Against Brinjal Fruit and Shoot Borer in Transgenic Brinjal (*Solanum melongena* L.) Plants. *Pakistan Journal of Zoology*, 56. |
| 136 | Yan, S. et al. (2020) Intercrops can mitigate pollen-mediated gene flow from transgenic cotton while simultaneously reducing pest densities. *Science of the Total Environment*, 711, 134855. |
| 137 | Yang, S. et al. (2013) Rapidly evolving R genes in diverse grass species confer resistance to rice blast disease. *Proceedings of the National Academy of Sciences*, 110, 18572-18577. |
| 138 | Yang, F. et al. (2022) Effects of cross-pollination among non-Bt and pyramided Bt corn expressing cry proteins in seed mixtures on resistance development of dual-gene resistant *Helicoverpa zea*. *Pest Management Science*, 78, 3260-3265. |
| 139 | Zanatta, C.B. et al. (2020) Stacked genetically modified soybean harboring herbicide resistance and insecticide rCry1Ac shows strong defense and redox homeostasis disturbance after glyphosate-based herbicide application. *Environmental Sciences Europe*, 32, 1-17. |
| 140 | Zhang, D. et al. (2011) Transgenic plants of *Petunia hybrida* harboring the CYP2E1 gene efficiently remove benzene and toluene pollutants and improve resistance to formaldehyde. *Genetics and Molecular Biology*, 34, 634-639. |
| 141 | Zhang, Y. et al. (2013) Overexpression of a novel Cry1Ie gene confers resistance to Cry1Ac-resistant cotton bollworm in transgenic lines of maize. *Plant Cell, Tissue and Organ Culture (PCTOC)*, 115, 151-158. |
| 142 | Schwekendiek, A. et al. (2007) Constitutive expression of a grapevine stilbene synthase gene in transgenic hop (*Humulus lupulus* L.) yields resveratrol and its derivatives in substantial quantities. *Journal of Agricultural and Food Chemistry*, 55, 7002-7009. |
| **Reason: without controlling (or descriptions) of genetic background between LR and HR.** | |
| 1 | Alia et al. (1998) Enhancement of the tolerance of *Arabidopsis* to high temperatures by genetic engineering of the synthesis of glycinebetaine. *The Plant Journal*, 16, 155-161. |
| 2 | Hauser, T.P. et al. (1998) Fitness of backcross and F2 hybrids between weedy *Brassica rapa* and oilseed rape (*B. napus*). *Heredity*, 81, 436-443. |
| 3 | Adam, K.D. and Köhler, W.H. (1996) Evolutionary genetic considerations on the goals and risks in releasing transgenic crops. *Transgenic Organisms: Biological and Social Implications*, 59-79. |
| 4 | Ahuja, M.R. (2011) Fate of transgenes in the forest tree genome. *Tree Genetics & Genomes*, 7, 221-230. |
| 5 | Ahuja, M.R. (2009) Transgene stability and dispersal in forest trees. *Trees*, 23, 1125-1135. |
| 6 | Al-Ahmad, H. et al. (2006) Mitigation of establishment of *Brassica napus* transgenes in volunteers using a tandem construct containing a selectively unfit gene. *Plant Biotechnology Journal*, 4, 7-21. |
| 7 | Allainguillaume, J. et al. (2006) Fitness of hybrids between rapeseed (*Brassica napus*) and wild *Brassica rapa* in natural habitats. *Molecular Ecology*, 15, 1175-1184. |
| 8 | Ammitzbøll, H. et al. (2005) Transgene expression and fitness of hybrids between GM oilseed rape and *Brassica rapa*. *Environmental Biosafety Research*, 4, 3-12. |
| 9 | Bartsch, D. et al. (2003) Environmental implications of gene flow from sugar beet to wild beet--current status and future research needs. *Environmental Biosafety Research*, 2, 105-115. |
| 10 | Betts, S.D. et al. (2019) Uniform Expression and Relatively Small Position Effects Characterize Sister Transformants in Maize and Soybean. *Frontiers in Plant Science*, 10, 1209. |
| 11 | Brown, J. et al. (1996) Competitive and reproductive fitness of transgenic canola×weed species hybrids. |
| 12 | Busconi, M. et al. (2012) Spread of herbicide-resistant weedy rice (red rice, *Oryza sativa* L.) after 5 years of Clearfield rice cultivation in Italy. *Plant Biology*, 14, 751-759. |
| 13 | Cao, Q.-J. et al. (2009) Performance of hybrids between weedy rice and insect-resistant transgenic rice under field experiments: implication for environmental biosafety assessment. *Journal of Integrative Plant Biology*, 51, 1138-1148. |
| 14 | Chauhan, H. et al. (2015) The wheat resistance gene Lr34 results in the constitutive induction of multiple defense pathways in transgenic barley. *The Plant Journal*, 84, 202-215. |
| 15 | Chen, J. et al. (2021) Dinitroaniline herbicide resistance and mechanisms in weeds. *Frontiers in Plant Science*, 12, 634018. |
| 16 | Cipollini, D.F. (2002) Does competition magnify the fitness costs of induced responses in *Arabidopsis thaliana*? A manipulative approach. *Oecologia*, 131, 514-520. |
| 17 | Corbett, B.P. et al. (2011) The effects of root-knot nematode infection and mi-mediated nematode resistance in tomato on plant fitness. *Journal of Nematology*, 43, 82. |
| 18 | Fan, A. et al. (2022) Heterologous expression of the *Haynaldia villosa* pattern-recognition receptor CERK1-V in wheat increases resistance to three fungal diseases. *Crop Journal*, 10, 1733-1745. |
| 19 | Frey, T.J. et al. (2011) Fitness evaluation of Rcg1, a locus that confers resistance to *Colletotrichum graminicola* (Ces.) GW Wils. using near-isogenic maize hybrids. *Crop Science*, 51, 1551-1563. |
| 20 | Guéritaine, G. et al. (2003) Emergence and growth of hybrids between *Brassica napus* and *Raphanus raphanistrum*. *New Phytologist*, 158, 561-567. |
| 21 | Hauser, T.P. et al. (1998) Fitness of F1 hybrids between weedy *Brassica rapa* and oilseed rape (*B. napus*). *Heredity*, 81, 429-435. |
| 22 | Hayakawa, T. et al. (1992) Genetically engineered rice resistant to rice stripe virus, an insect-transmitted virus. *Proceedings of the National Academy of Sciences*, 89, 9865-9869. |
| 23 | Hernández-Walias, F.J. et al. (2022) Transgenerational Tolerance to Salt and Osmotic Stresses Induced by Plant Virus Infection. *International Journal of Molecular Sciences*, 23, 12497. |
| 24 | Hooftman, D.A.P. et al. (2015) Seed bank dynamics govern persistence of *Brassica* hybrids in crop and natural habitats. *Annals of Botany*, 115, 147-157. |
| 25 | Hu, J. et al. (2017) An Empirical Assessment of Transgene Flow from a Bt Transgenic Poplar Plantation. *PLoS ONE*, 12. |
| 26 | Hu, Y.-q. et al. (2022) Sexual compatibility of transgenic soybean and different wild soybean populations. *Journal of Integrative Agriculture*, 21, 36-48. |
| 27 | Huang, Y. et al. (2019) Fitness of F1 hybrids between stacked transgenic rice T1c-19 with *cry1C*/bar genes and weedy rice. *Journal of Integrative Agriculture*, 18, 2793-2805. |
| 28 | Huang, L. et al. (2024) Fitness of the first backcross generations from the second to the sixth progenies of glyphosate-resistant transgenic *Brassica napus* and wild *Brassica juncea* in absence of the herbicide. *Journal of Plant Ecology*, 17, rtad030. |
| 29 | Huangfu, C. et al. (2011) Performance of hybrids between transgenic oilseed rape (*Brassica napus*) and wild *Brassica juncea*: An evaluation of potential for transgene escape. *Crop Protection*, 30, 57-62. |
| 30 | Ji, X.-q. et al. (2023) Fitness analysis of F2 hybrids from glyphosate-resistant transgenic soybean and wild soybean of Baotou, Inner Mongolia. |
| 31 | Kan GuiZhen et al. (2015) Fitness of hybrids between wild soybeans (*Glycine soja*) and the glyphosate-resistant transgenic soybean (*Glycine max*). |
| 32 | Klinger, T. and Ellstrand, N.C. (1994) Engineered genes in wild populations: fitness of weed-crop hybrids of *Raphanus sativus*. *Ecological Applications*, 4, 117-120. |
| 33 | Landbo, L. and Jørgensen, R.B. (1997) Seed germination in weedy *Brassica campestris* and its hybrids with *B. napus*: implications for risk assessment of transgenic oilseed rape. *Euphytica*, 97, 209-216. |
| 34 | Li, L. et al. (2023) Fitness of F1, F2 and F3 from T2A-1 transgenic rice stacked with Cry2A*/bar genes and weedy rice. |
| 35 | Liang, R. et al. (2022) Fitness and Rhizobacteria of F2, F3 Hybrids of Herbicide-Tolerant Transgenic Soybean and Wild Soybean. *Plants*, 11, 3184. |
| 36 | Liang, R. et al. (2023) Fitness and Hard Seededness of F2 and F3 Descendants of Hybridization between Herbicide-Resistant *Glycine max* and *G. soja*. *Plants*, 12, 3671. |
| 37 | Liu, J.Y. et al. (2021) Fitness of F1 hybrids between 10 maternal wild soybean populations and transgenic soybean. *Transgenic Research*, 30, 105-119. |
| 38 | Liu YongBo et al. (2012) The effects of seed size on hybrids formed between oilseed rape (*Brassica napus*) and wild brown mustard (*B. juncea*). |
| 39 | Lu, G. et al. (2014) Rice LTG1 is involved in adaptive growth and fitness under low ambient temperature. *The Plant Journal*, 78, 468-480. |
| 40 | Mercer, K.L. et al. (2006) Effects of Competition on the Fitness of Wild and Crop-Wild Hybrid Sunflower from a Diversity of Wild Populations and Crop Lines. *Evolution*, 60, 2044-2055. |
| 41 | Mercer, K.L. et al. (2007) Stress and domestication traits increase the relative fitness of crop–wild hybrids in sunflower. *Ecology Letters*, 10, 383-393. |
| 42 | Nam, K.-H. et al. (2019) Gene flow from transgenic PPO-inhibiting herbicide-resistant rice to weedy rice, and agronomic performance by their hybrids. *Journal of Plant Biology*, 62, 286-296. |
| 43 | Schmidt, J.J. et al. (2018) Growth, Fitness, and Overwinter Survival of a Shattercane (*Sorghum bicolor* ssp *drummondii*) × Grain Sorghum (*Sorghum bicolor* ssp *bicolor*) F2 Population. *Weed Science*, 66, 634-641. |
| 44 | Tang, T. et al. (2018) Transgene introgression from *Brassica napus* to different varieties of *Brassica juncea*. *Plant Breeding*, 137, 171-180. |
| 45 | Vacher, C. et al. (2004) Impact of ecological factors on the initial invasion of Bt transgenes into wild populations of birdseed rape (*Brassica rapa*). *Theoretical and Applied Genetics*, 109, 806-814. |
| 46 | Zhang, J. et al. (2018) Feral rice from introgression of weedy rice genes into transgenic herbicide-resistant hybrid-rice progeny. *Journal of Experimental Botany*, 69, 3855-3865. |
| 47 | Zhang, L. et al. (2023) Fitness changes in wild soybean caused by gene flow from genetically modified soybean. *BMC Plant Biology*, 23, 424. |
| 48 | Shengnan, L. et al. (2016) Fitness of hybrids between two types of transgenic rice and six japonica and indica weed rice accessions. *Crop Science*, 56, 2751-2765. |
| 49 | Shivrain, V.K. et al. (2009) Gene flow from weedy red rice (*Oryza sativa* L.) to cultivated rice and fitness of hybrids. *Pest Management Science*, 65, 1124-1129. |
| 50 | Wei, W. and Darmency, H. (2008) Gene flow hampered by low seed size of hybrids between oilseed rape and five wild relatives. *Seed Science Research*, 18, 115-123. |
| 51 | YE Ping-yang et al. (2008) Seed Germination Test for Hybrids between Transgenic Cultivated Rice (*Oryza sativa*) and Annual Common Wild Rice (*Oryza nivara*). *Journal of Fudan University (Natural Science)*, 47, 329-334. |
| 52 | Yook, M.-J. et al. (2021) Environmental risk assessment of glufosinate-resistant soybean by pollen-mediated gene flow under field conditions in the region of the genetic origin. *Science of the Total Environment*, 762. |
| 53 | Yoshimura, Y. et al. (2006) Transgenic oilseed rape along transportation routes and port of Vancouver in western Canada. *Environmental Biosafety Research*, 5, 67-75. |
| 54 | Ziska, L.H. et al. (2012) Recent and projected increases in atmospheric CO2 concentration can enhance gene flow between wild and genetically altered rice (*Oryza sativa*). *PLoS ONE*, 7, e37522. |

**Supplementary Table 20**

**List of publications** **included in the** **dataset of this study.**

| No. | Publications |
| --- | --- |
| 1 | Tian, D. et al. (2003) Fitness costs of R-gene-mediated resistance in *Arabidopsis thaliana*. *Nature*, 423, 74-77. |
| 2 | Liu, L. et al. (2022) Fitness and Ecological Risk of Hybrid Progenies of Wild and Herbicide-Tolerant Soybeans With EPSPS Gene. *Frontiers in Plant Science*, 13. |
| 3 | Guan, Z.-J. et al. (2015) Performance of hybrid progeny formed between genetically modified herbicide-tolerant soybean and its wild ancestor. *AOB Plants*, 7, plv121. |
| 4 | He, Y. et al. (2015) Involvement of 14-3-3 protein GRF9 in root growth and response under polyethylene glycol-induced water stress. *Journal of Experimental Botany*, 66, 2271-2281. |
| 5 | Campo, S. et al. (2012) Expression of the maize ZmGF14-6 gene in rice confers tolerance to drought stress while enhancing susceptibility to pathogen infection. *Journal of Experimental Botany*, 63, 983-999. |
| 6 | Zhou, H. et al. (2014) Inhibition of the Arabidopsis Salt Overly Sensitive Pathway by 14-3-3 Proteins. |
| 7 | Xu, W. et al. (2013) The Tomato 14-3-3 Protein TFT4 Modulates H+ Efflux, Basipetal Auxin Transport, and the PKS5-J3 Pathway in the Root Growth Response to Alkaline Stress. |
| 8 | Abe, H. et al. (2003) Arabidopsis AtMYC2 (bHLH) and AtMYB2 (MYB) Function as Transcriptional Activators in Abscisic Acid Signaling. *The Plant Cell*, 15, 63-78. |
| 9 | Qiu, Y. and Yu, D. (2009) Over-expression of the stress-induced OsWRKY45 enhances disease resistance and drought tolerance in *Arabidopsis*. *Environmental and Experimental Botany*, 65, 35-47. |
| 10 | Rizhsky, L. et al. (2004) The Zinc Finger Protein Zat12 Is Required for Cytosolic Ascorbate Peroxidase 1 Expression during Oxidative Stress in *Arabidopsis*. *Journal of Biological Chemistry*, 279, 11736-11743. |
| 11 | Ciftci-Yilmaz, S. et al. (2007) The EAR-motif of the Cys2/His2-type Zinc Finger Protein Zat7 Plays a Key Role in the Defense Response of Arabidopsis to Salinity Stress. *Journal of Biological Chemistry*, 282, 9260-9268. |
| 12 | Chen, Y.-F. et al. (2009) The WRKY6 Transcription Factor Modulates PHOSPHATE1 Expression in Response to Low Pi Stress in *Arabidopsis*. *The Plant Cell*, 21, 3554-3566. |
| 13 | Al-Ahmad, H. et al. (2005) Poor competitive fitness of transgenically mitigated tobacco in competition with the wild type in a replacement series. *Planta*, 222, 372-385. |
| 14 | Li, S. et al. (2011) *Arabidopsis thaliana* WRKY25, WRKY26, and WRKY33 coordinate induction of plant thermotolerance. *Planta*, 233, 1237-1252. |
| 15 | Chen, L. et al. (2010) Wounding-Induced WRKY8 Is Involved in Basal Defense in *Arabidopsis*. *Molecular Plant-Microbe Interactions*, 23, 558-565. |
| 16 | Zou, C. et al. (2010) Male gametophyte-specific WRKY34 transcription factor mediates cold sensitivity of mature pollen in *Arabidopsis*. *Journal of Experimental Botany*, 61, 3901-3914. |
| 17 | Wu, X. et al. (2009) Enhanced heat and drought tolerance in transgenic rice seedlings overexpressing OsWRKY11 under the control of HSP101 promoter. *Plant Cell Reports*, 28, 21-30. |
| 18 | Song, Y. et al. (2009) Overexpression of the stress-induced OsWRKY08 improves osmotic stress tolerance in *Arabidopsis*. *Science Bulletin*, 54, 4671-4678. |
| 19 | Chen, H. et al. (2010) Roles of *Arabidopsis* WRKY18, WRKY40 and WRKY60 transcription factors in plant responses to abscisic acid and abiotic stress. *BMC Plant Biology*, 10, 281. |
| 20 | Suzuki, N. et al. (2005) Enhanced Tolerance to Environmental Stress in Transgenic Plants Expressing the Transcriptional Coactivator Multiprotein Bridging Factor 1c. *Plant Physiology*, 139, 1313-1322. |
| 21 | Li, S. et al. (2010) Functional Characterization of *Arabidopsis thaliana* WRKY39 in Heat Stress. *Molecules and Cells*, 29, 475-484. |
| 22 | Wang, H. et al. (2007) Overexpression of rice WRKY89 enhances ultraviolet B tolerance and disease resistance in rice plants. *Plant Molecular Biology*, 65, 799-815. |
| 23 | Robatzek, S. and Somssich, I.E. (2002) Targets of AtWRKY6 regulation during plant senescence and pathogen defense. *Genes & Development*, 16, 1139-1149. |
| 24 | Wang, Z. et al. (2009) A WRKY transcription factor participates in dehydration tolerance in *Boea hygrometrica* by binding to the W-box elements of the galactinol synthase (BhGolS1) promoter. *Planta*, 230, 1155-1166. |
| 25 | Snow, A.A. et al. (2003) A Bt TRANSGENE REDUCES HERBIVORY AND ENHANCES FECUNDITY IN WILD SUNFLOWERS. *Ecological Applications*, 13. |
| 26 | Malik, M.K. et al. (2002) Modified expression of a carrot small heat shock protein gene, Hsp17.7, results in increased or decreased thermotolerance. *The Plant Journal*, 20, 89-99. |
| 27 | McKersie, B.D. et al. (1999) Winter Survival of Transgenic Alfalfa Overexpressing Superoxide Dismutase. |
| 28 | McKersie, B.D. (2000) et al. Iron-Superoxide Dismutase Expression in Transgenic Alfalfa Increases Winter Survival without a Detectable Increase in Photosynthetic Oxidative Stress Tolerance. |
| 29 | Roxas, V.P. et al. (1997) Overexpression of glutathione S-transferase/glutathioneperoxidase enhances the growth of transgenic tobacco seedlings during stress. *Nature Biotechnology*, 15, 988-991. |
| 30 | Kishitani, S. et al. (2000) Compatibility of glycinebetaine in rice plants: evaluation using transgenic rice plants with a gene for peroxisomal betaine aldehyde dehydrogenase from barley. *Plant, Cell & Environment*, 23, 107-114. |
| 31 | Alia et al. (1998) Transformation with a gene for choline oxidase enhances the cold tolerance of *Arabidopsis* during germination and early growth. *Plant, Cell & Environment*, 21, 232-239. |
| 32 | Sakamoto, A. et al. (2000) Transformation of *Arabidopsis* with the codA gene for choline oxidase enhances freezing tolerance of plants. *The Plant Journal*, 22, 449-453. |
| 33 | Shimada, T. et al. (2000) Modification of Fatty Acid Composition in Rice Plants by Transformation with a Tobacco Microsomal ω-3 Fatty Acid Desaturase Gene (NtFAD3). *Plant Biotechnology*, 17, 43-48. |
| 34 | Kasuga et al. (1999) Improving plant drought, salt, and freezing tolerance by gene transfer of a single stress-inducible transcription factor. *Nature Biotechnology*, 17, 287-291. |
| 35 | Ding, A. et al. (2022) A chimeric AtERF4 repressor modulates pleiotropic aspects of plant growth and abiotic stress tolerance in transgenic *Arabidopsis*. *Plant Growth Regulation*, 96, 255-267. |
| 36 | Chen, J. et al. (2018) A cold-induced pectin methyl-esterase inhibitor gene contributes negatively to freezing tolerance but positively to salt tolerance in *Arabidopsis*. *Journal of Plant Physiology*, 222, 67-78. |
| 37 | Welz, H.G. et al. (1995) Two unnecessary powdery mildew resistance genes in a synthetic rye population are neutral on fitness. *Euphytica*, 81, 163-170. |
| 38 | Tunc-Ozdemir, M. et al. (2013) A Cyclic Nucleotide-Gated Channel (CNGC16) in Pollen Is Critical for Stress Tolerance in Pollen Reproductive Development. *Plant Physiology*, 161, 1010-1020. |
| 39 | Lee, G. et al. (2024) A large-effect fitness trade-off across environments is explained by a single mutation affecting cold acclimation. *Proceedings of the National Academy of Sciences*, 121, e2317461121. |
| 40 | Kim, S.-G. et al. (2008) A membrane-bound NAC transcription factor NTL8 regulates gibberellic acid-mediated salt signaling in *Arabidopsis* seed germination. *The Plant Journal*, 55, 77-88. |
| 41 | Li, C. et al. (2014) An ABA-responsive DRE-binding protein gene from *Setaria italica*, SiARDP, the target gene of SiAREB, plays a critical role under drought stress. *Journal of Experimental Botany*, 65, 5415-5427. |
| 42 | Shen, Y.-G. et al. (2003) Characterization of a DRE-binding transcription factor from a halophyte *Atriplex hortensis*. *Theoretical and Applied Genetics*, 107, 155-161. |
| 43 | Tsutsui, T. et al. (2009) DEAR1, a transcriptional repressor of DREB protein that mediates plant defense and freezing stress responses in *Arabidopsis*. *Journal of Plant Research*, 122, 633-643. |
| 44 | Fukao, T. et al. (2006) A Variable Cluster of Ethylene Response Factor–Like Genes Regulates Metabolic and Developmental Acclimation Responses to Submergence in Rice. *The Plant Cell*, 18, 2021-2034. |
| 45 | Burke, J.J. and Chen, J. (2015) Enhancement of Reproductive Heat Tolerance in Plants. *PLoS ONE*, 10, e0122933. |
| 46 | Queitsch, C. et al. (2000) Heat Shock Protein 101 Plays a Crucial Role in Thermotolerance in *Arabidopsis*. *The Plant Cell*, 12, 479-492. |
| 47 | Chandran, D. et al. (2014) Atypical E2F Transcriptional Repressor DEL1 Acts at the Intersection of Plant Growth and Immunity by Controlling the Hormone Salicylic Acid. *Cell Host & Microbe*, 15, 506-513. |
| 48 | De Vleesschauwer, D. et al. (2018) Target of rapamycin signaling orchestrates growth–defense trade-offs in plants. *New Phytologist*, 217, 305-319. |
| 49 | Campos, M.L. et al. (2016) Rewiring of jasmonate and phytochrome B signalling uncouples plant growth-defense tradeoffs. *Nature Communications*, 7, 12570. |
| 50 | Kubo, A. et al. (2013) Characterization of hybrids between wild and genetically modified glyphosate-tolerant soybeans. *Plant Biotechnology*, 30, 335-U58. |
| 51 | Artlip, T. et al. (2014) Field evaluation of apple overexpressing a peach CBF gene confirms its effect on cold hardiness, dormancy, and growth. *Environmental and Experimental Botany*, 106, 79-86. |
| 52 | Hsieh, T.-H. et al. (2002) Heterology Expression of the *Arabidopsis* C-Repeat/Dehydration Response Element Binding Factor 1 Gene Confers Elevated Tolerance to Chilling and Oxidative Stresses in Transgenic Tomato. *Plant Physiology*, 129, 1086-1094. |
| 53 | Pino, M.-T. et al. (2008) Ectopic AtCBF1 over-expression enhances freezing tolerance and induces cold acclimation-associated physiological modifications in potato. *Plant, Cell & Environment*, 31, 393-406. |
| 54 | Kitashiba, H. et al. (2004) Expression of a sweet cherry DREB1/CBF ortholog in *Arabidopsis* confers salt and freezing tolerance. *Journal of Plant Physiology*, 161, 1171-1176. |
| 55 | Liu, Q. et al. (1998) Two Transcription Factors, DREB1 and DREB2, with an EREBP/AP2 DNA Binding Domain Separate Two Cellular Signal Transduction Pathways in Drought- and Low-Temperature-Responsive Gene Expression, Respectively, in *Arabidopsis*. *The Plant Cell*, 10, 1391-1406. |
| 56 | Welling, A. and Palva, E.T. (2008) Involvement of CBF Transcription Factors in Winter Hardiness in Birch. *Plant Physiology*, 147, 1199-1211. |
| 57 | Benedict, C. et al. (2006) The CBF1-dependent low temperature signalling pathway, regulon and increase in freeze tolerance are conserved in *Populus* spp. *Plant, Cell & Environment*, 29, 1259-1272. |
| 58 | Tillett, R.L. et al. (2012) The *Vitis vinifera* C-repeat binding protein 4 (VvCBF4) transcriptional factor enhances freezing tolerance in wine grape. *Plant Biotechnology Journal*, 10, 105-124. |
| 59 | Navarro, M. et al. (2010) Two EguCBF1 genes overexpressed in *Eucalyptus* display a different impact on stress tolerance and plant development. *Plant Biotechnology Journal*, 9, 50-63. |
| 60 | Wisniewski, M. et al. (2011) Ectopic expression of a novel peach (*Prunus persica*) CBF transcription factor in apple (*Malus × domestica*) results in short-day induced dormancy and increased cold hardiness. *Planta*, 233, 971-983. |
| 61 | Gilmour, S.J. et al. (2000) Overexpression of the *Arabidopsis* CBF3 Transcriptional Activator Mimics Multiple Biochemical Changes Associated with Cold Acclimation. *Plant Physiology*, 124, 1854-1865. |
| 62 | Su, C.-F. et al. (2010) A Novel MYBS3-Dependent Pathway Confers Cold Tolerance in Rice. *Plant Physiology*, 153, 145-158. |
| 63 | Mukhopadhyay, A. et al. (2004) Overexpression of a zinc-finger protein gene from rice confers tolerance to cold, dehydration, and salt stress in transgenic tobacco. *Proceedings of the National Academy of Sciences*, 101, 6309-6314. |
| 64 | Vannini, C. et al. (2004) Overexpression of the rice Osmyb4 gene increases chilling and freezing tolerance of *Arabidopsis thaliana* plants. *The Plant Journal*, 37, 115-127. |
| 65 | Pasquali, G. et al. (2008) Osmyb4 expression improves adaptive responses to drought and cold stress in transgenic apples. *Plant Cell Reports*, 27, 1677-1686. |
| 66 | Dai, X. et al. (2007) Overexpression of an R1R2R3 MYB Gene, OsMYB3R-2, Increases Tolerance to Freezing, Drought, and Salt Stress in Transgenic *Arabidopsis*. *Plant Physiology*, 143, 1739-1751. |
| 67 | Ma, Q. et al. (2009) Enhanced Tolerance to Chilling Stress in OsMYB3R-2 Transgenic Rice Is Mediated by Alteration in Cell Cycle and Ectopic Expression of Stress Genes. *Plant Physiology*, 150, 244-256. |
| 68 | Sato, H. et al. (2016) The *Arabidopsis* transcriptional regulator DPB3-1 enhances heat stress tolerance without growth retardation in rice. *Plant Biotechnology Journal*, 14, 1756-1767. |
| 69 | Whaley, C.M. et al. (2007) A new mutation in plant ALS confers resistance to five classes of ALS-inhibiting herbicides. *Weed Science*, 55, 83-90. |
| 70 | Wang, W. et al. (2014) A novel 5-enolpyruvoylshikimate-3-phosphate (EPSP) synthase transgene for glyphosate resistance stimulates growth and fecundity in weedy rice (*Oryza sativa*) without herbicide. *New Phytologist*, 202, 679-688. |
| 71 | Yang, X. et al. (2011) Transgenes for insect resistance reduce herbivory and enhance fecundity in advanced generations of crop–weed hybrids of rice. *Evolutionary Applications*, 4, 672-684. |
| 72 | Laughlin, K.D. et al. (2009) Risk assessment of genetically engineered crops: fitness effects of virus-resistance transgenes in wild *Cucurbita pepo*. *Ecological Applications*, 19, 1091-1101. |
| 73 | Spencer, L.J. and Snow, A.A. (2001) Fecundity of transgenic wild-crop hybrids of *Cucurbita pepo* (Cucurbitaceae): implications for crop-to-wild gene flow. *Heredity*, 86, 694-702. |
| 74 | Zhou, L.-J. et al. (2021) A novel transcription factor CmMYB012 inhibits flavone and anthocyanin biosynthesis in response to high temperatures in chrysanthemum. *Horticulture Research*, 8. |
| 75 | Huang, M.-X. et al. (2008) A ribosome-inactivating protein (curcin 2) induced from *Jatropha curcas* can reduce viral and fungal infection in transgenic tobacco. *Plant Growth Regulation*, 54, 115-123. |
| 76 | Lodge, J.K. et al. (1993) Broad-spectrum virus resistance in transgenic plants expressing pokeweed antiviral protein. *Proceedings of the National Academy of Sciences*, 90, 7089-7093. |
| 77 | Cai, M. et al. (2007) A rice promoter containing both novel positive and negative cis-elements for regulation of green tissue-specific gene expression in transgenic plants. *Plant Biotechnology Journal*, 5. |
| 78 | Pellegrineschi, A. et al. (2004) Stress-induced expression in wheat of the *Arabidopsis thaliana* DREB1A gene delays water stress symptoms under greenhouse conditions. *Genome*, 47, 493-500. |
| 79 | Schmitt, J. et al. (1995) A test of the adaptive plasticity hypothesis using transgenic and mutant plants disabled in phytochrome-mediated elongation responses to neighbors. *The American Naturalist*, 146, 937-953. |
| 80 | Heidel, A.J. et al. (2004) Fitness Costs of Mutations Affecting the Systemic Acquired Resistance Pathway in *Arabidopsis thaliana*. *Genetics*, 168, 2197-2206. |
| 81 | Korves, T. and Bergelson, J. (2004) A Novel Cost of R Gene Resistance in the Presence of Disease. *The American Naturalist*, 163, 489-504. |
| 82 | McCloskey, W.B. and Holt, J.S. (1990) Triazine Resistance in *Senecio vulgaris* Parental and Nearly Isonuclear Backcrossed Biotypes Is Correlated with Reduced Productivity. *Plant Physiology*, 92, 954-962. |
| 83 | Bergelson, J. et al. (1997) Costs of resistance: a test using transgenic *Arabidopsis thaliana*. *Proceedings of the Royal Society of London. Series B: Biological Sciences*, 263, 1659-1663. |
| 84 | Jackson, M.W. et al. (2004) Costs and benefits of cold tolerance in transgenic *Arabidopsis thaliana*. *Molecular Ecology*, 13, 3609-3615. |
| 85 | Boulter, D. et al. (1990) Additive protective effects of different plant-derived insect resistance genes in transgenic tobacco plants. *Crop Protection*, 9, 351-354. |
| 86 | Chen, J. et al. (2024) Affecting of Glyphosate Tolerance and Metabolite Content in Transgenic *Arabidopsis thaliana* Overexpressing EPSPS Gene from *Eleusine indica*. *Plants*, 14, 78. |
| 87 | Ouyang, C. et al. (2021) The Naturally Evolved EPSPS From Goosegrass Confers High Glyphosate Resistance to Rice. *Frontiers in Plant Science*, 12. |
| 88 | Bettini, P.P. et al. (2016) *Agrobacterium rhizogenes* rolA gene promotes tolerance to *Fusarium oxysporum* f. sp. *lycopersici* in transgenic tomato plants (*Solanum lycopersicum* L.). *Journal of Plant Biochemistry and Biotechnology*, 25, 225-233. |
| 89 | Bettini, P. et al. (2003) Pleiotropic effect of the insertion of the *Agrobacterium rhizogenes* rolD gene in tomato (*Lycopersicon esculentum* Mill.). *Theoretical and Applied Genetics*, 107, 831-836. |
| 90 | Arshad, W. et al. (2014) *Agrobacterium*-Mediated Transformation of Tomato with rolB Gene Results in Enhancement of Fruit Quality and Foliar Resistance against Fungal Pathogens. *PLoS ONE*, 9, e96979. |
| 91 | Karthik, K. et al. (2020) Transgenic Cotton (*Gossypium hirsutum* L.) to Combat Weed Vagaries: Utility of an Apical Meristem-Targeted in planta Transformation Strategy to Introgress a Modified CP4-EPSPS Gene for Glyphosate Tolerance. *Frontiers in Plant Science*, 11. |
| 92 | Awan, M.F. et al. (2015) Transformation of Insect and Herbicide Resistance Genes in Cotton (*Gossypium hirsutum* L.). *Journal of Agricultural Science and Technology*, 17, 287-298. |
| 93 | Chandrasekhar, K. et al. (2014) Development of Transgenic Rice Harbouring Mutated Rice 5-Enolpyruvylshikimate 3-Phosphate Synthase (Os-mEPSPS) and *Allium sativum* Leaf Agglutinin (ASAL) Genes Conferring Tolerance to Herbicides and Sap-Sucking Insects. *Plant Molecular Biology Reporter*, 32, 1146-1157. |
| 94 | Te, Z. et al. (2011) Development of transgenic glyphosate-resistant rice with g6 gene encoding 5-enolpyruvylshikimate-3-phosphate synthase. *Agricultural Sciences in China*, 10, 1309-1316. |
| 95 | Liang, C. et al. (2017) Co-expression of GR79 EPSPS and GAT yields herbicide-resistant cotton with low glyphosate residues. *Plant Biotechnology Journal*, 15, 1622-1629. |
| 96 | Zhang, X.B. et al. (2017) Development of glyphosate-tolerant transgenic cotton plants harboring the G2-aroA gene. *Journal of Integrative Agriculture*, 16, 551-558. |
| 97 | Dun, B. et al. (2014) Development of highly glyphosate-tolerant tobacco by coexpression of glyphosate acetyltransferase gat and EPSPS G2-aroA genes. *The Crop Journal*, 2, 164-169. |
| 98 | Meilan, R. et al. (2002) The CP4 transgene provides high levels of tolerance to Roundup herbicide in field-grown hybrid poplars. *Canadian Journal of Forest Research*, 32, 967-976. |
| 99 | Fartyal, D. et al. (2018) Co-expression of P173S Mutant Rice EPSPS and igrA Genes Results in Higher Glyphosate Tolerance in Transgenic Rice. *Frontiers in Plant Science*, 9, 144. |
| 100 | Saha, P. et al. (2006) A novel approach for developing resistance in rice against phloem limited viruses by antagonizing the phloem feeding hemipteran vectors. *Plant Molecular Biology*, 62, 735-752. |
| 101 | Yarasi, B. et al. (2008) Transgenic rice expressing *Allium sativum* leaf agglutinin (ASAL) exhibits high-level resistance against major sap-sucking pests. *BMC Plant Biology*, 8, 102. |
| 102 | Vajhala, C.S.K. et al. (2013) Development of Transgenic Cotton Lines Expressing *Allium sativum* Agglutinin (ASAL) for Enhanced Resistance against Major Sap-Sucking Pests. *PLoS ONE*, 8, e72542. |
| 103 | Bharathi, Y. et al. (2011) Pyramided rice lines harbouring *Allium sativum* (asal) and *Galanthus nivalis* (gna) lectin genes impart enhanced resistance against major sap-sucking pests. *Journal of Biotechnology*, 152, 63-71. |
| 104 | Chen, Y.-C.S. et al. (2006) Expression of CP4 EPSPS in microspores and tapetum cells of cotton (*Gossypium hirsutum*) is critical for male reproductive development in response to late-stage glyphosate applications. *Plant Biotechnology Journal*, 4, 477-487. |
| 105 | Raj, S.K. et al. (2005) *Agrobacterium*-mediated tomato transformation and regeneration of transgenic lines expressing *Tomato leaf curl virus* coat protein gene for resistance against TLCV infection. *Current Science*, 89, 1674-1679. |
| 106 | Hensel, G. et al. (2017) *Agrobacterium*-Mediated Transformation of Wheat Using Immature Embryos. *Methods in Molecular Biology*, 1679, 129-139. |
| 107 | Song, X. et al. (2011) Agronomic performance of F1, F2 and F3 hybrids between weedy rice and transgenic glufosinate-resistant rice. *Pest Management Science*, 67, 921-931. |
| 108 | Oard, J. et al. (2000) Field evaluation of seed production, shattering, and dormancy in hybrid populations of transgenic rice (*Oryza sativa*) and the weed, red rice (*Oryza sativa*). *Plant Science*, 157, 13-22. |
| 109 | Zhang, N. et al. (2003) Out-crossing frequency and genetic analysis of hybrids between transgenic glufosinate herbicide-resistant rice and the weed, red rice. *Euphytica*, 130, 35-45. |
| 110 | Zhang, W. et al. (2008) Genetic and agronomic analyses of red rice-Clearfield hybrids and their progeny produced from natural and controlled crosses. *Euphytica*, 164, 659-668. |
| 111 | Vercellino, R.B. et al. (2018) AHAS Trp574Leu substitution in *Raphanus sativus* L.: screening, enzyme activity and fitness cost. *Pest Management Science*, 74, 1600-1607. |
| 112 | Ortega, M.A. et al. (2024) Altering cold-regulated gene expression decouples the salicylic acid-growth trade-off in *Arabidopsis*. *The Plant Cell*, 36, 4293-4308. |
| 113 | Verberne, M.C. et al. (2000) Overproduction of salicylic acid in plants by bacterial transgenes enhances pathogen resistance. *Nature Biotechnology*, 18, 779-783. |
| 114 | Mauch, F. et al. (2001) Manipulation of salicylate content in *Arabidopsis thaliana* by the expression of an engineered bacterial salicylate synthase. *The Plant Journal*, 25, 67-77. |
| 115 | Xue, L.-J. et al. (2013) Constitutively Elevated Salicylic Acid Levels Alter Photosynthesis and Oxidative State but Not Growth in Transgenic *Populus*. *The Plant Cell*, 25, 2714-2730. |
| 116 | Ullah, C. et al. (2022) Lack of antagonism between salicylic acid and jasmonate signalling pathways in poplar. *New Phytologist*, 235, 701-717. |
| 117 | Bowling, S.A. et al. (1997) The cpr5 mutant of *Arabidopsis* expresses both NPR1-dependent and NPR1-independent resistance. *The Plant Cell*, 9, 1573-1584. |
| 118 | Clough, S.J. et al. (2000) The *Arabidopsis* dnd1 "defense, no death" gene encodes a mutated cyclic nucleotide-gated ion channel. *Proceedings of the National Academy of Sciences*, 97, 9323-9328. |
| 119 | Lee, J. et al. (2007) Salicylic acid-mediated innate immunity in *Arabidopsis* is regulated by SIZ1 SUMO E3 ligase. *The Plant Journal*, 49, 79-90. |
| 120 | Kim, S.H. et al. (2010) The *Arabidopsis* Resistance-Like Gene SNC1 Is Activated by Mutations in SRFR1 and Contributes to Resistance to the Bacterial Effector AvrRps4. *PLoS Pathogens*, 6, e1001172. |
| 121 | Xia, H. et al. (2016) Ambient insect pressure and recipient genotypes determine fecundity of transgenic crop-weed rice hybrid progeny: Implications for environmental biosafety assessment. *Evolutionary Applications*, 9, 847-856. |
| 122 | Burke, J.M. and Rieseberg, L.H. (2003) Fitness Effects of Transgenic Disease Resistance in Sunflowers. *Science*, 300, 1250. |
| 123 | Snow, A.A. et al. (1999) Costs of transgenic herbicide resistance introgressed from *Brassica napus* into weedy *B. rapa*. *Molecular Ecology*, 8, 605-615. |
| 124 | Warwick, S.I. et al. (2008) Do escaped transgenes persist in nature? The case of an herbicide resistance transgene in a weedy *Brassica rapa* population. *Molecular Ecology*, 17, 1387-1395. |
| 125 | Wuddineh, W.A. et al. (2023) Amino acid substitutions in grapevine (*Vitis vinifera*) acetolactate synthase conferring herbicide resistance. *Plant Cell, Tissue and Organ Culture (PCTOC)*, 154, 75-87. |
| 126 | Li, M. et al. (2013) ALS herbicide resistance mutations in *Raphanus raphanistrum*: evaluation of pleiotropic effects on vegetative growth and ALS activity. *Pest Management Science*, 69, 689-695. |
| 127 | Légère, A. et al. (2013) Growth Characterization of *Kochia scoparia* with Substitutions at Pro197 or Trp574 Conferring Resistance to Acetolactate Synthase–Inhibiting Herbicides. *Weed Science*, 61, 267-276. |
| 128 | Tardif, F.J. et al. (2006) A mutation in the herbicide target site acetohydroxyacid synthase produces morphological and structural alterations and reduces fitness in *Amaranthus powellii*. *New Phytologist*, 169, 251-264. |
| 129 | Tian, S. et al. (2018) Engineering herbicide-resistant watermelon variety through CRISPR/Cas9-mediated base-editing. *Plant Cell Reports*, 37, 1353-1356. |
| 130 | Yu, Q. et al. (2010) AHAS herbicide resistance endowing mutations: effect on AHAS functionality and plant growth. *Journal of Experimental Botany*, 61, 3925-3934. |
| 131 | Ashigh, J. and Tardif, F.J. (2009) An amino acid substitution at position 205 of acetohydroxyacid synthase reduces fitness under optimal light in resistant populations of *Solanum ptychanthum*. *Weed Research*, 49, 479-489. |
| 132 | Ashigh, J. and Tardif, F.J. (2011) Water and Temperature Stress Impact Fitness of Acetohydroxyacid Synthase–Inhibiting Herbicide-Resistant Populations of Eastern Black Nightshade (*Solanum ptychanthum*). *Weed Science*, 59, 341-348. |
| 133 | Thompson, C.R. et al. (1994) Growth and Competitiveness of Sulfonylurea-Resistant and -Susceptible *Kochia scoparia*. *Weed Science*, 42, 172-179. |
| 134 | Alcocer-Ruthling, M. et al. (1992) Differential Competitiveness of Sulfonylurea Resistant and Susceptible Prickly Lettuce (*Lactuca serriola*). *Weed Technology*, 6, 303-309. |
| 135 | Alcocer-Ruthling, M. et al. (1992) Seed Biology of Sulfonylurea-Resistant and -Susceptible Biotypes of Prickly Lettuce (*Lactuca serriola*). *Weed Technology*, 6, 858-864. |
| 136 | Jacobs, B.F. et al. (1988) Growth performance of triazine-resistant and -susceptible biotypes of *Solanum nigrum* over a range of temperatures. *Canadian Journal of Botany*, 66, 847-850. |
| 137 | Warwick, S.I. and Black, L. (1981) The relative competitiveness of atrazine susceptible and resistant populations of *Chenopodium album* and *C. strictum*. *Canadian Journal of Botany*, 59, 689-693. |
| 138 | Donnelly, M.J. and Hume, D.J. (1983) Photosynthetic rates and growth analysis of triazine resistant and triazine susceptible rapeseed. *Weed Research*, 38, 51. |
| 139 | Gasquez, J. et al. (1981) Comparaison de la germination et de la croissance de biotypes sensibles et résistants aux triazines chez quatre espèces de mauvaises herbes. *Weed Research*, 21, 219-225. |
| 140 | Chen, Y. et al. (2017) CRISPR/Cas9-mediated base-editing system efficiently generates gain-of-function mutations in *Arabidopsis*. *Science China Life Sciences*, 60, 520-523. |
| 141 | Gallas, N. et al. (2024) An ancient cis-element targeted by *Ralstonia solanacearum* TALE-like effectors facilitates the development of a promoter trap that could confer broad-spectrum wilt resistance. *Plant Biotechnology Journal*, 22, 602-616. |
| 142 | Liu, J. et al. (2023) An intrinsically disordered region-containing protein mitigates the drought-growth trade-off to boost yields. *Plant Physiology*, 192, 274-292. |
| 143 | Pogorelko, G. et al. (2013) *Arabidopsis* and *Brachypodium distachyon* transgenic plants expressing *Aspergillus nidulans* acetylesterases have decreased degree of polysaccharide acetylation and increased resistance to pathogens. *Plant Physiology*, 162, 9-23. |
| 144 | Molina, A. et al. (2021) *Arabidopsis* cell wall composition determines disease resistance specificity and fitness. *Proceedings of the National Academy of Sciences*, 118. |
| 145 | Yu, L.-H. et al. (2016) *Arabidopsis* EDT1/HDG11 improves drought and salt tolerance in cotton and poplar and increases cotton yield in the field. *Plant Biotechnology Journal*, 14, 72-84. |
| 146 | Yu, H. et al. (2008) Activated Expression of an *Arabidopsis* HD-START Protein Confers Drought Tolerance with Improved Root System and Reduced Stomatal Density. *The Plant Cell*, 20, 1134-1151. |
| 147 | Yu, L. et al. (2013) *Arabidopsis* Enhanced Drought Tolerance1/HOMEODOMAIN GLABROUS11 Confers Drought Tolerance in Transgenic Rice without Yield Penalty. *Plant Physiology*, 162, 1378-1391. |
| 148 | Ruan, L. et al. (2012) Expression of *Arabidopsis* HOMEODOMAIN GLABROUS 11 Enhances Tolerance to Drought Stress in Transgenic Sweet Potato Plants. *Journal of Plant Biology*, 55, 151-158. |
| 149 | Cao, Y.-J. et al. (2009) Ectopic overexpression of AtHDG11 in tall fescue resulted in enhanced tolerance to drought and salt stress. *Plant Cell Reports*, 28, 579-588. |
| 150 | He, C. et al. (2005) Expression of an *Arabidopsis* Vacuolar Sodium/Proton Antiporter Gene in Cotton Improves Photosynthetic Performance Under Salt Conditions and Increases Fiber Yield in the Field. *Plant and Cell Physiology*, 46, 1848-1854. |
| 151 | Pasapula, V. et al. (2011) Expression of an *Arabidopsis* vacuolar H+-pyrophosphatase gene (AVP1) in cotton improves drought- and salt tolerance and increases fibre yield in the field conditions. *Plant Biotechnology Journal*, 9, 88-99. |
| 152 | Lv, S. et al. (2008) Overexpression of an H+-PPase Gene from *Thellungiella halophila* in Cotton Enhances Salt Tolerance and Improves Growth and Photosynthetic Performance. *Plant and Cell Physiology*, 49, 1150-1164. |
| 153 | Qin, H. et al. (2013) Expression of the *Arabidopsis* vacuolar H+-pyrophosphatase gene AVP1 in peanut to improve drought and salt tolerance. *Plant Biotechnology Reports*, 7, 345-355. |
| 154 | Lv, S.L. et al. (2009) Overexpression of *Thellungiella halophila* H+-PPase (TsVP) in cotton enhances drought stress resistance of plants. *Planta*, 229, 899-910. |
| 155 | Zhang, H. et al. (2009) Increased glycine betaine synthesis and salinity tolerance in AhCMO transgenic cotton lines. *Molecular Breeding*, 23, 289-298. |
| 156 | Kuppu, S. et al. (2013) Water-Deficit Inducible Expression of a Cytokinin Biosynthetic Gene IPT Improves Drought Tolerance in Cotton. *PLoS ONE*, 8, e64190. |
| 157 | Liu, G. et al. (2014) Overexpression of Rice NAC Gene SNAC1 Improves Drought and Salt Tolerance by Enhancing Root Development and Reducing Transpiration Rate in Transgenic Cotton. *PLoS ONE*, 9, e86895. |
| 158 | Mao, X. et al. (2012) TaNAC2, a NAC-type wheat transcription factor conferring enhanced multiple abiotic stress tolerances in *Arabidopsis*. *Journal of Experimental Botany*, 63, 2933-2946. |
| 159 | Tran, L.-S.P. et al. (2004) Isolation and Functional Analysis of *Arabidopsis* Stress-Inducible NAC Transcription Factors That Bind to a Drought-Responsive cis-Element in the early responsive to dehydration stress 1 Promoter. *The Plant Cell*, 16, 2481-2498. |
| 160 | Hu, H. et al. (2006) Overexpressing a NAM, ATAF, and CUC (NAC) transcription factor enhances drought resistance and salt tolerance in rice. *Proceedings of the National Academy of Sciences*, 103, 12987-12992. |
| 161 | Jeong, J.S. et al. (2013) OsNAC5 overexpression enlarges root diameter in rice plants leading to enhanced drought tolerance and increased grain yield in the field. *Plant Biotechnology Journal*, 11, 101-114. |
| 162 | Yokotani, N. et al. (2009) Tolerance to various environmental stresses conferred by the salt-responsive rice gene ONAC063 in transgenic *Arabidopsis*. *Planta*, 229, 1065-1075. |
| 163 | Xue, G.-P. et al. (2011) Overexpression of TaNAC69 Leads to Enhanced Transcript Levels of Stress Up-Regulated Genes and Dehydration Tolerance in Bread Wheat. *Molecular Plant*, 4, 697-712. |
| 164 | Munis, M.F.H. et al. (2010) A thaumatin-like protein gene involved in cotton fiber secondary cell wall development enhances resistance against *Verticillium dahliae* and other stresses in transgenic tobacco. *Biochemical and Biophysical Research Communications*, 393, 38-44. |
| 165 | Cowan, A.K. et al. (2005) Effects of senescence-induced alteration in cytokinin metabolism on source-sink relationships and ontogenic and stress-induced transitions in tobacco. *Planta*, 221, 801-814. |
| 166 | Rivero, R.M. et al. (2007) Delayed leaf senescence induces extreme drought tolerance in a flowering plant. *Proceedings of the National Academy of Sciences*, 104, 19631-19636. |
| 167 | Peleg, Z. et al. (2011) Cytokinin-mediated source/sink modifications improve drought tolerance and increase grain yield in rice under water-stress. *Plant Biotechnology Journal*, 9, 747-758. |
| 168 | Qin, H. et al. (2011) Regulated Expression of an Isopentenyltransferase Gene (IPT) in Peanut Significantly Improves Drought Tolerance and Increases Yield Under Field Conditions. *Plant and Cell Physiology*, 52, 1904-1914. |
| 169 | Apse, M.P. et al. (1999) Salt Tolerance Conferred by Overexpression of a Vacuolar Na+/H+ Antiport in *Arabidopsis*. *Science*, 285, 1256-1258. |
| 170 | Zhang, H.-X. et al. (2001) Engineering salt-tolerant *Brassica* plants: Characterization of yield and seed oil quality in transgenic plants with increased vacuolar sodium accumulation. *Proceedings of the National Academy of Sciences*, 98, 12832-12836. |
| 171 | Zhang, H.-X. and Blumwald, E. (2001) Transgenic salt-tolerant tomato plants accumulate salt in foliage but not in fruit. *Nature Biotechnology*, 19, 765-768. |
| 172 | Fukuda, A. et al. (2004) Function, Intracellular Localization and the Importance in Salt Tolerance of a Vacuolar Na+/H+ Antiporter from Rice. *Plant and Cell Physiology*, 45, 146-159. |
| 173 | Ohta, M. et al. (2002) Introduction of a Na+/H+ antiporter gene from *Atriplex gmelini* confers salt tolerance to rice. *FEBS Letters*, 532, 279-282. |
| 174 | Wu, C.-A. et al. (2004) The Cotton GhNHX1 Gene Encoding a Novel Putative Tonoplast Na+/H+ Antiporter Plays an Important Role in Salt Stress. *Plant and Cell Physiology*, 45, 600-607. |
| 175 | Vigouroux, Y. and Darmency, H. (2017) Assessing fitness parameters of hybrids between weed beets and transgenic sugar beets. *Plant Breeding*, 136, 969-976. |
| 176 | Darmency, H. et al. (2007) Transgene escape in sugar beet production fields: data from six years farm scale monitoring. *Environmental Biosafety Research*, 6, 197-206. |
| 177 | Raybould, A. et al. (2012) Assessing the ecological risks from the persistence and spread of feral populations of insect-resistant transgenic maize. *Transgenic Research*, 21, 655-664. |
| 178 | Toguri, T. et al. (2008) Assessment of fertility of virustolerant transgenic chrysanthemum and survey on viruses in wild chrysanthemum populations in Western Japan. *Floriculture, Ornamental and Plant Biotechnology*, 5, 490-495. |
| 179 | Ogawa, T. et al. (2005) Double-stranded RNA-specific Ribonuclease Confers Tolerance against *Chrysanthemum Stunt Viroid* and *Tomato Spotted Wilt Virus* in Transgenic Chrysanthemum Plants. *Breeding Science*, 55, 49-55. |
| 180 | Ferradini, N. et al. (2015) Assessment of heat shock protein 70 induction by heat in alfalfa varieties and constitutive overexpression in transgenic plants. *PLoS ONE*, 10, e0126051. |
| 181 | Kølster, P. et al. (1986) Near-Isogenic Barley Lines with Genes for Resistance to Powdery Mildew. *Crop Science*, 26, 903-907. |
| 182 | Kjær, B. et al. (1990) Associations between three ml-o powdery mildew resistance genes and agronomic traits in barley. *Euphytica*, 46, 185-193. |
| 183 | Sharp, G.L. et al. (2002) Field Evaluation of Transgenic and Classical Sources of *Wheat streak mosaic virus* Resistance. *Crop Science*, 42, 105-110. |
| 184 | Li, X. et al. (2016) The Systemic Acquired Resistance Regulator OsNPR1 Attenuates Growth by Repressing Auxin Signaling through Promoting IAA-Amido Synthase Expression. *Plant Physiology*, 172, 546-558. |
| 185 | Mei, C. et al. (2006) Inducible Overexpression of a Rice Allene Oxide Synthase Gene Increases the Endogenous Jasmonic Acid Level, PR Gene Expression, and Host Resistance to Fungal Infection. *Molecular Plant-Microbe Interactions*, 19, 1127-1137. |
| 186 | Liu, H. et al. (2017) NBS-LRR Protein Pik-H4 Interacts with OsBIHD1 to Balance Rice Blast Resistance and Growth by Coordinating Ethylene-Brassinosteroid Pathway. *Frontiers in Plant Science*, 8. |
| 187 | Shimono, M. et al. (2007) Rice WRKY45 Plays a Crucial Role in Benzothiadiazole-Inducible Blast Resistance. *The Plant Cell*, 19, 2064-2076. |
| 188 | Goto, S. et al. (2015) Development of disease-resistant rice by optimized expression of WRKY45. *Plant Biotechnology Journal*, 13, 753-765. |
| 189 | Tao, Z. et al. (2011) OsWRKY45 alleles play different roles in abscisic acid signalling and salt stress tolerance but similar roles in drought and cold tolerance in rice. *Journal of Experimental Botany*, 62, 4863-4874. |
| 190 | Li, W. et al. (2017) A Natural Allele of a Transcription Factor in Rice Confers Broad-Spectrum Blast Resistance. *Cell*, 170, 114-126.e15. |
| 191 | Ortelli, S. et al. (1996) Leaf Rust Resistance Gene Lr9 and Winter Wheat Yield Reduction: I. Yield and Yield Components. *Crop Science*, 36, 1590-1595. |
| 192 | Karasov, T.L. et al. (2014) The long-term maintenance of a resistance polymorphism through diffuse interactions. *Nature*, 512, 436-440. |
| 193 | MacQueen, A. et al. (2016) Genetic architecture and pleiotropy shape costs of Rps2-mediated resistance in *Arabidopsis thaliana*. *Nature Plants*, 2, 16110. |
| 194 | Wang, X. et al. (2015) Genome-Wide Association of Rice Blast Disease Resistance and Yield-Related Components of Rice. *Molecular Plant-Microbe Interactions*, 28, 1383-1392. |
| 195 | Deng, Y. et al. (2017) Epigenetic regulation of antagonistic receptors confers rice blast resistance with yield balance. *Science*, 355, 962-965. |
| 196 | Wu, Y. et al. (2016) Development of near-isogenic lines with different alleles of Piz locus and analysis of their breeding effect under Yangdao 6 background. *Molecular Breeding*, 36, 12. |
| 197 | Wu, Y. et al. (2017) Development and Evaluation of Near-Isogenic Lines with Different Blast Resistance Alleles at the Piz Locus in *japonica* Rice from the Lower Region of the Yangtze River, China. *Plant Disease*, 101, 1283-1291. |
| 198 | Hu, K. et al. (2017) Improvement of multiple agronomic traits by a disease resistance gene via cell wall reinforcement. *Nature Plants*, 3, 17009. |
| 199 | Xu, G. et al. (2017) uORF-mediated translation allows engineered plant disease resistance without fitness costs. *Nature*, 545, 491-494. |
| 200 | Berens, M.L. et al. (2019) Balancing trade-offs between biotic and abiotic stress responses through leaf age-dependent variation in stress hormone cross-talk. *Proceedings of the National Academy of Sciences*, 116, 2364-2373. |
| 201 | Bartsch, D. et al. (2001) Biosafety of hybrids between transgenic virus-resistant sugar beet and Swiss chard. *Ecological Applications*, 11, 142-147. |
| 202 | Wang, L. et al. (2016) Both overexpression and suppression of an *Oryza sativa* NB-LRR-like gene OsLSR result in autoactivation of immune response and thiamine accumulation. *Scientific Reports*, 6, 24079. |
| 203 | Abdula, S.E. et al. (2016) bruge1 transgenic rice showed improved growth performance with enhanced drought tolerance. *Breeding Science*, 66, 226-233. |
| 204 | Liu, H.-l. et al. (2007) Over-expression of OsUGE-1 altered raffinose level and tolerance to abiotic stress but not morphology in *Arabidopsis*. *Journal of Plant Physiology*, 164, 1384-1390. |
| 205 | Halfhill, M.D. et al. (2002) Bt-transgenic oilseed rape hybridization with its weedy relative, *Brassica rapa*. *Environmental Biosafety Research*, 1, 19-28. |
| 206 | Roux, F. et al. (2006) Building of an Experimental Cline With *Arabidopsis thaliana* to Estimate Herbicide Fitness Cost. *Genetics*, 173, 1023-1031. |
| 207 | Roux, F. et al. (2004) The Dominance of the Herbicide Resistance Cost in Several *Arabidopsis thaliana* Mutant Lines. *Genetics*, 166, 449-460. |
| 208 | Cai, J. et al. (2021) Glycosylation of N-hydroxy-pipecolic acid equilibrates between systemic acquired resistance response and plant growth. *Molecular Plant*, 14, 440-455. |
| 209 | Smirnova, E. et al. (2017) Jasmonic Acid Oxidase 2 Hydroxylates Jasmonic Acid and Represses Basal Defense and Resistance Responses against *Botrytis cinerea* Infection. *Molecular Plant*, 10, 1159-1173. |
| 210 | Hu, P. et al. (2013) JAV1 Controls Jasmonate-Regulated Plant Defense. *Molecular Cell*, 50, 504-515. |
| 211 | Major, I.T. et al. (2020) A Phytochrome B-Independent Pathway Restricts Growth at High Levels of Jasmonate Defense. *Plant Physiology*, 183, 733-749. |
| 212 | Zavala, J.A. and Baldwin, I.T. (2004) Fitness benefits of trypsin proteinase inhibitor expression in *Nicotiana attenuata* are greater than their costs when plants are attacked. *BMC Ecology*, 4, 11. |
| 213 | Zavala, J.A. et al. (2004) Constitutive and inducible trypsin proteinase inhibitor production incurs large fitness costs in *Nicotiana attenuata*. *Proceedings of the National Academy of Sciences*, 101, 1607-1612. |
| 214 | Watrud, L.S. et al. (2011) Changes in constructed *Brassica* communities treated with glyphosate drift. *Ecological Applications*, 21, 525-538. |
| 215 | Brütting, C. et al. (2017) Changes in cytokinins are sufficient to alter developmental patterns of defense metabolites in *Nicotiana attenuata*. *The Plant Journal*, 89, 15-30. |
| 216 | Bergelson, J. (1994) Changes in Fecundity Do Not Predict Invasiveness: A Model Study of Transgenic Plants. *Ecology*, 75, 249-252. |
| 217 | Londo, J.P. et al. (2011) Changes in fitness-associated traits due to the stacking of transgenic glyphosate resistance and insect resistance in *Brassica napus* L. *Heredity*, 107, 328-337. |
| 218 | Kidokoro, S. et al. (2014) Soybean DREB1/CBF-type transcription factors function in heat and drought as well as cold stress-responsive gene expression. *The Plant Journal*, 81, 505-518. |
| 219 | Zhang, L. et al. (2025) Changes in the Stress Response and Fitness of Hybrids Between Transgenic Soybean and Wild-Type Plants Under Heat Stress. *Plants*, 14, 622. |
| 220 | Sohn, S.-I. et al. (2022) Characteristics and Fitness Analysis through Interspecific Hybrid Progenies of Transgenic *Brassica napus* and *B. rapa* L. ssp. *International Journal of Molecular Sciences*, 23, 10512. |
| 221 | Gui, X. et al. (2015) Response difference of transgenic and conventional rice (*Oryza sativa*) to nanoparticles (γFe2O3). *Environmental Science and Pollution Research*, 22, 17716-17723. |
| 222 | Tamizhselvan, P. et al. (2024) Chloroplast Auxin Efflux Mediated by ABCB28 and ABCB29 Fine-Tunes Salt and Drought Stress Responses in *Arabidopsis*. *Plants*, 13, 7. |
| 223 | Mozgova, I. et al. (2015) Chromatin assembly factor CAF-1 represses priming of plant defence response genes. *Nature Plants*, 1, 1-8. |
| 224 | Sun, T. et al. (2023) Co-chaperoning of chlorophyll and carotenoid biosynthesis by ORANGE family proteins in plants. *Molecular Plant*, 16, 1048-1065. |
| 225 | Wu, H. et al. (2012) Co-overexpression FIT with AtbHLH38 or AtbHLH39 in *Arabidopsis*-enhanced cadmium tolerance via increased cadmium sequestration in roots and improved iron homeostasis of shoots. *Plant Physiology*, 158, 790-800. |
| 226 | Liang, Y. et al. (2018) Coexistence of *Bacillus thuringiensis* (Bt)-transgenic and conventional rice affects insect abundance and plant fitness in fields. *Pest Management Science*, 74, 1646-1653. |
| 227 | Fuchs, M. et al. (2004) Comparative fitness of a wild squash species and three generations of hybrids between wild × virus-resistant transgenic squash. *Environmental Biosafety Research*, 3, 17-28. |
| 228 | Chan, Z. et al. (2012) Comparison of salt stress resistance genes in transgenic *Arabidopsis thaliana* indicates that extent of transcriptomic change may not predict secondary phenotypic or fitness effects. *Plant Biotechnology Journal*, 10, 284-300. |
| 229 | Ouyang, D. et al. (2021) Compensation of Wild Plants Weakens the Effects of Crop-Wild Gene Flow on Wild Rice Populations. *Frontiers in Plant Science*, 12, 681008. |
| 230 | Li, L. et al. (2016) Limited ecological risk of insect-resistance transgene flow from cultivated rice to its wild ancestor based on life-cycle fitness assessment. *Science Bulletin*, 61, 1440-1450. |
| 231 | Jin, W. et al. (2009) Improved Cold-resistant Performance in Transgenic Grape (*Vitis vinifera* L.) Overexpressing Cold-inducible Transcription Factors AtDREB1b. *HortScience*, 44, 35-39. |
| 232 | Molinari, H.B.C. et al. (2004) Osmotic adjustment in transgenic citrus rootstock Carrizo citrange (*Citrus sinensis* Osb. × *Poncirus trifoliata* L. Raf.) overproducing proline. *Plant Science*, 167, 1375-1381. |
| 233 | Karakas, B. et al. (1997) Salinity and drought tolerance of mannitol-accumulating transgenic tobacco. *Plant, Cell & Environment*, 20, 609-616. |
| 234 | Romero, C. et al. (1997) Expression of the yeast trehalose-6-phosphate synthase gene in transgenic tobacco plants: pleiotropic phenotypes include drought tolerance. *Planta*, 201, 293-297. |
| 235 | Walworth, A.E. et al. (2012) Overexpression of a blueberry-derived CBF gene enhances cold tolerance in a southern highbush blueberry cultivar. *Molecular Breeding*, 30, 1313-1323. |
| 236 | Riemenschneider, D.E. and Haissig, B.E. (1991) Producing Herbicide Tolerant *Populus* Using Genetic Transformation Mediated by *Agrobacterium tumefaciens* C58: A Summary of Recent Research. *Woody Plant Biotechnology*, 247-263. |
| 237 | Donahue, R.A. et al. (1994) Growth, photosynthesis, and herbicide tolerance of genetically modified hybrid poplar. *Canadian Journal of Forest Research*, 24, 2377-2383. |
| 238 | Ullah, C. et al. (2019) Salicylic acid activates poplar defense against the biotrophic rust fungus *Melampsora larici-populina* via increased biosynthesis of catechin and proanthocyanidins. *New Phytologist*, 221, 960-975. |
| 239 | Ferreira, S.A. et al. (2002) Virus Coat Protein Transgenic Papaya Provides Practical Control of *Papaya ringspot virus* in Hawaii. *Plant Disease*, 86, 101-105. |
| 240 | Fitch, M.M.M. et al. (1992) Virus Resistant Papaya Plants Derived from Tissues Bombarded with the Coat Protein Gene of *Papaya ringspot virus*. *Nature Biotechnology*, 10, 1466-1472. |
| 241 | Lius, S. et al. (1997) Pathogen-derived resistance provides papaya with effective protection against *Papaya ringspot virus*. *Molecular Breeding*, 3, 161-168. |
| 242 | Scorza, R. et al. (2013) 'Honeysweet' - A Transgenic Plum Pox Virus Resistant Plum - From Laboratory and Experimental Field Plots, to Regulatory Approval. *Acta Horticulturae*, 57-63. |
| 243 | Liang, H. et al. (2001) Increased *Septoria musiva* resistance in transgenic hybrid poplar leaves expressing a wheat oxalate oxidase gene. *Plant Molecular Biology*, 45, 619-629. |
| 244 | Wang, G. et al. (1996) Poplar (*Populus nigra* L.) plants transformed with a *Bacillus thuringiensis* toxin gene: insecticidal activity and genomic analysis. *Transgenic Research*, 5, 289-301. |

**Supplementary Table 21**

**The description of MAT from multi‑model mean (MMM).** Under three Shared Socioeconomic Pathways (SSP1‑2.6, SSP2‑4.5, SSP5‑8.5), the MMM data was derived from the average of five CMIP6 global climate models with different climate sensitivity: EC-Earth3-Veg (low), IPSL‑CM6A‑LR (high), MPI‑ESM1‑2‑HR (low), MRI‑ESM2‑0 (low), UKESM1‑0‑LL (high), following the ISIMIP3b protocol.

| ID | SSP | Period | Describe | Min_value | Max_value | resolution |
| --- | --- | --- | --- | --- | --- | --- |
| MMM126_now | ssp126 | 2021-2040 | Near | -53.58 | 32.52 | (0.042, 0.042) |
| MMM245_now | ssp245 | 2021-2040 | Near | -53.54 | 32.58 | (0.042, 0.042) |
| MMM585_now | ssp585 | 2021-2040 | Near | -53.4 | 32.68 | (0.042, 0.042) |
| MMM126_mid | ssp126 | 2041-2060 | Mid | -53.14 | 32.94 | (0.042, 0.042) |
| MMM245_mid | ssp245 | 2041-2060 | Mid | -52.72 | 33.32 | (0.042, 0.042) |
| MMM585_mid | ssp585 | 2041-2060 | Mid | -52.24 | 33.98 | (0.042, 0.042) |
| MMM126_long | ssp126 | 2061-2080 | Long | -52.94 | 33.06 | (0.042, 0.042) |
| MMM245_long | ssp245 | 2061-2080 | Long | -52.04 | 34 | (0.042, 0.042) |
| MMM585_long | ssp585 | 2061-2080 | Long | -50.76 | 35.5 | (0.042, 0.042) |
| MMM126_mostlong | ssp126 | 2081-2100 | End | -52.84 | 32.94 | (0.042, 0.042) |
| MMM245_mostlong | ssp245 | 2081-2100 | End | -51.54 | 34.52 | (0.042, 0.042) |
| MMM585_mostlong | ssp585 | 2081-2100 | End | -48.98 | 37.38 | (0.042, 0.042) |

**Supplementary Table 22**

**Publication bias assessment for the main meta‑analysis models.** k refers to the number of effect sizes (observations) included in each subset. Pooled estimates were derived from random‑effects models fitted to study‑level aggregated data (one observation per Control_ID). For the Field subset, the result of (k₀) and Fail‑safe N should be interpreted with caution given the non‑significant pooled estimates (p = 0.234).

| Subset | k | Estimate (95% CI) | p | Egger's test (p) | Trim‑&‑fill (k₀) | Fail‑safe N |
| --- | --- | --- | --- | --- | --- | --- |
| Total | 396 | -0.058 (-0.100, -0.016) | 0.007 | 0.0062 | 0 | 703 |
| Field | 138 | 0.047 (-0.031, 0.125) | 0.234 | 0.5318 | 40 | 0 |
| Control | 262 | -0.113 (-0.160, -0.065) | < 0.001 | < 0.0001 | 0 | 1,192 |

**Supplementary Table 23**

**Publication bias assessment for fitness dimension‑specific meta‑analyses.** k refers to the number of effect sizes (observations) included in each subset. Pooled estimates were derived from random‑effects models fitted to study‑level aggregated data (one observation per Control_ID). The Demographics‑Control subset was excluded due to insufficient sample size (only 2 observations). For the Demographics dimension, the small sample sizes (Overall: 25; Field: 23) mean that the bias assessment results (Egger's test and trim‑and‑fill) should be interpreted with caution.

| Dimension | Subset | k (obs) | Estimate (95% CI) | p | Egger's test (p) | Trim‑&‑fill (k₀) | Fail‑safe N |
| --- | --- | --- | --- | --- | --- | --- | --- |
| Growth | Total | 516 | -0.090 (-0.134, -0.046) | < 0.001 | < 0.001 | 0 | 934 |
| Growth | Field | 123 | 0.013 (-0.062, 0.089) | 0.726 | 0.58 | 24 | 0 |
| Growth | Control | 393 | -0.124 (-0.177, -0.072) | < 0.001 | < 0.001 | 0 | 1,036 |
| Reproduction | Total | 715 | -0.029 (-0.080, 0.023) | 0.28 | 0.007 | 0 | 0 |
| Reproduction | Field | 360 | 0.044 (-0.034, 0.121) | 0.268 | 0.148 | 35 | 0 |
| Reproduction | Control | 355 | -0.084 (-0.152, -0.016) | 0.016 | < 0.001 | 0 | 79 |
| Demographics | Total | 25 | -0.192 (-0.796, 0.412) | 0.51 | 0.776 | 6 | 0 |
| Demographics | Field | 23 | -0.191 (-0.841, 0.458) | 0.539 | 0.77 | 5 | 0 |
| Demographics | Control | — | — | — | — | — | — |
